# NanoCaller for accurate detection of SNPs and indels in difficult-to-map regions from long-read sequencing by haplotype-aware deep neural networks

**DOI:** 10.1101/2019.12.29.890418

**Authors:** Mian Umair Ahsan, Qian Liu, Li Fang, Kai Wang

## Abstract

Long-read sequencing enables variant detection in genomic regions that are considered difficult-to-map by short-read sequencing. To fully exploit the benefits of longer reads, here we present a deep-learning method NanoCaller, which detects SNPs using long-range haplotype information, then phases long reads with called SNPs and calls indels with local realignment. Evaluation on 8 human genomes demonstrated that NanoCaller generally achieves better performance than competing approaches. We experimentally validated 41 novel variants in a widely-used benchmarking genome, which cannot be reliably detected previously. In summary, NanoCaller facilitates the discovery of novel variants in complex genomic regions from long-read sequencing.

## Introduction

Single-nucleotide polymorphisms (SNPs) and small insertions/deletions (indels) are two common types of genetic variants in human genomes. They contribute to genetic diversity and critically influence phenotypic differences, including susceptibility to human diseases. The detection (i.e. “calling”) of SNPs and indels is thus a fundamentally important problem in using the new generations of high-throughput sequencing data to study genome variations and genome functions. A number of methods have been designed to call SNPs and small indels on Illumina short-read sequencing data. Short reads are usually 100-150 bp long and have per-base error rate less than 1%. Variant calling methods on short reads, such as GATK [1] and FreeBayes [2], achieved excellent performance to detect SNPs and small indels in genomic regions marked as traditional “high-confidence regions” in various benchmarking tests[3–5]. However, since these methods were developed for short-read sequencing data with low per-base error rates and low insertion/deletion errors, they do not work well on long-read sequencing data with high error rates. Additionally, due to inherent technical limitations of short-read sequencing, the data cannot be used to call SNPs and indels in complex or repetitive genomic regions; for example, only ∼81% of genomic regions are marked as “high-confidence region” to have reliable SNP/indel calls in the Genome In A Bottle (GIAB) project, suggesting that ∼19% of the human genome is inaccessible to conventional short-read sequencing technologies to find variants reliably (please refer to the ‘Supplementary Materials 1’ on how to calculate the percentage of genomic regions.).

Oxford Nanopore [6] and Pacific Biosciences (PacBio) [7] technologies are two leading long-read sequencing platforms, which have been rapidly developed in recent years with continuously decreased costs and continuously improved read length, in comparison to Illumina short-read sequencing technologies. Long-read sequencing techniques can overcome several challenging issues that cannot be solved using short-read sequencing, such as calling long-range haplotypes, identifying variants in complex genomic regions, identifying variants in coding regions for genes with many pseudogenes, sequencing across repetitive regions, phasing of distant alleles and distinguishing highly homologous regions [8]. To date, long-read sequencing techniques have been successfully used to sequence genomes for many species to powerfully resolve various challenging biological problems such as de novo genome assembly [9–13] and SV detection [14–19]. However, the per-base accuracy of long reads is much lower with raw base calling errors of 3-15% [20] compared with short-read data (although HiFi PacBio reads and Nanopore reads generated by the latest flowcells R10.3 have lower error rates, they can be still much higher than short-read data.). The high error rate challenges widely-used variant calling methods (such as GATK [1] and FreeBayes [2]), which were previously designed for Illumina short reads and cannot handle reads with higher error rates. It is also worth noting that (1) HiFi reads after circular consensus sequencing on PacBio long-read sequencing [21] or similar methods on the Nanopore platform can potentially improve the detection of SNPs/indels by adapting existing short-read variant callers, due to its much lower per-base error rates. However, HiFi reads would substantially increase sequencing cost given the same base output, so it may be more suitable now for specific application scenarios such as capture-based sequencing or amplicon sequencing. As more and more long-read sequencing data becomes available, there is an urgent need to detect SNPs and small indels to take the most advantage of long-read data.

Several recent works aimed to design accurate SNP/indel callers on long-read sequencing data using machine learning methods, especially deep learning-based algorithms. DeepVariant [22] is among the first successful endeavor to develop a deep learning variant caller for SNPs and indels across different sequencing platforms (i.e. Illumina, PacBio and Nanopore sequencing platforms). In DeepVariant, local regions of reads aligned against a variant candidate site were transformed into an image representation, and then a deep learning framework was trained to distinguish true variants from false variants that were generated due to noisy base calls. DeepVariant achieved excellent performance on short reads as previous variant calling methods did. Later on, Clairvoyante [23] and its successor Clair [24] implemented variant calling methods using deep learning, where the summary of adjacently aligned local genomic positions of putative candidate sites were used as input of deep learning framework. The three deep learning-based methods can work well on both short-read and long-read data, but they do not incorporate haplotype structure in variant calling; these methods consider each SNP separately, while a recent testing [21] with DeepVariant has shown that a phased BAM with haplotype-sorted reads can improve variant calling accuracy because long reads likely from same haplotype benefits neural network learning from an image of read pileup. However, this testing underutilizes the rich haplotype information from long reads, especially when it is explicitly provided in a phased BAM file as input. Moreover, enough variants need to be known beforehand to phase a BAM file. Two recent works have endeavored to improve variant calling by using phasing information from long-reads sequencing data. Longshot [25] uses a pair-Hidden Markov Model (pair-HMM) for a small local window around candidate sites to call SNPs on long-read data, and then improves genotyping of called SNPs using HapCUT2 [26] based on the mostly pair of haplotypes given the current variant genotypes. However, Longshot cannot identify indels. The Oxford Nanopore Technologies company also recently released a SNP/indel caller, i.e. Medaka [27], using deep learning on long-read data. Although not published, based on its GitHub repository, Medaka first predicts SNPs from unphased long reads, and then uses WhatsHap to phase reads. Medaka finally makes SNP and indel calling for each group of phased reads. In both methods, mutual information from long-range haplotype SNPs is ignored. In summary, although several methods for variant detection on long-read sequencing data have become available, there may be room in further improving these approaches especially for difficult-to-map regions. We believe that improved SNP/indel detection on long read data will enable widespread research and clinical applications of long-read sequencing techniques.

In this study, we propose a deep learning framework, NanoCaller, which integrates long-range haplotype structure in a deep convolutional neural network to improve variant detection on long-read sequencing data. It only uses haplotype information in SNP calling (without requiring a phased BAM alignment input) and generates input features of a SNP candidate site using long-range heterozygous SNPs sites, that are fed into a deep convolutional neural network for SNP calling. Please note that those long-range heterozygous SNPs sites can be hundreds or tens of thousands of bases away and NanoCaller does not use local neighboring bases, which is substantially different from DeepVariant, Clairvoyante [23] and its successor Clair [24], as well as Longshot and Medaka where local neighboring bases of SNP sites were used. After that, NanoCaller uses these predicted SNP calls to phase alignment reads with WhatsHap for indel calling. Local multiple sequence alignment of phased reads around indel candidate sites is used to generate consensus sequence and feature inputs for a deep convolutional neural network to predict indel variant zygosity. We assess NanoCaller on 8 human genomes, HG001 (NA12878), HG002 (NA24385), HG003 (NA24149), HG004 (NA24143), HG005 (NA24631), HG006 (NA24694), HG007 (NA24695) and HX1 with both 8 Nanopore and 4 PacBio long-read data. In particular, we evaluate NanoCaller in difficult-to-map genomic regions for the Ashkenazim trio (HG002, HG003 and HG004) to investigate the unique advantages provided by long reads. Our evaluation demonstrates competitive performance of NanoCaller against existing tools, with particularly improved performance in complex genomic regions which cannot be reliably called on short-read data. NanoCaller is publicly available at https://github.com/WGLab/NanoCaller.

## Results

### Overview of NanoCaller

NanoCaller takes alignment of a long-read sequencing data aligned against a reference genome as input, and generates a VCF file for predicted SNPs and indels (Figure S6 of “Supplementary Materials 1”). For SNP calling in NanoCaller, candidate SNP sites are selected according to the specified thresholds for minimum coverage and minimum frequency of alternative alleles (a fraction of them are likely to be false positives given the relaxed thresholds for candidate identification). Long-range haplotype features for the candidate sites (Figure 1) are generated and fed to a deep convolutional network to distinguish true variants from false candidate sites. The predicted SNPs and long-reads are phased and then used in identification of indels. Indel candidate sites are selected according to specified minimum coverage and insertion/deletion frequency thresholds applied to each haploid read set. To reduce the effect of poor alignment on indel calling, NanoCaller uses a sliding window across reference genome to estimate indel frequency. Input features for indel candidate sites are generated using multiple sequencing alignment on the set of diploid reads and on each set of haploid reads (Figure 2). After that, another deep convolutional neural network is used to determine indel calls and assign variant call quality scores. Allele sequence for the indels is predicted by comparing consensus sequences against reference sequence.

**Figure 1.**
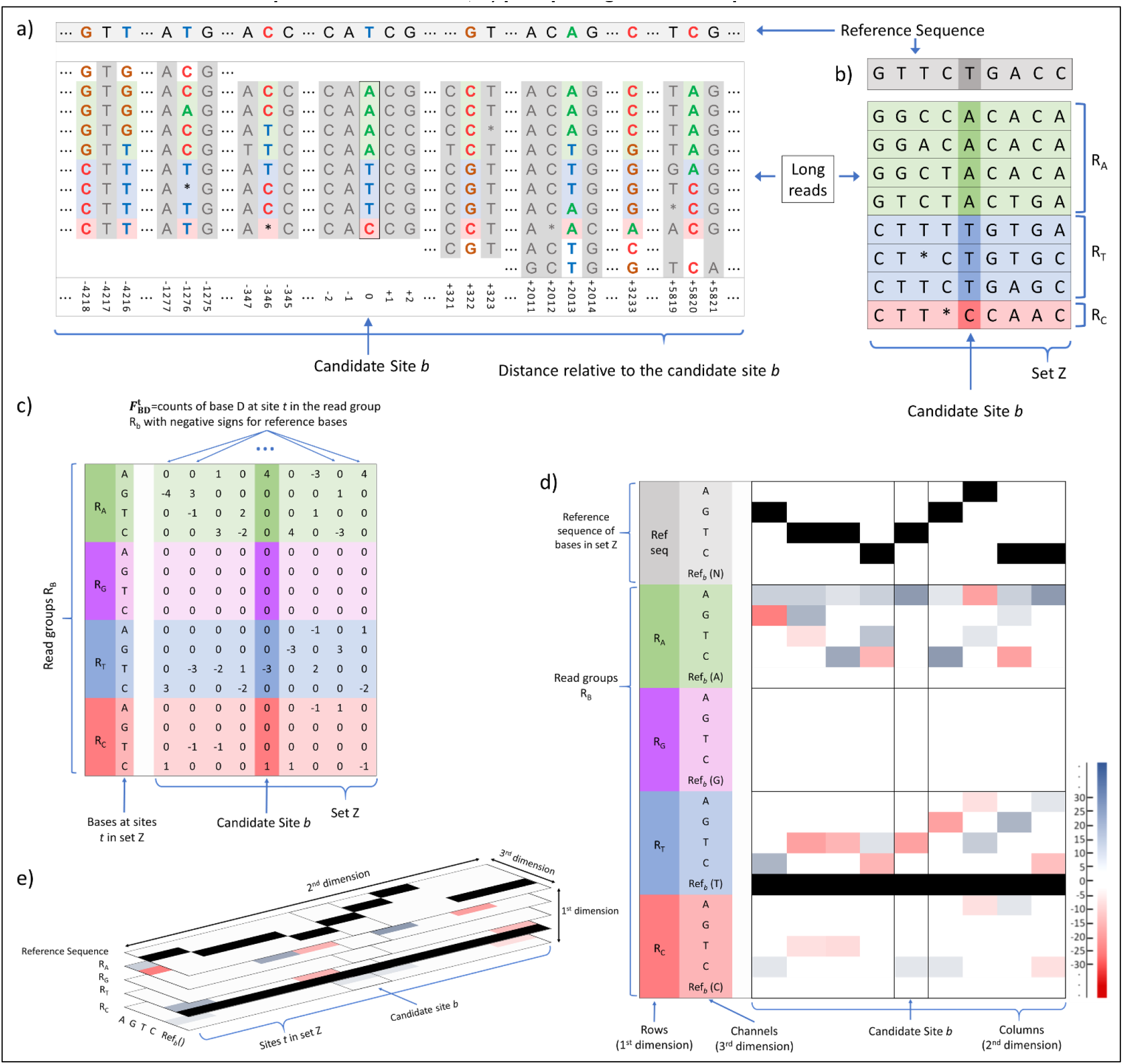
An example on how to construct image pileup for a SNP candidate site. a) reference sequence and reads pileup at candidate site *b* and at other genomic positions that share the same reads. The columns in grey are genomic positions that will not be used in input features for candidate site *b* as they do not satisfy the criteria for being highly likely heterozygous SNP sites. Only the columns with color bases will be used to generate input features for site *b* and will constitue the set Z as described in SNP pileups generation section. These neighboring likely heterozygous sites can be up to thousands of bases away from candidate site *b*; b) reference sequence and reads pileups for only the candidate site and neighboring highly likely heterozygous SNP sites; c) raw counts of bases at sites in set Z in each read group split by the nucleotide types at site *b*. These raw counts are multiplied with negative signs for reference bases; d) flattened pileup image with fifth channel after reference sequence row is added; e) pileup image used as input for NanoCaller.

**Figure 2.**
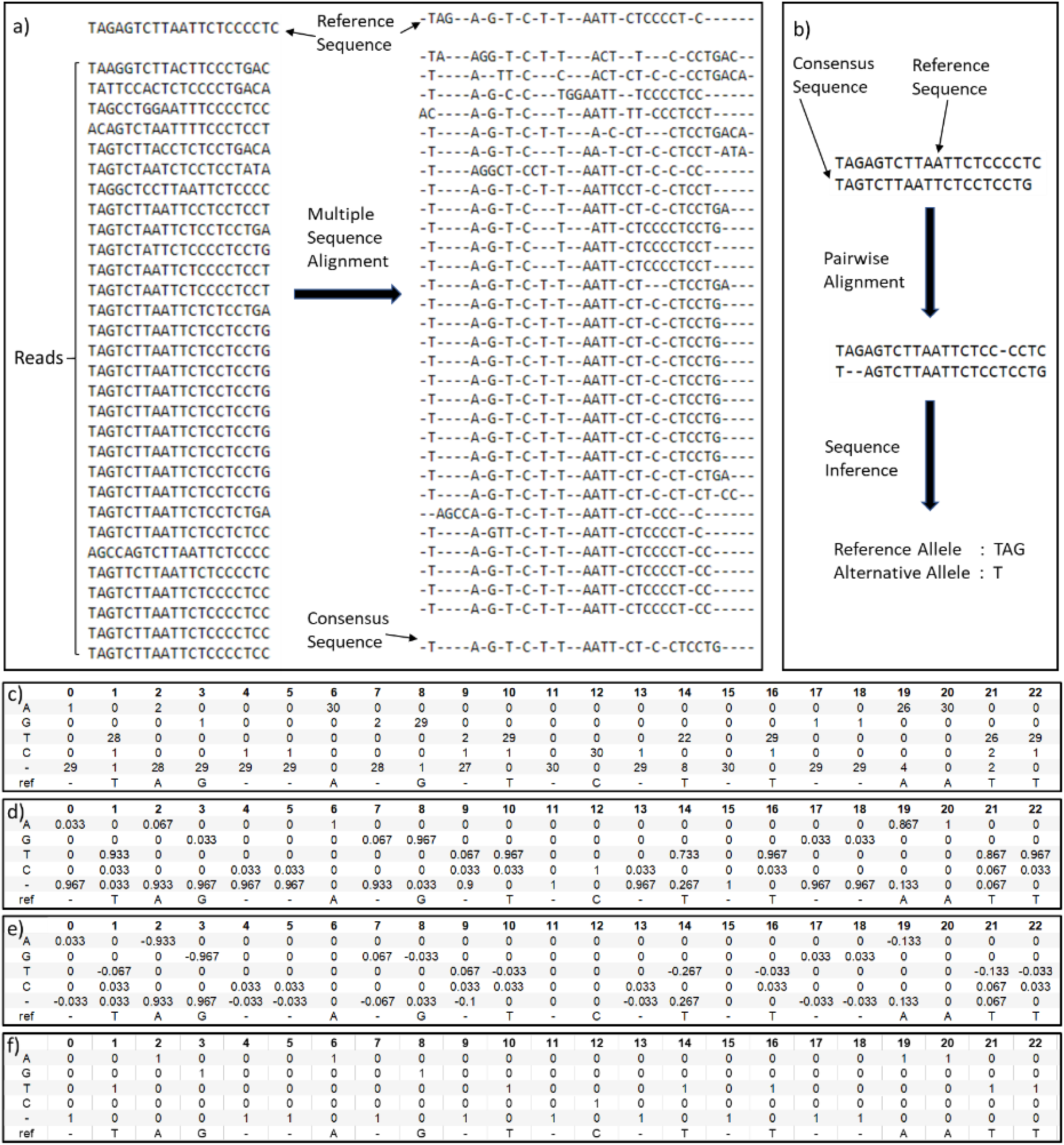
An example on how to construct image pileup for an indel site. a) reference sequence and reads pileup at the candidate site before and after multiple sequence alignment, and the consensus sequence after realignment; b) reference sequence and consensus sequence at the candidate site before and after pairwise alignment, and the inferred sequence; c) shows raw count of each symbol at each column of multiple sequence alignment pileup; d) matrix M, showing frequency of each symbol at each column of multiple sequence alignment pileup; e) first channel of input image, matrix M minus Q (one-hot encoding of realigned reference sequence); f) matrix Q, one-hot encoding of realigned reference sequence which forms the second channel of input image.

The performance of NanoCaller is evaluated on both Oxford Nanopore and PacBio reads, and compared with performances of Medaka (v0.10.0), Clair (v2.0.1), Longshot (v0.4.1), DeepVariant (v.1.0.0) and WhatsHap (v1.0) with their default parameters for each type of sequencing technology. By default, evaluation is on benchmark variants in high-confidence intervals of chromosomes 1-22 of the GRCh38 reference genome, unless stated otherwise. RTG tools (the commands for *vcfeval* submodule can be found in the ‘Supplementary Materials 1’ Pages 26-27) [28] is used to calculate various evaluation metrics, such as precision, recall and F1. For whole genome analysis, we show each variant caller’s performance using its recommended quality threshold if available, e.g. NanoCaller, Clair, Longshot and DeepVariant; for Medaka, we calculate the quality score thresholds that give highest F1 score for each genome using *vcfeval,* and use their average as the final quality score cut-off to report results. In particular, variant calling performance analysis in difficult-to-map genomic regions requires different quality score cut-offs from whole genome analysis due to highly specific error profiles of these difficult-to-map regions. Therefore, we use the average best quality score cut-off (in the same manner as the quality score cut-off is determined for Medaka) for each variant caller in each type of difficult genomic regions.

In the Results section, we present performances of five NanoCaller models: ONT-HG001 (trained on HG001 ONT reads), ONT-HG002 (trained on HG002 ONT reads), CCS-HG001 (trained on HG001 CCS reads), CCS-HG002 (trained on HG002 CCSS reads) and CLR-HG002 (trained on HG002 PacBio CLR dataset); the first four datasets have both SNP and indel deep learning models, whereas CLR-HG002 consists of only a SNP model. All NanoCaller HG001 models are trained using v3.3.2 of GIAB benchmark variant calls, whereas all NanoCaller HG002 models are trained using v4.2.1 of GIAB benchmark variant calls. Sequencing datasets used for training were aligned to the GRCh38 reference genome. For performance evaluation, the latest available GIAB benchmark variants are used for each genome, i.e. v3.3.2 for HG001 and HG005-7, and v4.2.1 for the Ashkenazim trio HG002-4). For HX1, variant calls produced by GATK on 300x Illumina reads of HX1 are used as benchmark, and high-confidence intervals for HX1 were created by removing difficult-to-map regions from chromosomes 1-22.

### Evaluation of NanoCaller on Oxford Nanopore Sequencing

#### Performance on SNP calling

We compared NanoCaller’s SNP calling performance on Oxford Nanopore sequencing reads against several existing tools. For NanoCaller, we used alternative allele frequency threshold of 0.15 for SNP candidates. For testing Clair, we used *‘1_124x ONT’* model trained on 124x coverage HG001 ONT reads using v3.3.2 GIAB benchmark variants, whereas for Medaka we used ‘*r941_min_diploid_snp_model*’ model for testing, which is trained on several bacteria and eukaryotic read datasets and variant call sets. Longshot pair-HMM model is not trained on any genome as it estimates parameters during each run. We compared the performance of those method on eight genomes: HG001-7 and HX1 under two testing strategies: cross-genome testing and cross-reference testing.

Cross-genome testing is critical to demonstrate the performance of a variant caller when used in a real-world scenario: the machine learning model of a variant caller is trained on a set of genomes and tested on other genomes. Under this testing strategy, the performance of SNP calling for NanoCaller, together with four other variant callers, Medaka, Clair, and Longshot, is shown in Table 2 and Figure 3 on Nanopore reads of eight genomes.

**Table 1.** Statistics of benchmark variants in chromosomes 1-22 of each genome aligned to the GRCh38 reference genome. Four genomes with GIAB benchmark variant calls, with v3.3.2 for HG001 and HG005-7, and v4.2.1 for HG002-4, together with the statistics within the high- confidence regions. For HX1, high-confidence regions are created by removing GIAB ‘all difficult- to-map’ regions from the GRCh38 reference genome.

|  | Whole genome |  | High confidence region |  |  |  | Non homo-polymer region |
| --- | --- | --- | --- | --- | --- | --- | --- |
| Genome | SNPs | Indels | SNPs | Indels | Total Length | % of genome | Indels |
| HG001 | 3,002,314 | 517,177 | 2,960,486 | 483,941 | 2,330,204,759 | 81.05 | 181,036 |
| HG002 | 3,459,843 | 587,987 | 3,365,115 | 525,466 | 2,542,724,465 | 88.44 | 210,352 |
| HG003 | 3,430,611 | 569,180 | 3,327,480 | 504,497 | 2,529,085,210 | 87.97 | 199,302 |
| HG004 | 3,454,689 | 576,301 | 3,346,597 | 510,516 | 2,525,035,837 | 87.83 | 200,556 |
| HG005 | 2,945,666 | 432,747 | 2,904,403 | 403,859 | 2,290,538,775 | 79.67 | 172,678 |
| HG006 | 3,030,507 | 435,520 | 2,982,278 | 405,828 | 2,348,035,455 | 81.67 | 158,063 |
| HG007 | 3,048,404 | 437,866 | 3,000,039 | 407,892 | 2,345,850,549 | 81.59 | 157,966 |
| HX1 | 3,489,068 | 697,736 | 2,788,450 | 176,587 | 2,182,959,159 | 75.93 | -- |

**Table 2.** Performances (F1 scores in %) of SNP and indel predictions by NanoCaller, Medaka, Clair, Longshot and DeepVariants on ONT and PacBio (CCS and CLR) data. These evaluation is based on v3.3.2 benchmark variants for HG001 and HG005-7 and v4.2.1 benchmark variants for the Ashkenazim trio (HG002, HG003, and HG004). Bonito and R10.3 refer to different HG002 ONT datasets.

| Prediction | Variant Caller | HG001 | HG002 | HG003 | HG004 | HG005 | HG006 | HG007 | HX1 | Bonito | R10.3 |
| --- | --- | --- | --- | --- | --- | --- | --- | --- | --- | --- | --- |
| SNPs on ONT data in high-confidence intervals | NanoCaller ONT-HG001 | 98.58 | 98.35 | 98.99 | 98.97 | 98.10 | 98.23 | 97.81 | 98.45 | 99.33 | 98.28 |
|  | NanoCaller ONT-HG002 | 98.63 | **98.66** | **99.09** | **99.11** | **98.38** | 98.43 | 98.06 | 98.60 | **99.34** | **98.44** |
|  | Medaka | **99.03** | 98.59 | 99.02 | 99.04 | 98.17 | 98.50 | 98.24 | **98.94** | 99.24 | 96.94 |
|  | Clair | 98.79 | 97.77 | 98.60 | 98.58 | 97.73 | 97.90 | 97.50 | 98.53 | 98.75 | 90.44 |
|  | Longshot | 98.78 | 98.03 | 97.88 | 97.90 | 98.34 | **98.53** | **98.51** | 98.59 | 98.59 | 98.18 |
| Indels on ONT data in high-confidence intervals | NanoCaller ONT-HG001 | **57.33** | 53.94 | **58.52** | **57.71** | 56.31 | 56.14 | 53.78 | 73.67 | **62.07** | **61.59** |
|  | NanoCaller ONT-HG002 | 56.69 | **54.37** | 58.47 | 57.69 | **56.93** | **56.56** | **54.44** | 73.90 | 61.17 | 60.56 |
|  | Medaka | 48.67 | 48.10 | 53.59 | 50.19 | 55.89 | 52.49 | 51.83 | **81.13** | 51.03 | 53.09 |
|  | Clair | 49.72 | 47.64 | 52.06 | 51.20 | 52.58 | 51.90 | 50.63 | 80.59 | 50.11 | 44.80 |
| Indels on ONT data in non-homopolymer regions | NanoCaller ONT-HG001 | **87.65** | 82.28 | **87.93** | 87.93 | 81.92 | 85.70 | 83.41 | **59.47** | **86.12** | **84.43** |
|  | NanoCaller ONT-HG002 | 87.19 | **82.80** | **87.93** | **88.04** | **82.60** | **86.10** | **83.92** | 59.17 | 85.76 | 83.51 |
|  | Medaka | 82.07 | 78.70 | 85.74 | 84.23 | 80.97 | 84.41 | 82.91 | 55.17 | 78.24 | 78.75 |
|  | Clair | 75.25 | 70.06 | 75.55 | 74.85 | 72.60 | 75.92 | 75.04 | 58.43 | 70.99 | 62.93 |
| SNPs on PacBio CCS data in high-confidence intervals | NanoCaller CCS-HG001 | 99.25 | 99.80 | 99.79 | 99.71 |  |  |  |  |  |  |
|  | NanoCaller CCS-HG002 | 99.17 | 99.80 | 99.79 | 99.75 |  |  |  |  |  |  |
|  | Clair | 99.66 | 99.84 | 99.72 | 99.79 |  |  |  |  |  |  |
|  | Longshot | 99.37 | 99.03 | 99.05 | 99.05 |  |  |  |  |  |  |
|  | DeepVariant | **99.82** | **99.93** | **99.91** | **99.84** |  |  |  |  |  |  |
| Indels on PacBio CCS data in high-confidence intervals | NanoCaller CCS-HG001 | 92.67 | 93.30 | 93.42 | 93.10 |  |  |  |  |  |  |
|  | NanoCaller CCS-HG002 | 93.13 | 94.10 | 94.34 | 93.97 |  |  |  |  |  |  |
|  | Clair | 94.87 | 96.71 | 97.51 | 95.57 |  |  |  |  |  |  |
|  | DeepVariant | **98.21** | **99.28** | **99.48** | **98.42** |  |  |  |  |  |  |
| SNPs on PacBio CLR data in high-confidence intervals | NanoCaller CLR-HG002 | 94.42 | **98.75** | 94.41 | 93.41 |  |  |  |  |  |  |
|  | Clair | 95.83 | 98.38 | **94.89** | **94.15** |  |  |  |  |  |  |
|  | Longshot | **96.81** | 98.41 | 94.35 | 93.27 |  |  |  |  |  |  |

**Figure 3.**
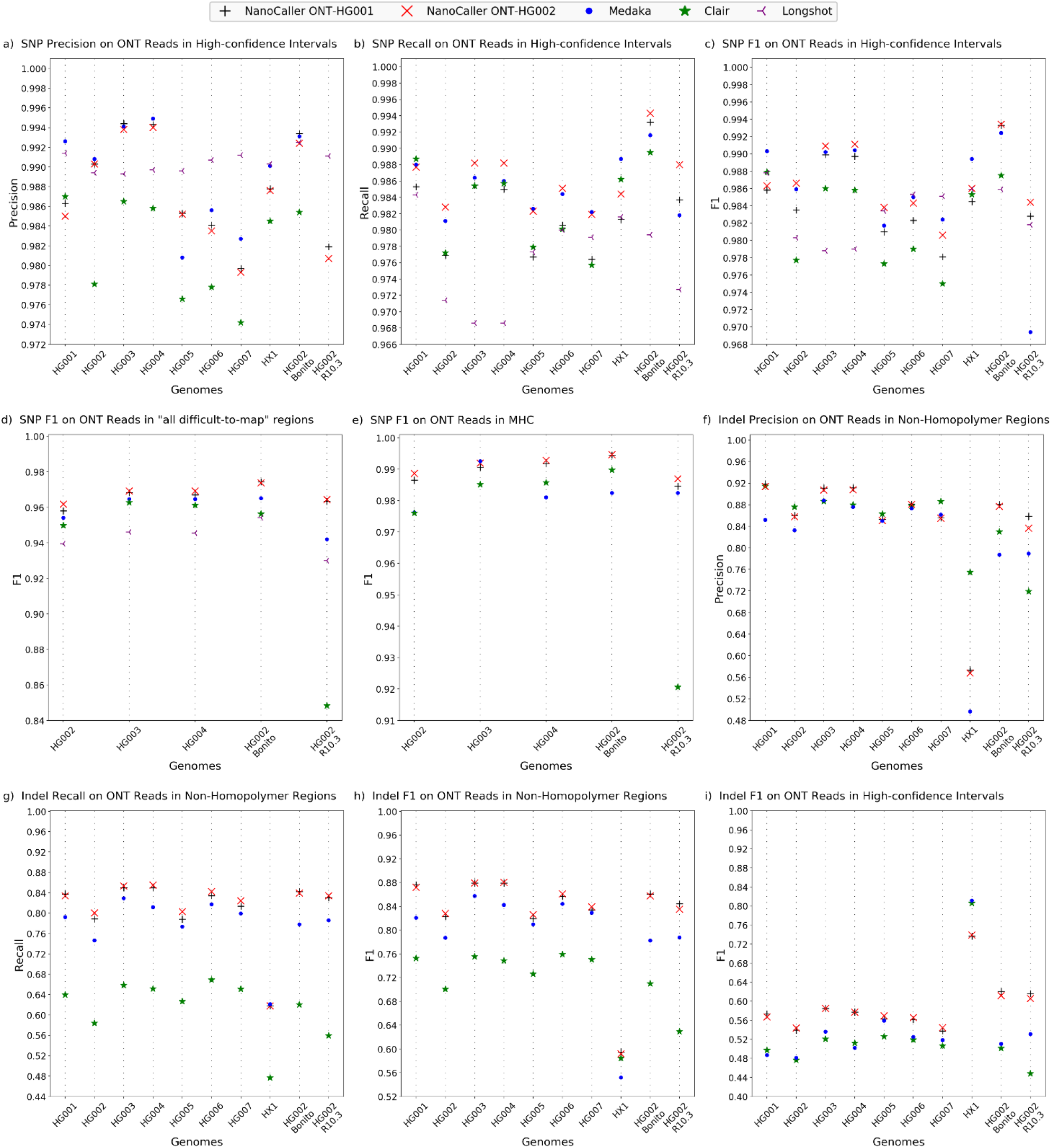
Performances of NanoCaller and other variant callers on ten ONT datasets. SNP performance on whole genome high-confidence intervals: a) precision, b) recall, c) F1 score. F1 scores of SNP performances on: d) “all difficult-to-map” regions, e) MHC. Indel performance non- homopolymer regions: f) precision, g) recall, h) F1 score. F1-score of indel performance in whole genome high-confidence intervals.

On the Ashkenazim trio (HG002, HG003, HG004), NanoCaller has better performance than Medaka, Clair and Longshot in terms of F1-score as shown in Table 2 and Figure 3 (a), (F1-scores are ONT-HG001: 98.35, 98.99, 98.97%; ONT-HG002: 98.66, 99.09, 99.11% vs Medaka: 98.59, 99.02, 99.04% vs Clair: 97.77, 98.60, 98.58% and Longshot: 98.03, 97.88, 97.90%, on HG002, HG003 and HG004 respectively). For example, ONT-HG002 model exceeds Longshot’s by 1.2% F1-score on HG003 and HG004. More details of the performances (precision and recall) can be found in Figure 3 (b) and (c), and ‘Supplementary Materials 2’ Table S30. Furthermore, we evaluated NanoCaller on two additional HG002 ONT datasets. The first data is produced by R10.3 flowcells and basecalled by Guppy 4.0.11, and the second dataset is produced by R9.4.1 flowcells and basecalled with Bonito 0.30. We found that NanoCaller models performed better than other variant callers in terms of F1-score by significant margins. For example, on HG002 ONT data generated by R10.3 flowcells, the F1-scores are 99.33%, 99.34%, 99.24%, 98.75% and 98.59% for ONT-HG001, ONT-HG002, Medaka, Clair and Longshot respectively, whereas on HG002 Bonito dataset, the F1-scores are 98.28%, 98.44%, 96.94%, 90.44% and 98.18% for ONT-HG001, ONT-HG002, Medaka, Clair and Longshot respectively. NanoCaller achieves better F1- score than all other methods. More details for these performances can be found in Figure 3 (a) and (b), and ‘Supplementary Materials 2’ Table S35.

For HG001 and HG005-7, NanoCaller performs competitively against other methods as shown in Table 2, Figure 3: On HG001, F1-scores are 98.58% for ONT-HG001, 98.63% for ONT-HG002, 98.79% for Clair, 98.78 for Longshot and 99.03% for Medaka. On HG005/HG006/HG007, F1-scores are 98.10/98.23/97.81% for ONT-HG001, 98.38/98.43/98.06% for ONT-HG002, 98.17/98.50/98.24% for Medaka, 97.73/97.90/97.50% for Clair and 98.34/98.53/98.51% for Longshot, respectively. It is clear that sometimes NanoCaller has best F1-score (for example on HG005), but sometimes other methods show best F1-score (such as Longshot on HG007 and Medaka on HG001). Please note that benchmark variants of HG001and HG005-7 are older (v3.3.2) than the Ashkenazim trio HG002-4 which has v4.2.1 benchmark variants. The Ashkenazim trio HG002-4 has a larger variant call set than HG001 and HG005- 7 (370-400k more SNPs per genome than HG001), and a larger high-confidence region which includes more difficult genomic regions (at least 200mbp larger than HG001 and covering an extra 7% of the reference genome). This might contribute to the performance variations of different methods.

Next, we show SNP calling performance on HX1 genome sequenced by our lab. On 48x coverage HX1 reads basecalled with Guppy 4.5.2, NanoCaller models perform slightly better Clair and Longshot (F1- scores are 98.45% for ONT-HG001, 98.60% for ONT-HG002, 98.53% for Clair, 98.59% for Longshot, and 98.94% for Medaka). This demonstrates NanoCaller’s ability to accurately identify variants in non-GIAB datasets in real-life applications.

Lastly, we show that the performance of NanoCaller SNP models is independent of the reference genome used. Under this cross-reference testing, we evaluated ONT-HG001 model (trained on reads aligned against GRCh38) on HG002 ONT reads aligned to both GRCh38 and GRCh37 reference genomes. For reads aligned to GRCh38, we obtained 99.03%, 97.69% and 98.35% precision, recall and F1-score respectively. Whereas for reads aligned to GRCh37, we obtained 98.99%, 97.70% and 98.34% precision, recall and F1-score respectively. The similar performance on GRCh38 and GRCh37 indicates that NanoCaller could be used on alignment generated by mapping to different reference genomes.

#### Performance of SNP calling in difficult-to-map genomic regions

We further demonstrate that NanoCaller has a unique advantage in calling SNPs in difficult-to-map genomic regions. We tested both ONT-HG001 SNP model (trained on HG001 ONT reads with v3.3.2 benchmark variants) and ONT-HG002 SNP model (trained on HG002 ONT reads with benchmark variants v4.2.1) on ONT reads of the three genomes of the Ashkenazim trio together with other variant callers. v4.2.1 of GIAB benchmark variants for the trio HG002-4 are used for testing because they have a more exhaustive list of true variants and high confidence intervals in difficult-to-map genomic regions, as shown in ‘Supplementary Materials 2’ Table S27. Difficult-to-map genomic regions here are defined by GA4GH Benchmarking Team and the Genome in a Bottle Consortium and downloaded as BED files from GIAB v2.0 genome stratification. These regions contain all tandem repeats, all homopolymers >6bp, all imperfect homopolymers >10bp, all low mappability regions, all segmental duplications, GC <25% or >65%, Bad Promoters, and other difficult regions such Major Histocompatibility Complex. We intersected the BED files with high-confidence intervals for each genome in the trio and evaluated SNP performance in the intersected regions. As shown in ‘Supplementary Materials 2’ Table S27, each genome has at least 600k SNPs in the intersection of difficult-to-map regions and high-confidence intervals, which is a significant fraction (18-19%) of all SNPs in the high-confidence regions.

The evaluation on these SNPs is shown in Figure 3 (d) and Table 3 for NanoCaller and other methods. For HG002/HG003/HG004, F1-scores are 95.80/96.83/96.70% for ONT-HG001, 96.18/96.92/96.92% for ONT-HG002, 95.41/96.46/96.46% for Medaka, 94.98/96.27/96.12% for Clair, and 93.95/94.61/94.55% for Longshot, respectively). NanoCaller performs better than all other variant callers for each genome. In ‘Supplementary Materials 2’ Table S33, we show performances of NanoCaller SNP models trained on v3.3.2 benchmark variants of HG002 which also perform significantly better than other variant callers. In ‘Supplementary Materials 2’ Table S33, we further show a detailed breakdown of performances in the difficult-to-map regions and demonstrate that NanoCaller generally performs better than other variant callers for SNPs in each of the following categories of difficult-to-map regions: segmental duplications, tandem and homopolymer repeats, low mappability regions, and Major Histocompatibility Complex.

**Table 3.**
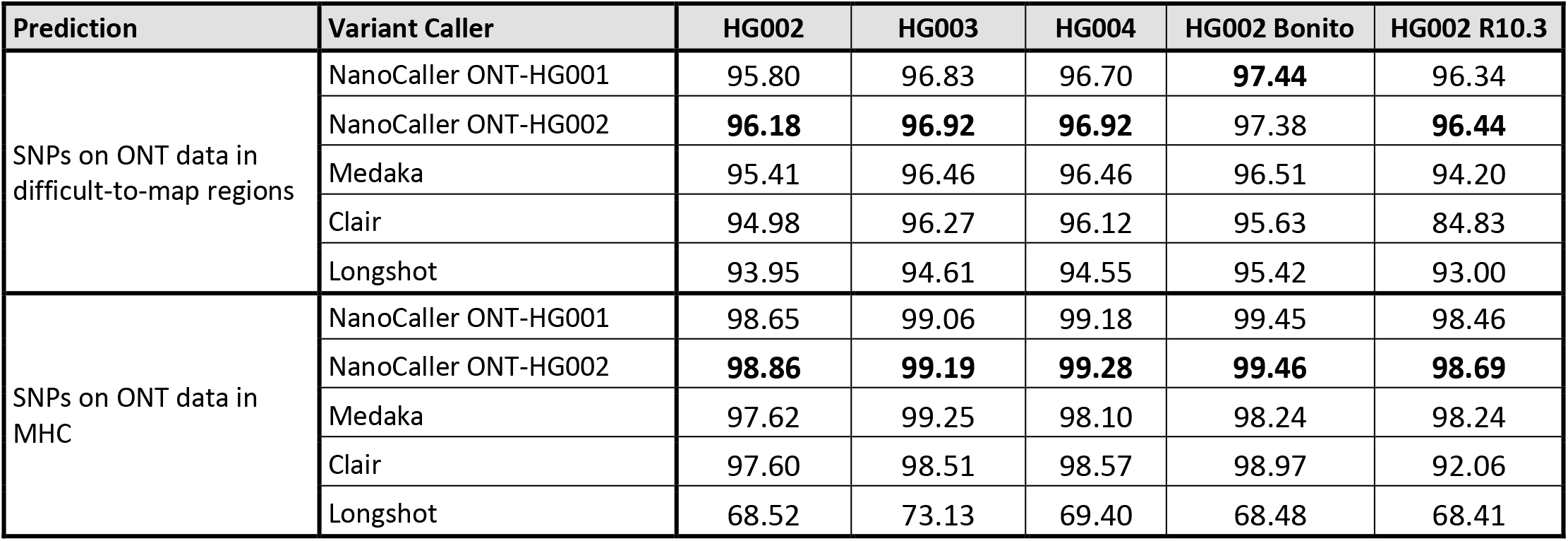
Performances (F1 scores in %) of SNP predictions in difficult-to-map regions and Major Histocompatibility Complex (MHC) by NanoCaller, Medaka, Clair, and Longshot on ONT data. These evaluations are performed against v4.2.1 benchmark variants for the Ashkenazim trio (HG002, HG003, and HG004), whereas “HG002 Bonito” and “HG002 R10.3” are different HG002 ONT datasets.

| Prediction | Variant Caller | HG002 | HG003 | HG004 | HG002 Bonito | HG002 R10.3 |
| --- | --- | --- | --- | --- | --- | --- |
| SNPs on ONT data in difficult-to-map regions | NanoCaller ONT-HG001 | 95.80 | 96.83 | 96.70 | **97.44** | 96.34 |
|  | NanoCaller ONT-HG002 | **96.18** | **96.92** | **96.92** | 97.38 | **96.44** |
|  | Medaka | 95.41 | 96.46 | 96.46 | 96.51 | 94.20 |
|  | Clair | 94.98 | 96.27 | 96.12 | 95.63 | 84.83 |
|  | Longshot | 93.95 | 94.61 | 94.55 | 95.42 | 93.00 |
| SNPs on ONT data in MHC | NanoCaller ONT-HG001 | 98.65 | 99.06 | 99.18 | 99.45 | 98.46 |
|  | NanoCaller ONT-HG002 | **98.86** | **99.19** | **99.28** | **99.46** | **98.69** |
|  | Medaka | 97.62 | 99.25 | 98.10 | 98.24 | 98.24 |
|  | Clair | 97.60 | 98.51 | 98.57 | 98.97 | 92.06 |
|  | Longshot | 68.52 | 73.13 | 69.40 | 68.48 | 68.41 |

To further investigate NanoCaller’s performance, we split difficult-to-map regions into different subgroups according to their length: 0-10kbp, 10-100kbp, 100-500kbp, and >500kbp, and test NanoCaller, Medaka, Clair and Longshot on HG002, HG003 and HG004. We find that when the length of interval increases, the performance advantage of NanoCaller over other methods becomes larger: for example on HG004, NanoCaller’s F1-score is 0.02 higher than Longshot for 0-10kbp subgroup, whereas NanoCaller’s F1- score is 0.1793 higher than Longshot for >500kbp. This demonstrates NanoCaller can benefit SNP calling with long difficult-to-map regions compared to other methods. The details of these performances can be found in ‘Supplementary Materials 2’ Table S41.

Finally, NanoCaller team participated in PrecisionFDA truth challenge v2 for difficult-to-map genomic regions (held in July 2020, see https://precision.fda.gov/challenges/10), and submitted variant calls for the Ashkenazim trio made by an ensemble of NanoCaller, Medaka and Clair models described above. The challenge consisted of using Guppy3.6 basecalled ONT reads for HG003 and HG004 to predict variant calls, which were then evaluated on GIAB v4.1 benchmark variants of HG003 and HG004 that were made public after the challenge ended. At the conclusion of the challenge GIAB released v4.2 benchmark variants. Our ensemble submission won the award for best performance in Major Histocompatibility Complex (MHC) using Nanopore reads [29]. ‘Supplementary Materials 1’ Figure S4 (e) and (f), and ‘Supplementary Materials 1’ Table S11 show the F1 score of SNPs and overall variants performance of the ensemble, NanoCaller model (trained on ONT reads of HG001 basecalled with Guppy2.3.8), Medaka and Clair evaluated during the challenge. While the ensemble performs better than all other variant callers, in general, NanoCaller’s performance on HG002 and HG004 is very close to the ensemble and is significantly better than the performances of Medaka and Clair (F1-scores NanoCaller: 98.53%, 99.07% vs ensemble: 98.97%, 99.17%; Medaka: 97.15%, 94.29% and Clair: 97.55%, 98.59% for HG002 and HG004 respectively). NanoCaller always outperforms Longshot for SNP calling in MHC regions.

Therefore, this is an independent assessment of the real-world performance of NanoCaller in detecting variants in complex genomic regions. For performance of NanoCaller and other variant callers on MHC using latest ONT reads for HG002-4, please refer to Figure 3 (e), Table 3 and Table S33 of ‘Supplementary Materials 2’.

#### Performance on Indel Calling

We tested NanoCaller indel models and other variant callers on ONT reads of eight genomes: HG001-7 and HX1, similar to SNP evaluation. The settings of NanoCaller are given below: to determine indel candidates, thresholds for haplotype insertion allele frequency and deletion frequency were set to 0.4 and 0.6, respectively, due to the abundance of deletion errors in Nanopore reads. Since Nanopore sequencing is unreliable in homopolymer regions, we break down performances for each genome into three categories: high-confidence intervals, homopolymer regions and non-homopolymer regions. Homopolymers regions for indel evaluation are defined as perfect homopolymer regions of length greater than or equal to 4bp as well as imperfect homopolymer regions of length greater than 10bp. Non-homopolymer regions are created by removing homopolymer regions from high-confidence intervals (more details on homopolymer and non-homopolymer regions are shown in ‘Supplementary Material 1’ Pages 5,6, and 27). Performances evaluated by RTG *vcfeval* are shown in Table 2 and Figure 3 (f), (g) and (h) for NanoCaller together with Medaka and Clair. According to the F1 scores in Table 2, NanoCaller performs better than Clair by 8-10% and Medaka by 2-5% in non-homopolymer regions. It is also worth noting that NanoCaller has a higher recall than Medaka and especially Clair: for example, NanoCaller has ∼15-20% and ∼1-4% higher recall than Clair and Medaka respectively on all 7 genomes HG001-7. On high-confidence and homopolymer regions, NanoCaller also achieves higher F1-scores than Medaka and Clair; more details can be found in Table 2 and ‘Supplementary Materials 2’ Table S31. This demonstrates the improved performance of NanoCaller for indel calling. In particular, ‘Supplementary Materials 2’ Figure S7 shows a true insertion and a true deletion in HG002 ONT reads that are predicted correctly by NanoCaller but missed by Clair and Medaka. Both indels show high discordance in position of indels among the long reads, which makes it harder to identify these indels. Furthermore, Figure S1 of ‘Supplementary Materials 1’ further shows concordance of ground truth variants in high-confidence regions (including homopolymer repeat regions) of the Ashkenazim trio correctly predicted by NanoCaller, Medaka and Clair. Figure S1 (b) of ‘Supplementary Materials 1’ shows each tool has a significant number (ranging from 19k-60k) of correctly predicted indel calls, that are not correctly predicted by other variant callers.

### Performance on PacBio Sequencing data

On PacBio datasets of four genomes: HG001, HG002, HG003 and HG004, we evaluated NanoCaller SNP models CCS-HG001 (trained on HG001 CCS reads using benchmark variants v3.3.2) and CCS-HG002 (trained on HG001 CCS reads using benchmark variants v4.2.1). The settings of compared tools are given below: in NanoCaller, minimum alternative allele frequency threshold for SNP calling was set to 0.15 was used to identify SNP candidates for both PacBio CCS and CLR reads, and NanoCaller models are trained with v3.3.2 benchmark variants. For Clair, the PacBio model trained on HG001 and HG005 was used for testing CCS reads, whereas the model trained on seven genomes HG001-HG007 was used for testing CLR reads; both Clair models used v3.3.2 benchmark variants for training; The provided PacBio model in the new DeepVariant release v1.0.0 is used for testing, and this model was trained on CCS reads of HG001-HG006 with v3.3.2 benchmark variants for HG001, HG005-6 and v4.2 for HG002- HG004.

The results for SNP performance on CCS reads are shown in Table 2 and Figure 4 (a), (b) and (c) along with Clair, Longshot, and DeepVariant. It can be seen from Table 2 and Figure 4 that on the Ashkenazim trio, both CCS-HG001 and CCS-HG002 models perform significantly better than Longshot, and NanoCaller shows competitive performance against Clair (F1-scores are CCS-HG001: 99.80, 99.79, 99.71%; CS-HG002: 99.80, 99.79, 99.75% vs Clair: 99.84, 99.72, 99.79% vs Longshot: 99.03, 99.05, 99.05% vs DeepVariant: 99.93, 99.93, 99,84% for G002, HG003 and HG004 respectively). More details performance can be found in ‘Supplementary Materials 2’ Table S37.

**Figure 4.**
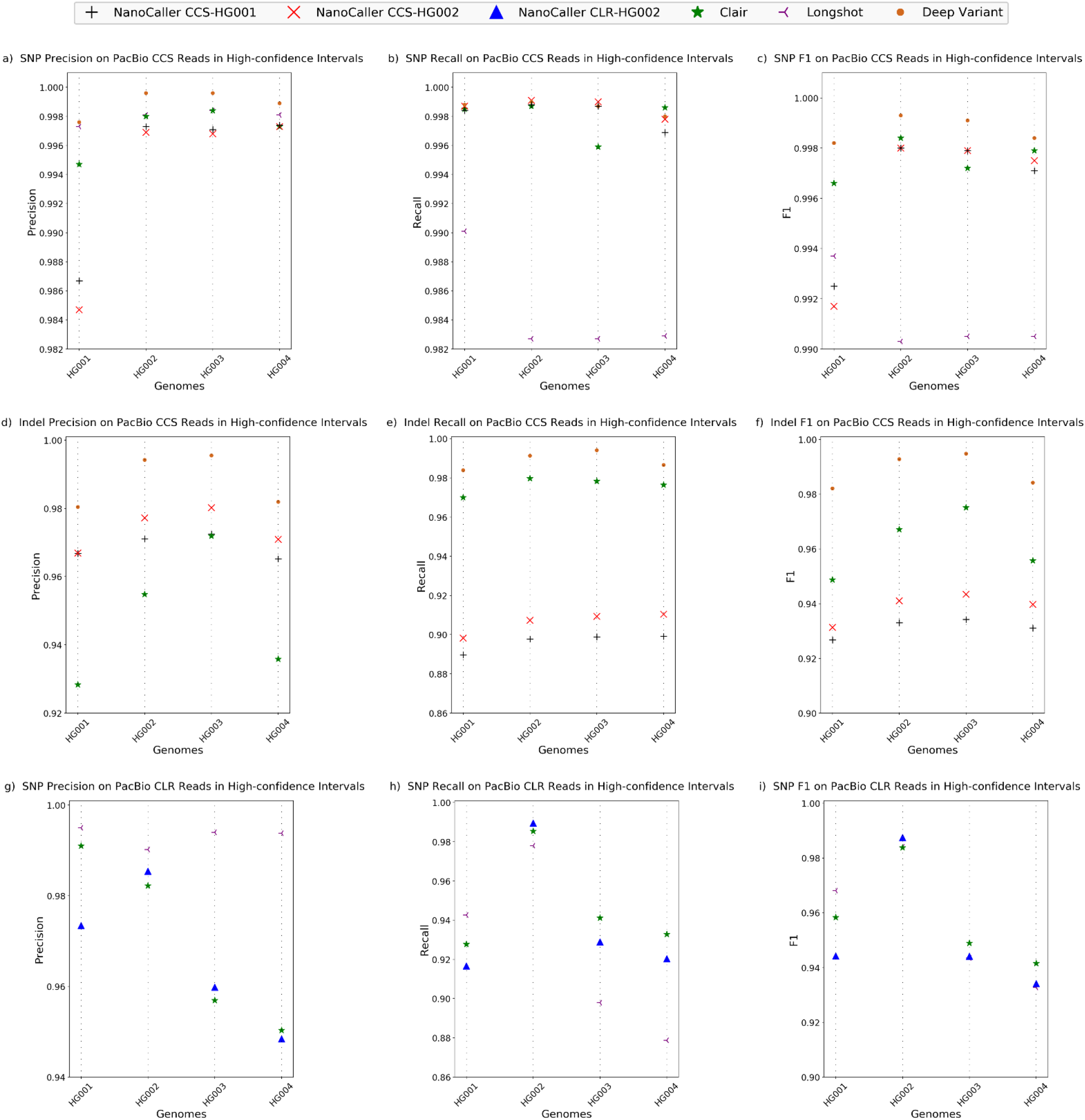
Performances of NanoCaller and other variant callers on four PacBio CCS and four PacBio CLR datasets. SNP performance on whole genome high-confidence intervals using CCS reads: a) precision, b) recall, c) F1 score. Indel performance on whole genome high-confidence intervals using CCS reads: d) precision, e) recall, f) F1 score. SNP performance on whole genome high-confidence intervals using CLR reads: g) precision, h) recall, i) F1 score.

We also evaluated NanoCaller Pacbio indel models CCS-HG001 (trained on HG001 CCS reads using benchmark variants v3.3.2) and CCS-HG002 (trained on HG001 CCS reads using benchmark variants v4.2.1) and the results are shown in Table 2 and Figure 4 (d), (e) and (f) along with Clair and DeepVariant indel performance. The F1-scores on the trio suggest that NanoCaller performs competitively against Clair. As expected, DeepVariant performs very well on CCS reads because CCS reads have much lower error rates. More details of the performance can be found in ‘Supplementary Materials 2’ Table S38.

For PacBio Continuous Long Read Sequencing (CLR) datasets, we evaluated NanoCaller SNP model CLR-HG002 (trained on HG002 PacBio CLR reads using benchmark variants v4.2.1) on the following genomes: HG001 (reads aligned to GRCh37) and the Ashkenazim trio: HG002, HG003 and HG004 (as shown in Table 2 and Figure 4 (g), (h) and (i)). Due to drastic differences in coverage of CLR datasets, we used a higher NanoCaller quality score cut-off for HG003 and HG004, compared to HG001 and HG002. NanoCaller performs competitively Longshot and Clair (F1-scores are CLR-HG002: 98.75, 94.41, 93.41%% vs Clair: 98.38%, 94.89%, 94.15% and Longshot: 98.41%, 94.35%, 93.27%). More details of the performance can be found in Supplementary Table S39.

### Novel Variants called by NanoCaller

We also analyzed SNP calls made by NanoCaller on HG002 (ONT reads basecalled by Guppy 2.3.4) that are absent in the GIAB ground truth calls (version 3.3.2) [30], and validated 17 regions of those SNP calls by Sanger sequencing before v4 benchmark for HG002 was made available (Sanger sequencing signals along with inferred sequences of both chromosomes for each region are shown in the supplementary folder of “Sanger Sequencing Files” and *Sanger_sequences.xlsx)*. By deciphering Sanger sequencing results, we identified 41 novel variants (25 SNPs, 10 insertions and 6 deletions), as shown in Table 5. Based on the 41 novel variants, we conducted the variant calling evaluation by different methods on both older ONT HG002 reads and newly released ONT HG002 reads (as described in the Methods section) to see how more accurate long reads improve variant calling. We find that (1) on the newly released ONT HG002 reads, Medaka correctly identified 15 SNPs, 6 insertions and 2 deletions, Clair identified 14 SNPs, 6 insertions and 2 deletions, and Longshot correctly identified 18 SNPs, while NanoCaller was able to correctly identify 20 SNPs,6 insertions and 2 deletions, as shown in the ‘Supplementary Materials 1’ Table S12, whereas one of these 2 deletions was not called correctly by other variant callers; and (2) on the older ONT HG002 reads, as shown in Table 5, Medaka correctly identified 8 SNPs, 3 insertions and 1 deletion, and Clair identified 8 SNPs, 2 insertions and 1 deletion, whereas Longshot correctly identified 8 SNPs; In contrast, NanoCaller was able to correctly identify 18 SNPs and 2 insertions, whereas 10 of these 18 SNPs and 1 of these 2 insertions were not called correctly by other variant callers on the older HG002 (ONT reads). This indicates that the improvements in per-base accuracy during base-calling significantly enhances the variant calling performance. Also in Table 5, there are 2 multiallelic SNPs which can be identified by NanoCaller but cannot be correctly called by all other 3 methods. One of the multiallelic SNPs at chr3:5336450 (A>T,C) is shown in Figure 5, where both the IGV plots and Sanger results clearly show a multiallelic SNP that was correctly identified by NanoCaller but was missed by other variant callers, likely due to the unique haplotype-aware feature of NanoCaller. In summary, the prediction on these novel variants clearly demonstrate the usefulness of NanoCaller for SNP calling.

**Table 5.**
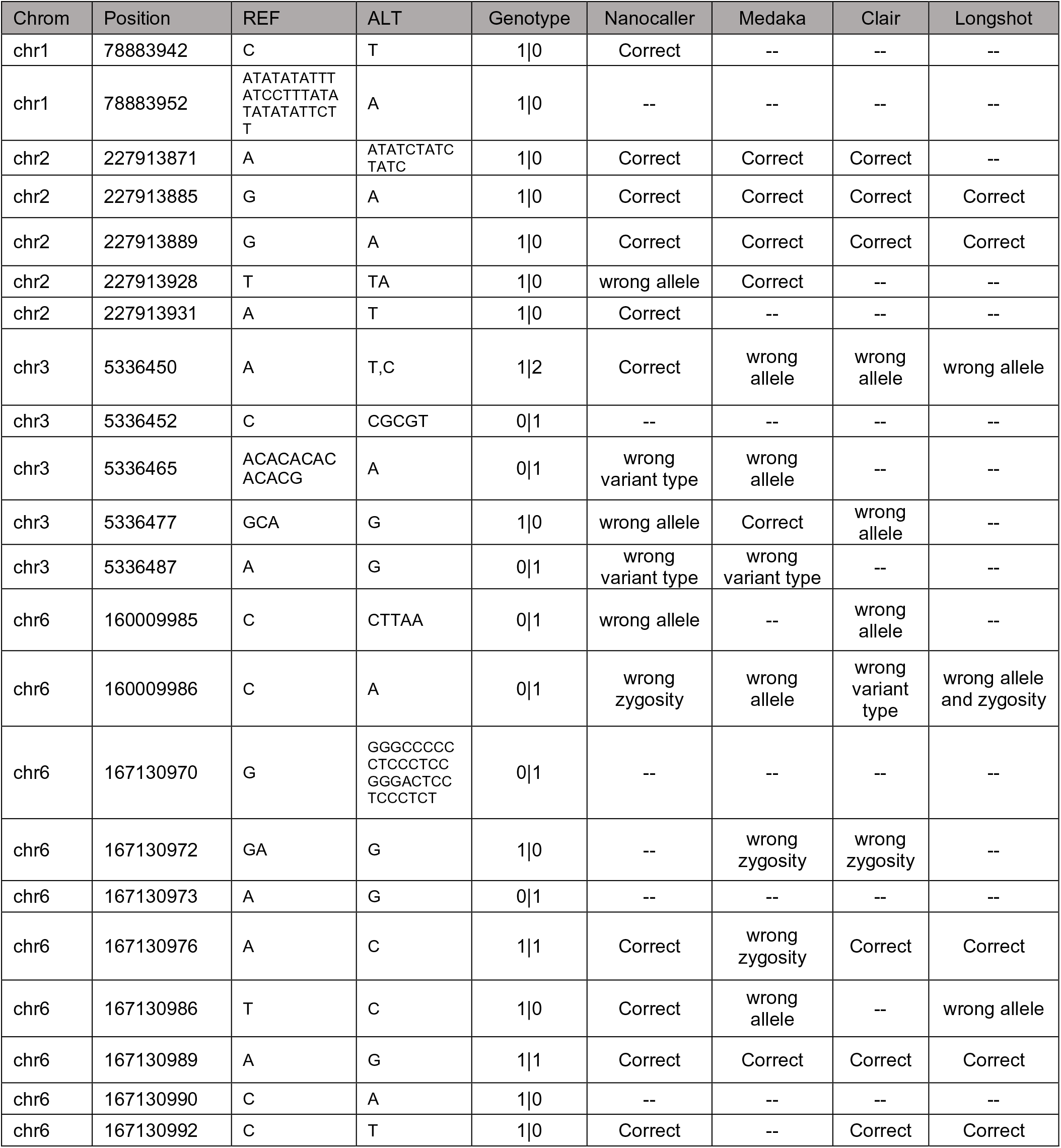
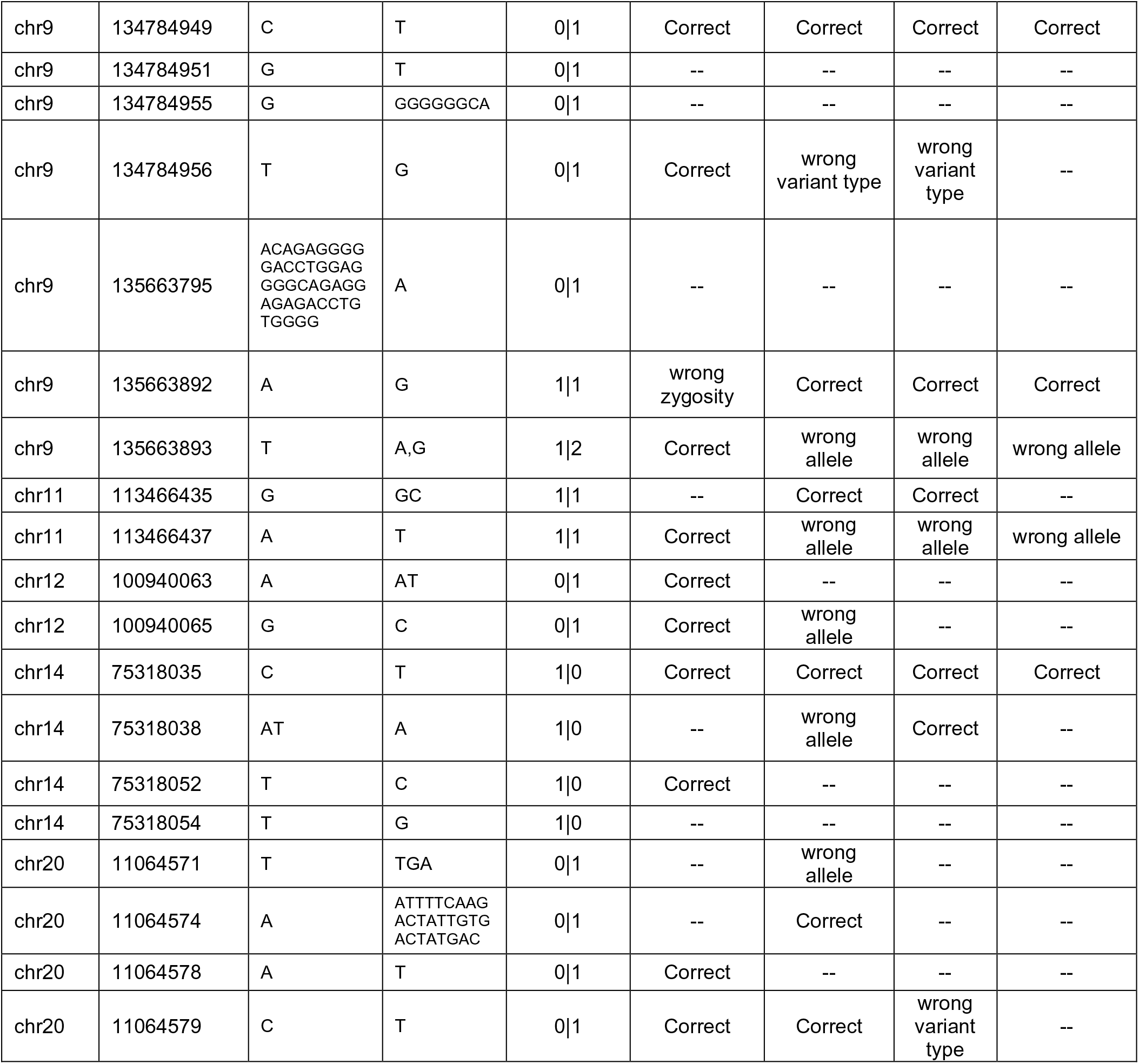
Novel variants in HG002 genome, missing in v3.3.2 benchmark variant, discovered by Sanger sequencing together with the prediction information by NanoCaller and other variant callers using ONT reads basecall with Guppy 2.3.4. NanoCaller model used was trained on ONT HG001 Guppy 2.3.8 basecalled reads.

**Figure 5.**
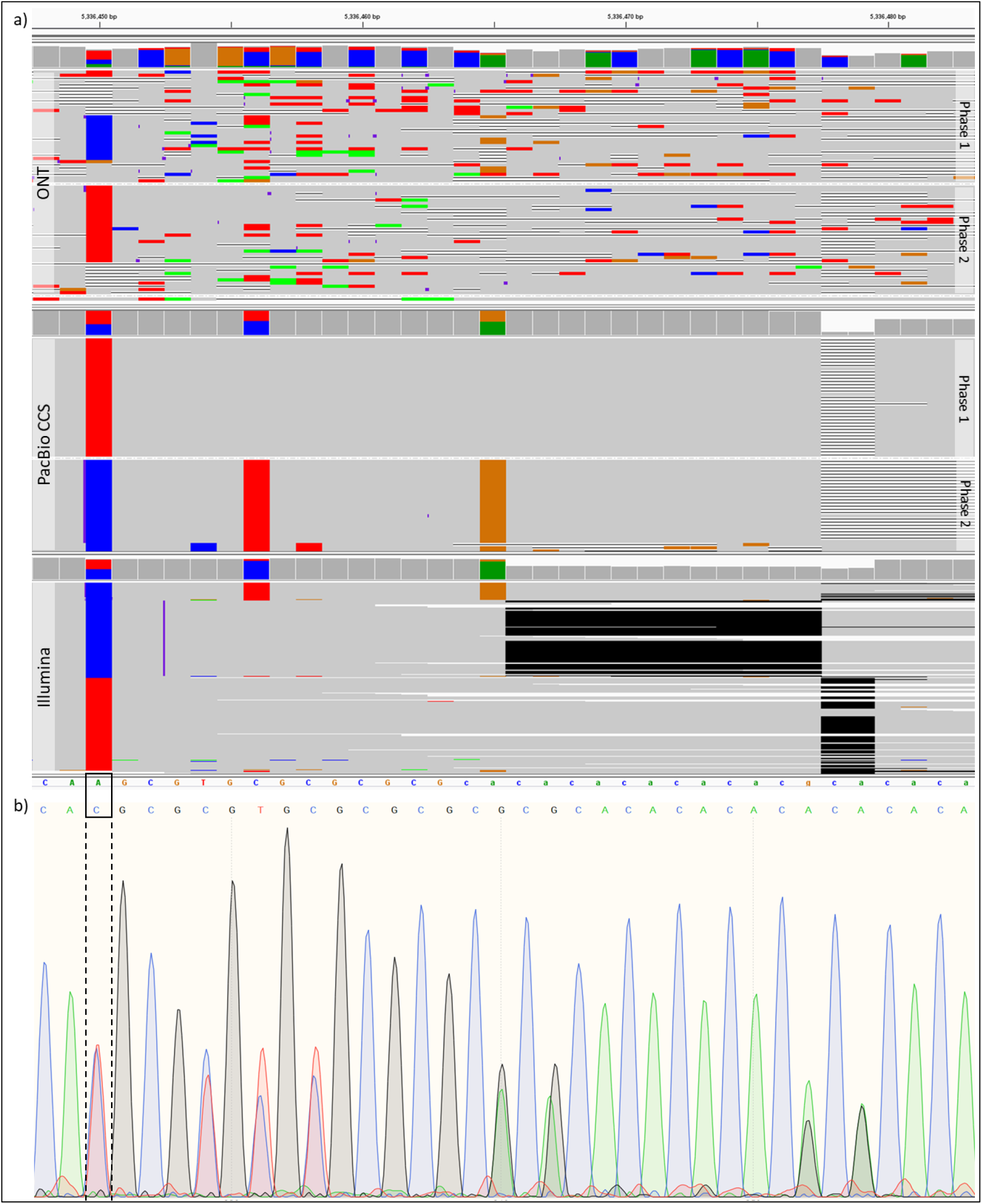
Evidence for novel multiallelic SNP. a) IGV plots of Nanopore, PacBio CCS and Illumina reads of HG002 genome at chr3:5336450-5336480. b) Sanger sequencing signal data for the same region. NanoCaller on older HG002 data correctly identified the multi-allelic SNP at chr3:5336450 (A>T,C) shown in black box.

To demonstrate the performance of NanoCaller for indel calling, we use Figure 6 to illustrate those variants that can be detected by long-read variant callers but cannot be detected by short-read data. In Figure 6, the validated deletion is at chr9:135663795 or chr9:135663804 (there are two correct alignments at the deletion and thus both genomic coordinates are correct.). NanoCaller detects the deletion at chr9:135663805, while Medaka and Clair detect the deletion at chr9:135663799. Although they are several bps away from the expected genomic coordinates (which is normal in long-read based variant calling), the prediction provides accurate information of the deletion compared with short-read data where little evidence supports the deletion as shown in Figure 6 (a). Sanger sequencing signal data, shown in Figure 6 (b), confirms the presence of a heterozygous deletion at the same location which is causing frameshift between the signals from maternal and paternal chromosomes. This is an example to demonstrate how long-read variant callers on long-read data can detect variants that fail to be reliably called on short-read sequencing data.

**Figure 6.**
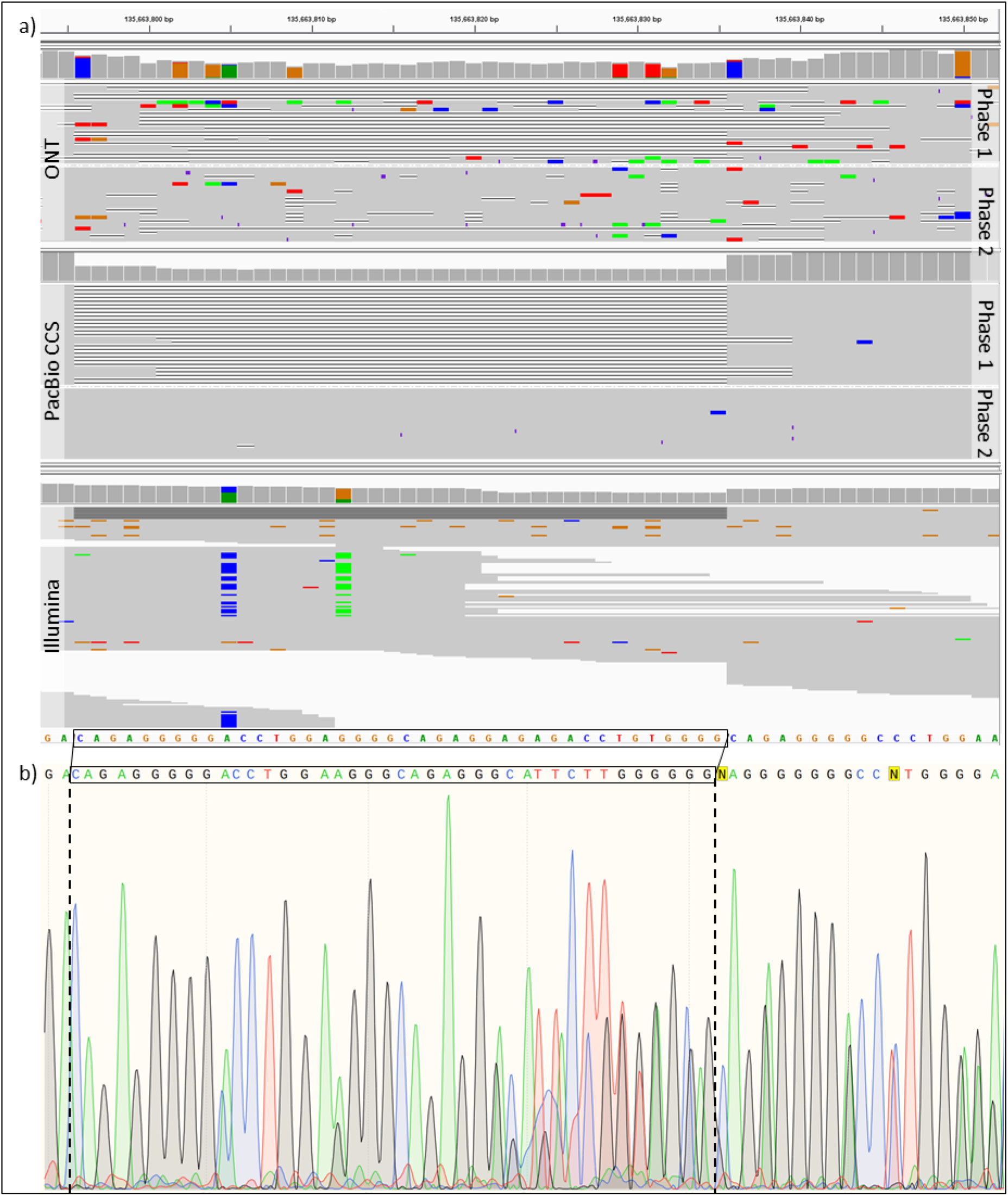
Evidence for novel deletions. a) IGV plots of Nanopore, PacBio CCS and Illumina reads of HG002 genome at chr9:135663780-chr9:135663850. The 40bp long deletion shown below in black box was identified using Sanger sequencing at chr9:135663795 or chr9:135663804 (both are correct and the difference is due to two different alignments). b): Sanger sequencing signal data around the deletion.

### NanoCaller Runtime Comparison

We assessed NanoCaller’s running time in four modes: ‘snps_unphased’, ‘snps’, ‘indels’, and ‘both’. In ‘snps_unphased’ mode, NanoCaller uses deep neural network model to predict SNP calls only, whereas in the ‘snps’ mode, NanoCaller SNP calling is followed by an additional step of phasing SNP calls by external haplotyping tools such as WhatsHap. In the ‘indels’ mode, NanoCaller uses phased reads in a BAM input to predict indels only. The entire NanoCaller workflow is the ‘both’ mode, where NanoCaller first runs in ‘snps’ mode to predict phased SNP calls, then uses WhatsHap to phase reads with the SNP calls from the ‘snps’ mode, followed by running ‘indels’ mode on phased reads from the previous step.

Table 4 shows the wall-clock runtime of each mode of NanoCaller using 16 CPUs (IntelXeon CPU E5- 2683 v4 @ 2.10GHz) on 49x HG002 ONT (reads basecalled by Guppy 3.6), 35x PacBio CCS (15kn library size) and 58x PacBio CLR reads. NanoCaller takes ∼18.4 hours and ∼2.8 hours to run ‘both’ and ‘snps_unphased’ modes on 49x HG002 ONT reads, compared to ∼181.6 hours for Medaka and ∼5.6 hours for Clair, on the same 16 CPUs. On 35x CCS reads, NanoCaller takes ∼11.2 hours and ∼2.7 hours to run to run ‘both’ and ‘snps_unphased’ modes, compared to ∼1.8 hours by Clair and ∼11.8 by DeepVariant, on 16CPUs. NanoCaller usually runs faster than other tools. We summarize the runtime of all variant callers in ‘Supplementary Materials 1 ’Table S17.

**Table 4.** Wall-clock runtime in hours of different modes of NanoCaller using 16 CPUs on 49x ONT, 35x CCS and 58x CLR reads of HG002.

| NanoCaller Mode | snps_unphased | snps | indels | both |
| --- | --- | --- | --- | --- |
| Description | Only SNP calling by NanoCaller deep convolutional network model | 'snps_unphased' mode followed by WhatsHap SNP call phasing | Only indel calling by NanoCaller using phased BAM input | 'snps' mode, followed by read phasing by WhatsHap, and 'indels' mode |
| ONT | 3.8 | 5.0 | 12.6 | 18.4 |
| CCS | 2.7 | 3.2 | 7.6 | 11.2 |
| CLR | 3.6 | 4.8 | -- | -- |

Please note that Medaka’s first step also produces unphased SNP calls using a recurrent neural network on mixed haplotypes (Medaka later uses WhatsHap to phase SNP calls and reads for haplotype separated variant calling). Compared with Medaka’s first step, NanoCaller’s unphased SNP calling not only takes a fraction of the time required for Medaka’s first step (∼2.8 hours vs ∼70.7 hours), but also gives much better performance (precision, recall and F1-score on ONT HG002 (reads basecalled by Guppy 3.6) are NanoCaller: 98%, 97.99%, 97.99% vs Medaka 98.01%, 92.16%, 94.99%). Similarly, Longshot’s first step uses a pair-HMM model to produce SNP calls from mixed haplotypes (Longshot later uses HapCUT2 to update the genotypes of these SNP calls in an iterative manner); on HG002 ONT (reads basecalled by Guppy 3.6), Longshot’s first step takes 15.2 hours with 93.01%, 95.69%, and 94.33% precision, recall and F1-score respectively. Longshot and WhatsHap cannot use multiple CPUs to produce on SNP calls. With single CPU on 49x HG002 ONT reads, Longshot needs ∼49.7 hours and WhatsHap needs ∼84.3 hours for SNP calling.

### Effects of Various Parameters on NanoCaller’s Performance

In this section, we discuss the effects of tuning various parameters on NanoCaller’s performance. The NanoCaller SNP models presented here are: NanoCaller1 (trained on ONT HG001 reads basecalled with Guppy 2.3.8 using benchmark variants v3.3.2), NanoCaller2 (trained on ONT HG002 reads basecalled with Guppy 2.3.4 using benchmark variants v3.3.2), and NanoCaller3 (trained on HG003 PacBio CLR reads using benchmark variants v3.3.2). For testing, we used ONT reads for HG002-4 basecalled with Guppy3.6, CCS reads for HG002-4 (15kb library size), and HG002-4 CLR reads.

#### Strategies of choosing heterozygous SNPs for SNP features generation

In NanoCaller, we generate input features for a SNP candidate site by choosing potentially heterozygous SNP sites that share a read with the candidate site. In the implementation of NanoCaller, at most 20 heterozygous SNP candidates are chosen for downstream and upstream of a candidate site of interest. With an expectation that 1 SNP occurring per 1000bp, a simple way is to include 20kb downstream and upstream sequence centered at the candidate site of interest, and then select 20 nearest heterozygous SNP sites. But in some smaller genomic regions, a cluster of heterozygous SNP candidates may be found but due to the noise with many false positives, and these false SNPs in a very smaller regions would provide a strong co-occurrence evidence for each other but little information for the candidate site of interest. Longshot notices this issue and overcomes high false positive rates due to dense clusters of false positive SNP by simply removing these dense clusters if the number of SNP calls exceeds a certain threshold in a specified range; however applying hard limits like that can lead to missing out on true SNPs that do occur in dense clusters in certain genomic regions.

We decide to use a difference method for selecting nearby potential heterozygous sites by forcing NanoCaller to pick a certain number of site that were some distance away from the candidate site. More precisely, we force NanoCaller to pick 2, 3, 4, 5 and 6 heterozygous SNP sites between the following distances from the candidate site: 0, 2kbp, 5kbp, 10kbp, 20kbp and 50kbp. This is illustrated in Figure S3 and Table S18 of ‘Supplementary Materials 1’. We found that using this method, we achieve better SNP calling performance for ONT reads; ‘Supplementary Materials 1’ Table S21 shows that under this strategy, for each genome in the Ashkenazim trio, we achieve higher precision, recall and F1 score for whole genome analysis as well as in each difficult-to-map genomic regions. On the other hand, SNP calling performance of PacBio CCS and CLR reads is not affected by using this method of selecting heterozygous SNP sites, as shown in “Supplementary Materials 1” Table S22. This might be due to the fact that ONT reads have significantly higher N50 and mean read lengths compared to PacBio CCS and CLR reads, as shown in ‘Supplementary Materials 1’ Table S1. ‘Supplementary Materials 1’ Figure S2 shows the read lengths distribution of HG004 ONT, CCS and CLR datasets of 88X, 35X and 27X coverages respectively. In these datasets, 99.4% of CCS reads and 97% of CLR reads are shorter than 20,000bp; on the other hand, only 69.3% and 87.4% of the ONT reads are shorter than 20,000bp and 50,000bp. This simple comparison clearly demonstrates that NanoCaller is able to utilize the longer reads to improve SNP calling, and comparatively shorter PacBio reads might in part contribute to the less improvement of NanoCaller. Thus, as read length increases, we expect NanoCaller can have better performance.

#### Different thresholds for heterozygous SNP sites for SNP features generation

In order to generate haplotype structure features from long reads for a SNP candidate site, we need to select potentially heterozygous sites. Ideally heterozygous sites should have approximately 0.5 alternative allele frequency, which is rarely the case due to alignment and sequencing errors. Therefore, a SNP candidate site is determined to potentially heterozygous if its alternative allele frequency is in a small range centered at 0.5: typically this range is 0.4-0.6 or 0.3-0.7 depending upon the sequencing technology and is called neighbor threshold. Table S19 of “Supplementary Materials 1” shows how the choice of this threshold effects SNP calling performance for ONT, PacBio CCS and CLR reads, which have different characteristics of error rates and read lengths (which in turn determines the number of candidate sites). We can observe that, generally, using a narrower range around 0.5% allows higher precision, but recall decreases because not enough heterozygous sites are chosen to give informative features. In particular, the performance is very sensitive to increases in the upper limit of threshold, and decreases drastically when the threshold is increased. We determined that 0.4-0.6 threshold works best for ONT reads, with 0.3-0.7 and 0.3-0.6 being the best thresholds for CCS and CLR reads. Using a narrower threshold for ONT reads makes sense since longer ONT reads give us plenty of heterozygous sites to choose from, compared to CLR or CCS reads. It should be noted that this threshold is used for testing a sequencing data only, and during training of NanoCaller SNP models, we simply use benchmark heterozygous SNPs.

We check how minimum numbers of heterozygous SNP candidates for NanoCaller affect the performance and show the result in Table S20 of Supplementary materials. In NanoCaller, a SNP candidates with less than a minimum number of heterozygous SNP candidates will be considered as false negatives without prediction. By default, a minimum number of heterozygous SNP candidates is 1. In Table S20 of “Supplementary Materials 1”, different minimum numbers of heterozygous SNP candidates are checked on Nanopore reads and Pacbio reads. On both data, as this minimum threshold increase, precision increases and recall decreases. The increasing precision suggests that more heterozygous SNP candidates can benefit SNP prediction.

#### Using WhatsHap ‘distrust genotype’ option for phasing

WhatsHap is able to call SNPs on ONT and PacBio reads, as shown in ‘Supplementary Materials 1’ Table S26. WhatsHap shows similar performance to NanoCaller, albeit much slower, while for ONT reads, WhatsHap shows poor performance with F1-scores around 88-93%. In NanoCaller, WhatsHap is used for phasing SNPs and reads but not for variant calling. Further, WhatsHap allows ‘distrust genotype’ setting for phasing which allows WhatsHap to change genotypes of any SNP, from hetero- to homozygous and vice versa, in an optimal likelihood solution based upon the haplotypes created. In NanoCaller, this setting is disabled by default.

If users of NanoCaller wants to use ‘distrust genotype’ setting in WhatsHap, negligible effect of SNP calling performance is expected on ONT reads, while an increase in F1-score of 0.15-0.5% and 0.17- 1.4% are expected on PacBio CCS and CLR reads, as shown in Table S23 of ‘Supplementary Materials 1’. Nevertheless, one should note that using this setting will significantly increase the runtime.

## Discussion

In this study, we present NanoCaller, a deep learning framework to detect SNPs and small indels from long-read sequencing data. Depending on library preparation and sequencing techniques, long-read data usually have much higher error rates than short-read sequencing data, which poses a significant challenge to variant calling and thus stimulates the development of error-tolerant deep learning methods for accurate variant calling. However, the benefits of much longer read length of long-read sequencing are not fully exploited for variant calling in previous studies. The NanoCaller tool that we present here solely integrates haplotype structure in deep convolutional neural network for the detection of SNPs from long-read sequencing data, and uses multiple sequence alignment to re-align indel candidate sites to generate indel calling. Our evaluations under the cross-genome testing, cross-reference genome testing, and cross-platform testing demonstrate that NanoCaller performs competitively against other long-read variant callers, and outperforms other methods in difficult-to-map genomic regions.

NanoCaller has several advantages to call variants from long-read sequencing data. (1) NanoCaller uses pileup of candidate SNPs from haplotyped set of long-range heterozygous SNPs (with hundreds or thousands bp away rather than adjacent neighborhood local region of a candidate SNP of interest), each of which is shared by a long read with the candidate site. Given a long read with >20kb, there are on averagely >20 heterozygous sites, and evidence of SNPs from the same long reads can thus improve SNP calling by deep learning. Evaluated on several human genomes with benchmarking variant sets, NanoCaller demonstrates competitive performance against existing variant calling methods on long reads and with phased SNPs. (2) NanoCaller is able to make accurate predictions cross sequencing platforms and cross reference genomes. In this study, we have tested NanoCaller models trained on Nanopore data for performance. We also test NanoCaller models calling variants on PacBio long-read data and achieved similar prediction trained on GRCh38 for GRCh37 and achieve the same level SNP calling performance. (3) With the advantage of long-read data on repetitive regions, NanoCaller is able to detect SNPs/indels outside high-confidence regions which cannot be reliably detected by short-read sequencing techniques, and thus NanoCaller provides more candidate SNPs/indels sites for investigating causal variants on undiagnosed diseases where no disease-causal candidate variants were found by short-read sequencing. (4) NanoCaller uses rescaled statistics to generate pileup for a candidate site, and rescaled statistics is independent on the coverage of a test genome, and thus, NanoCaller is able to handle a test data set with different coverage from the training data set, which might be a challenge for other long-read callers. That is, NanoCaller trained on a whole-genome data has less biases on other data sets with much lower or higher coverage, such as target-sequencing data with thousands folds of coverage. (5) With very accurate HiFi reads (<1% error rate) generated by PacBio, NanoCaller is able to yield competitive variant calling performance. (6) NanoCaller has flexible design to call multi-allelic variants, which Clairvoyante and Longshot cannot handle. In NanoCaller, the probability of each nucleotide type is assessed separately, and it is allowed that the probability of 2 or 3 or 4 nucleotide type is larger than 0.5 or even close to 1.0, and thus suggests strong evidence for a specific position with multiple bases in a test genome. Therefore, NanoCaller can easily generate multi-allelic variant calls, where all alternative alleles differ from the reference allele.

However, there are several limitations of NanoCaller that we wish to discuss here. One is that NanoCaller relies on the accurate alignment and pileup of long-read sequencing data, and incorrect alignments in low-complexity regions might still occur, complicating the variant calling process. For instance, most variants missed by NanoCaller in MHC region cannot be observed through IGV either due to alignment errors. Both continuingly improved sequencing techniques and improved alignment tools can benefit NanoCaller with better performance. But if the data is targeted at very complicated regions or aligned with very poor mapping quality, the performance of NanoCaller would be affected. Another limitation of NanoCaller is that the indel detection from mononucleotide repeats might not be accurate, especially on Nanopore long-read data which has difficulty in the basecalling of homopolymers [31, 32]. In Nanopore long-read basecalling process, it is challenging to determine how many repeated nucleotides for a long consecutive array of similar Nanopore signals, potentially resulting in false indel calls at these regions, which can be post-processed from the call set. Please also note that although NanoCaller might be able to call somatic multi-allelic variants in tumor samples with clonal heterogeneity and variable tumor content, NanoCaller is currently designed to call diploid alleles. However, the frequency of some somatic variants in tumor samples might be too low to be distinguished from noises in long reads. Therefore, the variant calling on tumor samples needs a careful design and parameter tuning if NanoCaller is used. Additionally, better performance could be achieved for a specific training model when more benchmarking tumor data sets are available.

In summary, we propose a deep-learning tool solely using long-range haplotype information for SNP calling and local multiple sequence alignments for accurate indel calling. Our evaluation on several human genomes suggests that NanoCaller performs competitively against other long-read variant callers and can generate SNPs/indels calls in complex genomic regions. NanoCaller enables the detection of genetic variants from genomic regions that are previously inaccessible to genome sequencing and may facilitate the use of long-read sequencing in finding disease variants in human genetic studies.

## Methods

### Datasets

#### Long-read data

Long-read data sets for eight human genomes are used for the evaluation of NanoCaller: HG001, the Ashkenazim trio (consisting of son HG002, father HG003 and mother HG004), the Chinese trio (consisting of son HG005, father HG006 and mother HG007) and HX1. For HG001-7, Oxford Nanopore Technology (ONT) FASTQ files basecalled with Guppy 4.2.2 were downloaded from Human Pangenome Reference Consortium Database. HX1 genome was sequenced by us using PacBio [10] and Nanopore sequencing [35], and Nanopore reads for HX1 were re-basecalled using Guppy 4.5.2. All ONT datasets were aligned to GRCh38 using minimap2 [33]. PacBio CCS alignment files for HG002 and HG003, and FASTQ files for HG001 and HG004 were downloaded from the GIAB database [30, 34]; FASTQ files were aligned to GRCh38 reference genome using minimap2. These CCS datasets were prepared with 15k and 20kb library size selection, and therefore have longer reads than CCS reads used for precisionFDA challenge. PacBio CLR alignment reads for HG001-4 were downloaded from the GIAB database [30, 34]. ‘Supplementary Materials 2’ Table S29 shows the statistics of mapped reads in the eight genomes where the coverage of ONT data ranges from 34 to 84 and the coverage of PacBio data is between 27 and 58.

#### Benchmark variant calls

The benchmark set of SNPs and indels for HG001 (version 3.3.2), the Ashkenazim trio HG002-4 (v4.2.1 and v3.3.2), and the Chinese trio HG005-7 (v3.3.2) are download from the Genome in a Bottle (GIAB) Consortium [30] together with high-confidence regions for each genome. There are 3,002,314; 3,459,843; 3,430,611; 3,454,689; 2,945,666; 3,030,507; 3,048,404; 3,489,068 SNPs for HG001, HG002, HG003, HG004, Hg005, HG006 and HG007 respectively, and 517,177;5 87,987; 569,180; 576,301; 432,747; 435,520; 437,866; 697,736 indels for them, as shown in Table 1. Benchmark variant calls for HX1 were generated by using GATK on Illumina ∼300X reads sequenced by us [10].

### NanoCaller framework for variant calling

In the framework of NanoCaller for variant calling, candidate sites of SNPs and indels are defined according to an input alignment and a reference sequence. NanoCaller has two convolutional neural networks, one for SNP calling and the other for indel prediction, each requiring a different type of input. Input pileup images generated for SNP candidate sites only use long-range haplotype information. For

indel candidate sites, alignment reads are phased with WhatsHap using SNP calls from NanoCaller, and then NanoCaller uses phased alignment reads to generate input pileup images by carrying out local multiple sequence alignment around each site. Afterwards, NanoCaller combines SNP and indel calls to give a final output. The details are described below.

#### SNP Calling in NanoCaller

There are four steps in NanoCaller to generate SNP calling result for an input genome: candidate site selection, pileup image generation of haplotype SNPs, deep learning prediction, and phasing of SNP calls.

##### Candidate site selection

Candidate sites of SNPs are defined according to the depth and alternative allele frequency for a specific genomic position. In NanoCaller, “SAMtools mpileup” [36] is used to generate aligned bases against each genomic position. In NanoCaller, SNP candidate sites are determined using the criteria below. For a genomic site *b* with reference base *R*,

1. 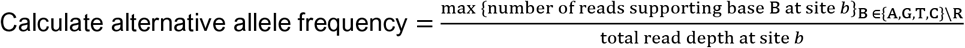
2. *b* is considered a SNP candidate site if the total read depth and the alternative allele frequency are both greater than specified thresholds. We set the alternative allele frequency threshold to be 15%.

##### Pileup image generation

After selecting all SNP candidate sites above, we determine a subset of SNP candidate sites as the set of highly likely heterozygous SNP sites (denoted by V). We extract long-range haplotype information from this subset of likely heterozygous SNP sites to create input images for all SNP candidate sites to be used in a convolutional neural network. This subset consists of SNP candidate sites with alternative allele frequencies in a specified range around 50%, and the default range is 40% to 60% for heterozygous site filtering. This range can be specified by the user depending upon the sequencing technology and read lengths. In detail, the procedure of pileup image generation is described below (as shown in Figure 1). For a SNP candidate site *b*:

1. We select sites from the set V that share at least one read with *b* and are at most 50,000bp away from *b*. For SNP calling on PacBio datasets, we set this limit at 20,000bp.
2. In each direction, upstream or downstream, of the site *b*, we choose 20 sites from V. If there are less than 20 such sites, we just append the final image with zeros. We denote the set of these potential heterozygous SNP sites nearby *b* (including *b)* by Z. An example is shown in Figure 1 (a). More details for how to choose these 40 nearby heterozygous sites from the set V can be found at ‘Supplementary Materials 1’ Tables S18-S22 and Figures S2-S3.
3. The set of reads covering *b* is divided into four groups, R_*B*_ = {reads that support base B at b}, B ∈ {A, G, T, C}. Reads that do not support any base at *b* are not used.
4. For each read group in R_*B*_ with supporting base B, we count the number 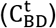 of supporting reads for site *t* ∈ Z with base D ∈ {A, G, T, C}.
5. Let 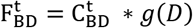, where g(*D*) is a function that returns -1 if *D* is the reference base at site *t* and 1 otherwise. An example is shown in Figure 1(c).
6. We obtain a 4x41x4 matrix M with entries 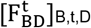 (as shown Figure 1(d)) where the first dimension corresponds to nucleotide type B at site *b*, second dimension corresponds to the number of sites *t*, and the third dimension corresponds to nucleotide type D at site *t*. Our image has read groups as rows, various base positions as columns, and has 4 channels, each recording frequencies of different bases in the given read group at the given site.
7. We add another channel to our image which is a 4x41 matrix 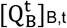 where 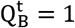 if B is the reference base at site *b* and 0 otherwise (as shown in Figure 1(d)). In this channel, we have a row of ones for reference base at *b* and rows of zeroes for other bases.
8. We add another row to the image which encodes reference bases of site in Z, and the final image is illustrated in Figure 1(e).

##### Deep learning prediction

In NanoCaller, we present a convolutional neural networks [37] for SNP prediction, as shown in Figure S6 of “Supplementary Materials 1”. The neural network has three convolutional layers: the first layer uses kernels of three different dimensions and combines the convolved features into a single output: one capture local information from a row, another from a column and the other from a 2D local region; the second and third layers use kernels of size 2x3. The output from third convolutional layer is flattened and used as input for a fully connected network with dropout (using 0.5 drop date). The first fully connected layer is followed by two different neural networks of fully connected layers to calculate two types of probabilities. In the first network, we calculate the probability of each nucleotide type B to indicate that B is present at the genomic candidate site; thus for each nucleotide type B, we have a binary label prediction. The second network combines logit output of first network with output of first fully connected hidden layer to estimate probability for zygosity (homozygous or heterozygous) at the candidate site. The second network is used only in the training to propagate errors backwards for incorrect zygosity predictions. During testing, we infer zygosity from the output of first network only.

In order to call SNPs for a test genome, NanoCaller calculates probabilities of presence of each nucleotide type at the candidate site. If a candidate site has at least two nucleotide types with probabilities exceeding 0.5, it is considered to be heterozygous, otherwise it is regarded as homozygous. For heterozygous sites, two nucleotide types with highest probabilities are chosen with a heterozygous variant call. For homozygous sites, only the nucleotide type with the highest probability is chosen: if that nucleotide type is not the reference allele, a homozygous variant call is made, otherwise, it is homozygous reference. Each of called variants is also assigned with a quality score which is calculated as −100 log_10_ *Probability* (1 − *P*(*B*)), where *P*(*B*) is the probability of the alternative *B* (in case of multiallelic prediction we choose *B* to be the alternative allele with smaller probability) and recorded as a float in QUAL field of the VCF file to indicate the chance of false positive prediction: the larger the score is, the less likely that the prediction is wrong.

##### Phasing of SNP calls

After SNP calling, NanoCaller phases predicted SNP calls using WhatsHap [38]. By default, NanoCaller disables ‘distrust-genotypes’ and ‘include-homozygous’ settings of WhatsHap for phasing SNP calls, which would otherwise allow WhatsHap to switch variants from hetero- to homozygous and vice versa in an optimal phasing solution. Enabling these WhatsHap settings has minimal impact on NanoCaller’s SNP calling performance (as shown in the Supplementary Tables S2, S3 and S4), but increases the time required for phasing by 50-80%. NanoCaller outputs both unphased VCF file generated by NanoCaller and phased VCF file generated by WhatsHap.

#### Indel Calling in NanoCaller

Indel calling in NanoCaller takes a genome with phased reads as input and uses several steps below to generate indel predictions: candidate site selection, pileup image generation, deep learning prediction, and then indel sequence determination. In NanoCaller, long reads are phased with SNPs calls that are predicted by NanoCaller and phased by WhatsHap [38] (as described above).

**Candidate site selection:** Indel candidate sites are determined using the criteria below. For a genomic site *b*,

1. Calculate:

a. For *i* ∈ {0, 1}, depth_*i*_ = total number of reads in phase *i* at site *b*
b. 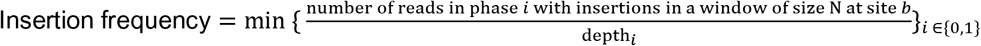
c. 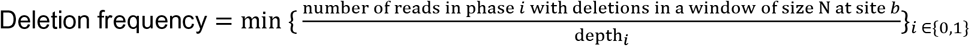
2. *b* is considered an indel candidate site if:

a. Both depth_0_ and depth_1_ are greater than a specified depth threshold
b. Either insertion frequency is greater than a specified insertion frequency threshold or the deletion frequency is greater than a specified deletion frequency threshold.

Thresholds for alternative allele frequency, insertion frequency, deletion frequency and read depths can be specified by the user depending on coverage and base calling error rate of the genome sequencing data. We slide two windows along the reference to calculate what fraction of reads in that window contain an insertion or deletion. The first window uses a larger window size (default is 10) to calculate how many reads contain an insertion or deletion longer than or equal to 3bp, whereas the second window uses a smaller window size (default is 4) to calculate how many reads contain an insertion or deletion shorter than or equal to 10bp. The reason for allowing only indels longer than or equal to 3bp in the large window is to prevent sequencing errors that give rise to several small 2bp indels from producing falsely high indel frequency.

##### Pileup image generation

Input image of indel candidate site is generated using the procedure below as shown in Figure 2. For an indel candidate site *b*:

1. Denote by *S*_*all*_, *S*_*phase*1_ and *S*_*phase*2_ the set of all reads, reads in a phase and reads in the other phase at site *b*, respectively.
2. Let *seq*_*ref*_ be the reference sequence of length 160bp starting at site *b*
3. For each set *S* ∈ {*S*_*all*_, *S*_*phase*1_, *S*_*phase*2_ }, do the following:

a. For each read *r* ∈ *S*, let *seq*_*r*_ be the 160bp long subsequence of the read starting at the site *b* (for PacBio datasets, we use reference sequence and alignment sequences of length 260bp).
b. Use MUSCLE to carry out multiple sequence alignment of the following set of sequences {*seq*_*ref*_ } ∪ {*seq*_*r*_}_*r*∈*S*_ as shown in Figure 2 a).
c. Let {*seq*′_*ref*_ } ∪ {*seq*′_*r*_}_*r*∈*S*_ be the realigned sequences, where *seq*′_*ref*_ denotes the realigned reference sequence, and *seq*′_*r*_ denotes realignment of sequence *seq*_*r*_. We truncate all sequences at the length 128 from the end.
d. For *B* ∈ {*A, G, T, C*, −} and 1 ≤ *p* ≤ 128, calculate

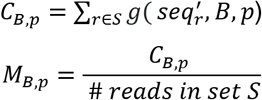
where *g*(*seq*_r_^′^, *B*, *p*) returns 1 if the base at index *p* of *seq_r_*^′^ is *B* and 0 otherwise. Figure 2 c) shows raw counts *C*_*B*,*p*_ for each symbol.
e. Let *M* be the 5 × 128 matrix with entries *M*_*B*,*p*_ as shown in Figure 2 d).
f. Construct a 5 × 128 matrix *Q*, with entries 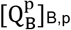, where 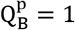 if *seq*_ref_^′^ has symbol *B* at index *p* and 0 otherwise as shown in Figure 2 f). Both matrices *M* and *Q* have first dimension corresponding to the symbols {*A, G, T, C*, −}, and second dimension corresponding to pileup columns of realigned sequences.
g. Construct a 5 × 128 × 2 matrix *Mat*_*S*_ whose first channel is the matrix *M* − *Q* as shown in Figure 2 e) and the second channel is the matrix *Q*.
4. Concatenate the three matrices *Mat*_*S_all_*_, *Mat*_*S*_*phase*1__ and *Mat*_*S*_*phase*2__ together to get a 15 × 128 × 2 matrix as input to convolutional neural network.

##### Deep learning prediction

In NanoCaller, we present another convolutional neural networks [37] for indel calling, as shown in Figure 2. This neural network has a similar structure as SNP calling, and the difference is the fully connected network: for indel model, the first fully connected layer is followed by two fully connected hidden layers to produce probability estimates for each of the four zygosity cases at the candidate site: homozygous-reference, homozygous-alternative, heterozygous-reference and heterozygous-alternative (i.e. heterozygous with no reference allele).

##### Indel sequence determination

After that, NanoCaller calculates the probabilities for four cases of zygosities: homozygous-reference, homozygous-alternative, heterozygous-reference and heterozygous-alternative. No variant call is made if the homozygous-reference label has the highest probability. If homozygous-alternative label has the highest probability, we determine consensus sequence from the multiple sequence alignment of *S*_*all*_, and align it against reference sequence at the candidate site using BioPython’s pairwise2 global alignment algorithm with affine gap penalty. Alternative allele is inferred from the indel of the pairwise alignment of the two sequences. In case either of the heterozygous predictions has the highest probability, we use *S*_*phase*1_ and *S*_*phase*2_ to determine consensus sequences for each phase separately and align them against reference sequence. Indel calls from both phases are combined to make a final phased indel call at the candidate site. Please note that NanoCaller does not filter any indel prediction based on predicted indel length, but 50bp can be the threshold of predicted indel definition since majority of predicted indels is <50bp.

#### Training and testing

For SNP calling, we have trained five convolutional neural network models on two different genomes that users can choose from: ONT-HG001 (trained on HG001 ONT reads basecalled with Guppy 4.2.2), ONT- HG002 (trained on HG002 ONT reads basecalled with Guppy 4.2.2), CCS-HG001 (trained on HG001 CCS reads with 11-20kb library size), CCS-HG002 (trained on HG002 CCS reads with 15-20kb library size) and CLR-HG002 (trained on HG002 PacBio CLR dataset). The first four datasets have both SNP and indel models, whereas CLR-HG002 only has SNP model. All training sequencing datasets were aligned to GRCh38, with only chromosomes 1-22 used for training. We used v3.3.2 GIAB’s benchmark variants for training all HG001 models and v4.2.1 benchmark variants for all HG002 models. Please refer to supplementary materials for the performance of more models, such as NanoCaller models trained on ONT reads of HG001 and HG002 basecalled by older Guppy versions, models trained using v3.3.2 benchmark variants for HG002, and models trained on chromosomes 1-21 of Nanopore R10.3 flowcell reads and Bonito basecalled dataset of HG002.

In NanoCaller, the SNP and indel models have 137,678 parameters in total, a significantly lower number than Clair [24](2,377,818) Clairvoyante[23] (1,631,496). All parameters in NanoCaller are initiated by Xavier’s method [39]. Each model was trained for 100 epochs, using a learning rate of 1e-3 and 1e-4 for SNP and indel models respectively. We also applied L2-norm regularization, with coefficients 1e-3 and 1e-5 for SNP and indel models respectively, to prevent overfitting of the model.

To use NanoCaller on a test genome, it is reasonable that the test genome has different coverage as the genome used for training NanoCaller. To reduce the bias caused by different coverages, after generating pileup images for SNP calling, NanoCaller by default scales the raw counts of bases in pileup images to adjust for the difference between coverages of the testing genome and the genome used for training of the model selected by user, i.e we replace the counts *C*_*B*,*p*_ shown in Figure 1 (b) by

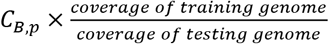

#### Performance measurement

The performance of SNP/indel calling by a variant caller is evaluted against the benchmark variant tests. Several measurements of performance evaluation are used, such as precision (p), recall (r) and F1 score as defined below.

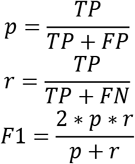

where *TP* is the number of benchmark variants correctly predicted by a variant caller, and *FP* is the number of miscalled variants which are not in benchmark variant sets, *FN* is the number of benchmark variants which cannot be called by a variant caller. *F*1 is the weighted average of *p* and *r*, a harmonic measurement of precision and recall. The range of the three measurements is [0, 1]: the larger, the better.

#### Sanger validation of selected sites on HG002

To further demonstrate the performance of NanoCaller and other variant callers, we select 17 genomic regions whose SNPs/indels are not in the GIAB ground truth calls (version 3.3.2), and conduct Sanger sequencing for them on HG002. Firstly, we design PCR primers within ∼400 bp of a select site of interest and then use a high-fidelity PCR enzyme (PrimeSTAR GXL DNA Polymerase, TaKaRa) to amplify each of the target selected repeat regions. The PCR products are purified using AMPure XP beads and sequenced by Sanger sequencing. We then decipher two sequences from Sanger results for variant analysis. The data and deciphered sequences are in the supplementary files. Please note that more than 17 variant sites are detected in the Sanger results, because each PCR region can contain 1+ variants.

## Acknowledgements

The authors would like to thank members of the Wang lab for valuable comments and feedback. We would like to thank GIAB and nanopore-wgs-consortium for providing the sequencing data sets and the gold standard variant call data for use in our evaluation. We would like to thank PrecisionFDA team for organizing the variant calling Truth Challenge on difficulty-to-map regions and for scoring the submissions. Finally, we would like to thank two anonymous reviewers in offering insightful comments and suggestions on improving NanoCaller. This study is in part supported by NIH/NIGMS grant GM132713 to KW.

## Author contributions

UA and QL developed the computational method and drafted the manuscript. UA implemented the software tool and evaluated its performance. LF conducted wet-lab experiments of Sanger sequencing of candidate variants. KW conceived the study, advised on model design and guided implementation/evaluation. All authors read, revised, and approved the manuscript.

## Statistics of long-reads datasets

**Table S1.** Whole genome statistics of datasets on five human genomes by Nanopore and PacBio sequencing. Each genome is aligned to the GRCh38 reference genome, and only the mapped reads are used to calculate the statistics. Total number of bases is calculated as the sum of the length of all mapped reads, and the coverage is defined as the number of mapped bases divided by the reference genome length.’Mean’ and ‘Median’ represent the mean and median of read length in long reads for a genome. ONT HG001 reads were basecalled by Guppy 2.3.8, ONT HG002-4 reads were basecalled by Guppy 3.6, and HX1 reads were basecalled with Albacore. HG001 CCS (11kb library size) reads were obtained GIAB’s database, whereas HG002-4 CCS (15kb library size) reads were obtained from precisionFDA truth challenge V2.

| Platform | Genome | # reads | # bases | N50 | Mean | Median | Coverage |
| --- | --- | --- | --- | --- | --- | --- | --- |
| ONT | HG001 | 15,666,888 | 132.9 Gb | 13,630 | 8,485 | 5,387 | 43X |
| ONT | HG002 | 7,788,089 | 144.2 Gb | 52,549 | 18,514 | 6,672 | 49X |
| ONT | HG003 | 13,601,716 | 249.4 Gb | 46,163 | 18,335 | 8,168 | 85X |
| ONT | HG004 | 12,530,839 | 251.3 Gb | 49,563 | 20,051 | 8,371 | 88X |
| ONT | HX1 | 20,497,769 | 271.8 Gb | 22,273 | 13,261 | 10,123 | 88X |
| PacBio (CLR) | HG001 | 23,808,790 | 106.3 Gb | 6,309 | 4,466 | 3,180 | 37X |
| PacBio (CLR) | HG002 | 21,511,145 | 179.0 Gb | 11,264 | 8,323 | 7,500 | 58X |
| PacBio (CLR) | HG003 | 10,564,465 | 85.3 Gb | 10,943 | 8,078 | 7,251 | 28X |
| PacBio (CLR) | HG004 | 10,369,228 | 83.4 Gb | 10,869 | 8,040 | 7,545 | 27X |
| PacBio (CCS) | HG001 | 8,487,716 | 84.6 Gb | 10,004 | 9,963 | 9,973 | 29X |
| PacBio (CCS) | HG002 | 7,979,372 | 102.6 Gb | 12,883 | 12,855 | 12,781 | 36X |
| PacBio (CCS) | HG003 | 6,872,414 | 100.5 Gb | 14,761 | 14,627 | 14,627 | 35X |
| PacBio (CCS) | HG004 | 6,578,333 | 98.9 Gb | 15,100 | 15,031 | 14,949 | 34X |

## Performance of NanoCaller and other variant callers on old Nanopore datasets of the Ashkenazim trio

From Table S2-S5, three NanoCaller models are evaluated: NanoCaller SNP models NanoCaller1 (trained on ONT HG001 Guppy 2.3.8) and NanoCaller2 (trained on ONT HG002 Guppy 2.3.4), and NanoCaller1 indel model trained on ONT HG001 Guppy 2.3.8 reads. All models are trained using GIAB v3.3.2 benchmark variants. From Table S2-S5, the testing is done on HG002 ONT reads basecalled with Guppy 2.3.4, and HG003-4 ONT reads basecalled with Guppy 3.2.

**Table S2.**
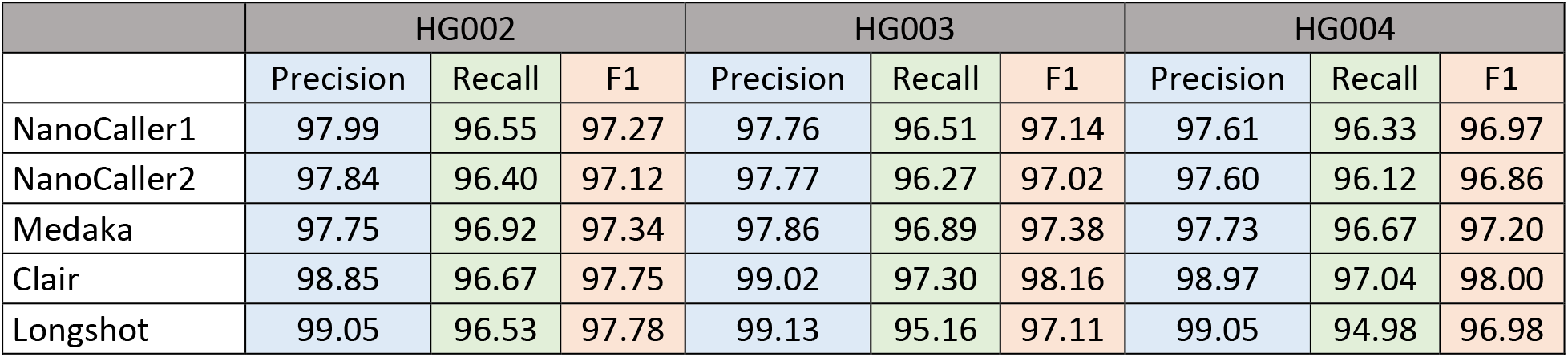
Performances of SNP predictions of NanoCaller1 and NanoCaller2 SNP models, along with existing variant callers, on ONT data of the Ashkenazim trio, evaluated against v3.3.2 of GIAB benchmark variants.

**Table S3.** Performances of SNP predictions of NanoCaller1 and NanoCaller2 SNP models, along with existing variant callers, on ONT data of the Ashkenazim trio, evaluated against v4.2 of GIAB benchmark variants.

|  | HG002 |  |  | HG003 |  |  | HG004 |  |  |
| --- | --- | --- | --- | --- | --- | --- | --- | --- | --- |
|  | Precision | Recall | F1 | Precision | Recall | F1 | Precision | Recall | F1 |
| NanoCaller1 | 98.90 | 96.77 | 97.83 | 99.03 | 96.77 | 97.89 | 98.96 | 96.50 | 97.72 |
| NanoCaller2 | 98.79 | 96.63 | 97.70 | 98.87 | 96.75 | 97.80 | 98.85 | 96.47 | 97.65 |
| Medaka | 98.81 | 96.98 | 97.89 | 98.95 | 97.04 | 97.99 | 99.02 | 96.84 | 97.92 |
| Clair | 99.01 | 96.61 | 97.80 | 99.07 | 97.51 | 98.29 | 99.09 | 97.24 | 98.16 |
| Longshot | 98.97 | 95.89 | 97.41 | 99.11 | 94.60 | 96.81 | 99.13 | 94.32 | 96.67 |

**Table S4.** Performances of indel predictions in non-homopolymer regions of NanoCaller, along with existing variant callers, on ONT data of the Ashkenazim trio, evaluated against v3.3.2 of GIAB benchmark variants.

|  | HG002 |  |  | HG003 |  |  | HG004 |  |  |
| --- | --- | --- | --- | --- | --- | --- | --- | --- | --- |
|  | Precision | Recall | F1 | Precision | Recall | F1 | Precision | Recall | F1 |
| NanoCaller1 | 77.37 | 59.65 | 67.37 | 75.90 | 60.09 | 67.08 | 73.56 | 58.30 | 65.05 |
| Medaka | 76.56 | 63.64 | 69.51 | 78.27 | 65.98 | 71.61 | 77.13 | 63.90 | 69.90 |
| Clair | 84.92 | 55.26 | 66.96 | 85.39 | 58.53 | 69.46 | 85.76 | 56.48 | 68.11 |

**Table S5.**
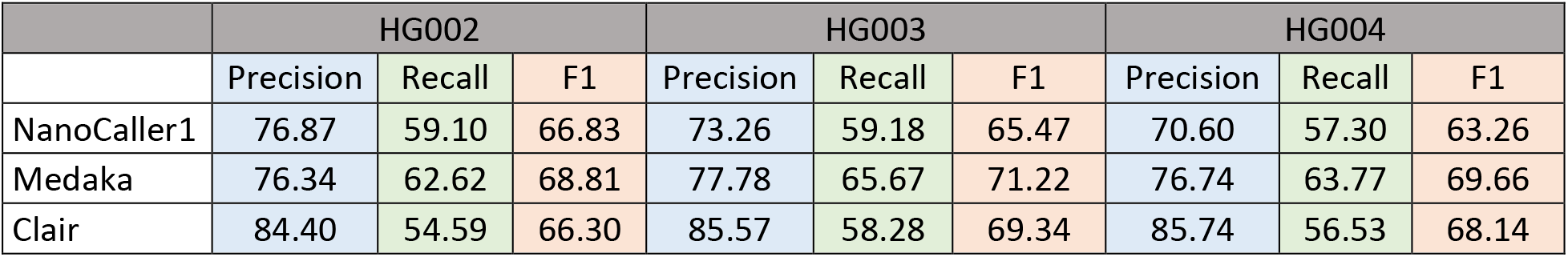
Performances of indel predictions in non-homopolymer regions of NanoCaller, along with existing variant callers, on ONT data of the Ashkenazim trio, evaluated against v4.2 of GIAB benchmark variants.

## SNP performance of NanoCaller and other variant callers in difficult-to-map regions on Nanopore reads of the Ashkenazim trio

BED file sources of different difficult-to-map regions with respect to GRCh38 reference genome:

1. All difficult regions.
2. Source: ftp://ftp-trace.ncbi.nlm.nih.gov/ReferenceSamples/giab/release/genome-stratifications/v2.0/GRCh38/union/GRCh38_alldifficultregions.bed.gz
3. Low mappability regions in Table S8.
4. Source: ftp://ftp-trace.ncbi.nlm.nih.gov/ReferenceSamples/giab/release/genome-stratifications/v2.0/GRCh38/mappability/GRCh38_lowmappabilityall.bed.gz
5. Segmental duplications in Table S9.
6. Source: ftp://ftp-trace.ncbi.nlm.nih.gov/ReferenceSamples/giab/release/genome-stratifications/v2.0/GRCh38/SegmentalDuplications/GRCh38_segdups.bed.gz
7. Tandem and homopolymer repeats (perfect homopolymers longer than 6bp and imperfect homopolymers longer than 10bp) in Table S10.
8. Source: ftp://ftp-trace.ncbi.nlm.nih.gov/ReferenceSamples/giab/release/genome-stratifications/v2.0/GRCh38/LowComplexity/GRCh38_AllTandemRepeatsandHomopolymers_slop5.bed.gz
9. <u>Perfect</u> homopolymers longer than 6bp and imperfect homopolymers longer than 10bp. Source: https://ftp-trace.ncbi.nlm.nih.gov/ReferenceSamples/giab/release/genome-stratifications/v2.0/GRCh38/LowComplexity/GRCh38_AllHomopolymers_gt6bp_imperfectgt10bp_slop5.bed.gz
10. Perfect homopolymers of lengths 4-6bp.
11. Source: https://ftp-trace.ncbi.nlm.nih.gov/ReferenceSamples/giab/release/genome-stratifications/v2.0/GRCh38/LowComplexity/GRCh38_SimpleRepeat_homopolymer_4to6_slop5.bed.gz
12. Table S11.
13. Source: ftp://ftp-trace.ncbi.nlm.nih.gov/ReferenceSamples/giab/release/genome-stratifications/v2.0/GRCh38/OtherDifficult/GRCh38_MHC.bed.gz

The details of these files are provided here: https://ftp-trace.ncbi.nlm.nih.gov/ReferenceSamples/giab/release/genome-stratifications/v2.0/GRCh38/LowComplexity/v2.0-GRCh38-LowComplexity-README.txt

### Easy genomic regions

Easy genomic regions are also used during performance comparison and are defined as the complement of “all difficult-to-map” regions (GRCh38_alldifficultregions.bed). Easy genomic regions are obtained by removing “all difficult-to-map” regions from each genome’s high-confidence intervals.

### Various difficult-to-map regions

The following difficult-to-map regions are used for performance evaluation and comparison: 1) “all difficult-to-map” regions, 2) low mappability regions, 3) segmental duplications, 4) tandem and homopolymer repeats, and 5) MHC. For each of these categories, we get the evaluation regions by intersecting the corresponding BED file with high-confidence intervals for each genome using BEDtools.

### Definition of homopolymer and non-homopolymer regions for indel performance evaluation

GIAB genome stratification BED is used to obtain homopolymer regions and non-homopolymer regions for evaluating indel performance. Homopolymer regions are obtained by combining GRCh38_AllHomopolymers_gt6bp_imperfectgt10bp_slop5.bed and GRCh38_SimpleRepeat_homopolymer_4to6_slop5.bed BED files. These first BED file contains intervals with perfect homopolymer regions longer than 6bp and imperfect homopolymers longer than 10bp (defined by GIAB as a ‘single base was repeated >10bp except for a 1bp interruption by a different base’, e.g. AAAATAAAAAA). The second BED file contains all intervals of homopolymer regions of lengths between 4 to 6bp. *It is important to note that GRCh38_AllTandemRepeatsandHomopolymers_slop5.bed contains homopolymer regions from the first BED file only, and does not contain homopolymers of lengths 4-6bp from the second BED file*.

Meanwhile, non-homopolymer regions are obtained by subtracting the homopolymer regions from GIAB high-confidence intervals for each genome.

**Table S6.** Statistics of number ground truth variants in GIAB v4.2 benchmark of the Ashkenazim trio within various difficult genomic regions identified in GIAB genome stratification v2.0. Numbers shown below are for SNPs, with the exception of MHC region for which both SNP and indel counts are shown.

| Region | HG002 | HG003 | HG004 |
| --- | --- | --- | --- |
| All Difficult Regions | 628,157 | 611,270 | 619,719 |
| Tandem and Homopolymer Repeats | 182,901 | 166,398 | 166,807 |
| Segmental Duplications | 121,179 | 122,393 | 125,711 |
| Low Mappability regions | 192,749 | 193,153 | 195,332 |
| MHC (SNPs) | 20,139 | 19,730 | 20,576 |
| MHC (indels) | 1667 | 1655 | 1738 |

Please note that in Tables S7-S11, the performance of NanoCaller1 model is tested on precisionFDA challenge HG002-4 ONT reads basecalled with Guppy3.6.

**Table S7.** Performances of SNP predictions in all difficult genomic regions by NanoCaller1 SNP models, along with existing variant callers on ONT data, using 4.2 benchmark variants.

|  |  | NanoCaller1 | Medaka | Clair | Longshot |
| --- | --- | --- | --- | --- | --- |
|  | Threshold | 144 | 3 | 370 | 118 |
| Precision | HG002 | 96.16 | 94.74 | 94.68 | 97.48 |
|  | HG003 | 97.30 | 96.14 | 96.47 | 96.84 |
|  | HG004 | 97.23 | 96.41 | 96.23 | 96.87 |
| Recall | HG002 | 95.04 | 95.48 | 95.33 | 91.12 |
|  | HG003 | 96.27 | 96.54 | 96.11 | 92.34 |
|  | HG004 | 96.14 | 96.12 | 96.22 | 92.06 |
| F1 | HG002 | **95.60** | 95.11 | 95.01 | 94.19 |
|  | HG003 | **96.78** | 96.34 | 96.29 | 94.53 |
|  | HG004 | **96.68** | 96.27 | 96.22 | 94.40 |

**Table S8.**
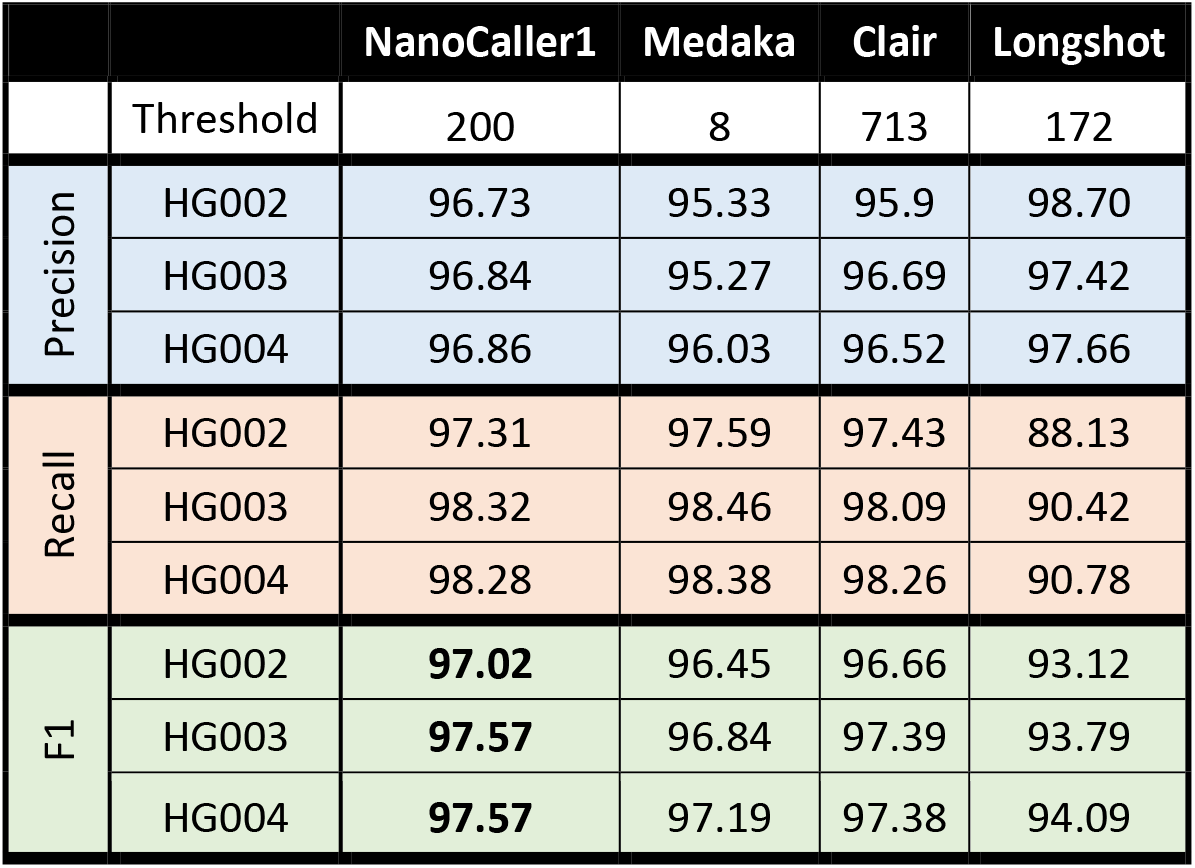
Performances of SNP predictions in low mappability regions by NanoCaller1 SNP models, along with existing variant callers on ONT data, using 4.2 benchmark variants.

**Table S9.** Performances of SNP predictions in segmental duplication regions by NanoCaller1 SNP models, along with existing variant callers on ONT data, using 4.2 benchmark variants.

|  |  | NanoCaller1 | Medaka | Clair | Longshot |
| --- | --- | --- | --- | --- | --- |
|  | Threshold | 228 | 11 | 768 | 186 |
| Precision | HG002 | 94.19 | 90.79 | 93.07 | 97.99 |
|  | HG003 | 94.43 | 92.2 | 94.55 | 95.86 |
|  | HG004 | 94.58 | 93.64 | 94.3 | 96.30 |
| Recall | HG002 | 95.64 | 96.54 | 96.23 | 84.99 |
|  | HG003 | 97.19 | 97.46 | 97.16 | 88.9 |
|  | HG004 | 97.17 | 97.31 | 97.27 | 89.27 |
| F1 | HG002 | **94.91** | 93.58 | 94.62 | 91.03 |
|  | HG003 | 95.79 | 94.75 | **95.84** | 92.25 |
|  | HG004 | **95.86** | 95.44 | 95.76 | 92.66 |

**Table S10.** Performances of SNP predictions in tandem and homopolymer repeat regions by NanoCaller1 SNP models, along with existing variant callers on ONT data, using 4.2 benchmark variants.

|  |  | NanoCaller1 | Medaka | Clair | Longshot |
| --- | --- | --- | --- | --- | --- |
|  | Threshold | 93 | 2 | 300 | 115 |
| Precision | HG002 | 93.20 | 90.09 | 90.48 | 93.96 |
|  | HG003 | 95.48 | 93.49 | 93.38 | 93.23 |
|  | HG004 | 95.20 | 92.63 | 92.85 | 92.70 |
| Recall | HG002 | 88.53 | 88.30 | 89.05 | 89.67 |
|  | HG003 | 90.30 | 90.14 | 90.29 | 92.31 |
|  | HG004 | 89.78 | 89.57 | 90.01 | 91.81 |
| F1 | HG002 | 90.81 | 89.19 | 89.76 | **91.76** |
|  | HG003 | **92.82** | 91.78 | 91.81 | 92.76 |
|  | HG004 | **92.41** | 91.07 | 91.41 | 92.25 |

**Table S11.** Performances of SNPs, indels and overall variants predictions in MHC by NanoCaller1 SNP models, along with existing variant callers on ONT data, using 4.2 benchmark variants. Overall variant performance is calculated by combining the SNPs and indels performances. The ensemble variant calls are created by combining NanoCaller1, Medaka and Clair variant calls and was submitted to precisionFDA Truth Challenge V2.

| SNPs Performance |  |  |  |  |  |  |
| --- | --- | --- | --- | --- | --- | --- |
|  |  | Ensemble | NanoCaller1 | Medaka | Clair | Longshot |
|  | Threshold | -- | 145 | 0 | 272 | 15 |
| Precision | HG002 | 99.44 | 98.88 | 99.02 | 97.88 | 98.77 |
|  | HG003 | 99.58 | 99.37 | 99.61 | 98.76 | 99.10 |
|  | HG004 | 99.51 | 99.18 | 97.92 | 98.77 | 99.00 |
| Recall | HG002 | 98.50 | 98.18 | 95.34 | 97.23 | 52.09 |
|  | HG003 | 99.11 | 99.09 | 99.23 | 98.60 | 57.24 |
|  | HG004 | 98.84 | 98.97 | 90.93 | 98.40 | 51.05 |
| F1 | HG002 | 98.97 | **98.53** | 97.15 | 97.55 | 68.21 |
|  | HG003 | 99.34 | 99.23 | **99.42** | 98.68 | 72.57 |
|  | HG004 | 99.17 | **99.07** | 94.29 | 98.59 | 67.37 |
| Indels Performance |  |  |  |  |  |  |
|  |  | Ensemble | NanoCaller1 | Medaka | Clair | Longshot |
|  | Threshold | -- | 205 | 19 | 509 | -- |
| Precision | HG002 | 79.40 | 72.77 | 74.02 | 81.00 | -- |
|  | HG003 | 80.23 | 75.02 | 80.20 | 78.99 | -- |
|  | HG004 | 76.39 | 74.59 | 75.80 | 81.32 | -- |
| Recall | HG002 | 57.21 | 59.47 | 64.78 | 57.31 | -- |
|  | HG003 | 66.18 | 61.89 | 78.47 | 61.59 | -- |
|  | HG004 | 64.00 | 64.55 | 68.84 | 60.86 | -- |
| F1 | HG002 | 66.51 | 65.45 | **69.09** | 67.13 | -- |
|  | HG003 | 72.53 | 67.82 | **79.33** | 69.21 | -- |
|  | HG004 | 69.64 | 69.21 | **72.15** | 69.62 | -- |
| Overall Variants Performance |  |  |  |  |  |  |
|  |  | Ensemble | NanoCaller1 | Medaka | Clair | Longshot |
| Precision | HG002 | 98.30 | 97.20 | 97.19 | 96.92 | -- |
|  | HG003 | 98.30 | 97.78 | 98.06 | 97.51 | -- |
|  | HG004 | 97.94 | 97.48 | 96.16 | 97.70 | -- |
| Recall | HG002 | 95.34 | 95.22 | 92.99 | 94.18 | -- |
|  | HG003 | 96.56 | 96.21 | 97.62 | 95.73 | -- |
|  | HG004 | 96.12 | 96.28 | 89.20 | 95.46 | -- |
| F1 | HG002 | 96.79 | **96.20** | 95.05 | 95.53 | -- |
|  | HG003 | 97.42 | 96.99 | **97.84** | 96.61 | -- |
|  | HG004 | 97.02 | **96.87** | 92.55 | 96.57 | -- |

## Novel variants validated by Sanger sequencing

**Table S12.**
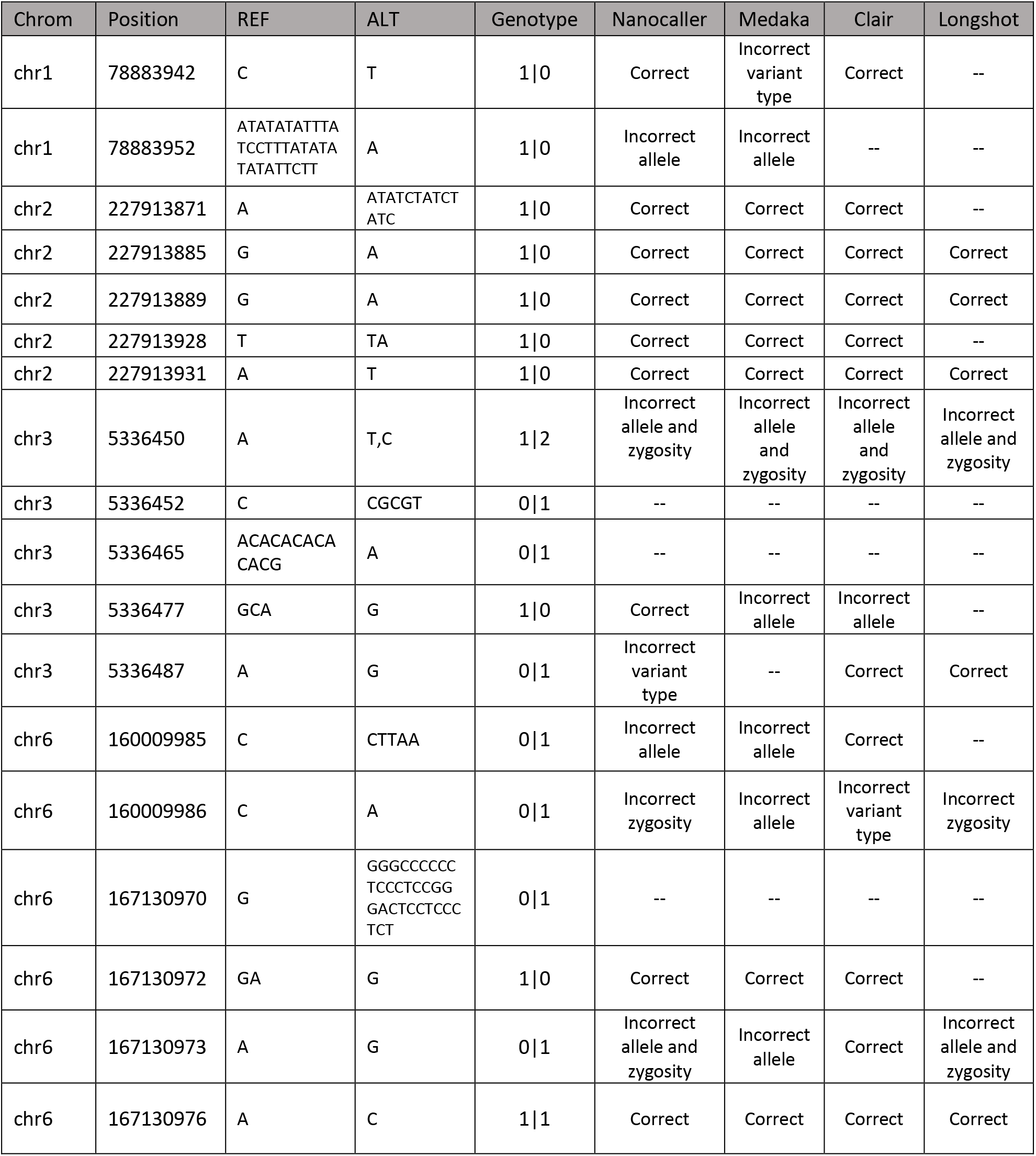
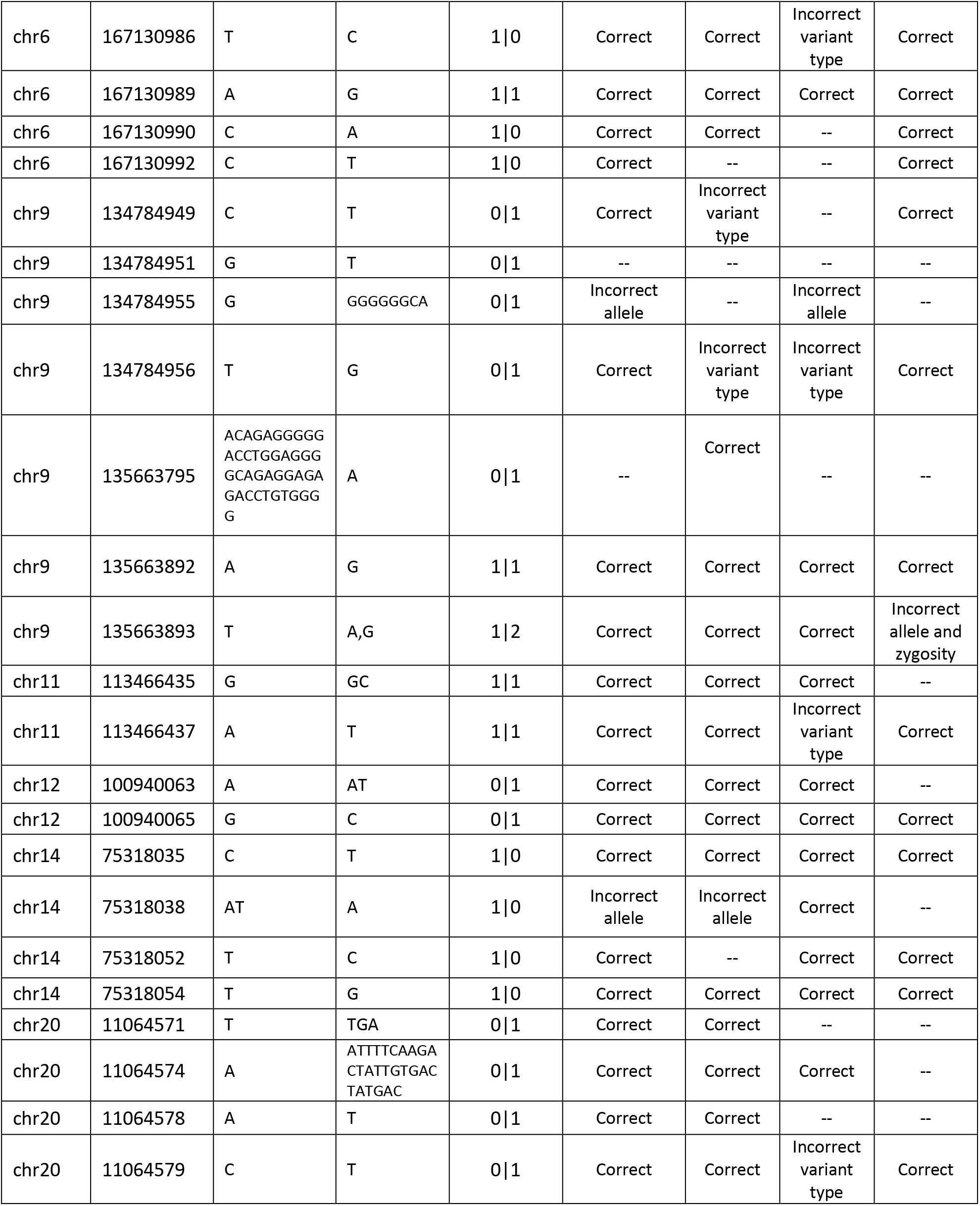
Sanger validated variants, and performance of predictions in HG002 genome by various variant callers using HG002 ONT reads basecalled with Guppy 3.6. These variants are missing in GIAB v3.3.2 of HG002.\

## Quality score thresholds for filtering variant calls

Precision/recall/F1 statistics for each variant caller are investigated with respect to the recommended quality score, or by taking average over quality scores giving highest F1-score when evaluated by RTG’s *vcfeval*. Here the quality thresholds for different Nanopore models on various datasets can show how the quality of reads, depth and quality of ground truth variants affect the optimal performance evaluation. In summary, the range of SNP quality scores is 30-999, but **Table S13** shows that SNP F1- scores of both NanoCaller1 and NanoCaller2 SNP models are very resilient to small changes on quality score thresholds. For NanoCaller1 and NanoCaller3 SNP models, the best performance on PacBio CCS reads can be obtained without any specialized threshold (which corresponds to a quality score threshold of 30). For PacBio CLR reads, genome coverage significantly effects SNP calling performance, thus we choose different thresholds for HG001/HG002 and HG003/HG004, as shown in **Table S14**.

Please note in Tables S13-S16, NanoCaller models are tested on precisionFDA challenge HG002-4 ONT reads basecalled with Guppy3.6, HG001 ONT reads basecalled with Guppy 2.3.8, HX1 ONT reads basecalled with Albacore, HG001 CCS (11kb library size) reads obtained GIAB’s database, and HG002-4 CCS (15kb library size) reads downloaded from precisionFDA truth challenge V2.

**Table S13.** F1 scores of NanoCaller1 and NanoCaller2 SNP models on ONT datasets. 162 and 78 are recommended as quality thresholds for NanoCaller1 and NanoCaller2 models, respectively. The column ‘Quality Score Range’ shows the range of quality scores that give the optimal SNP F1-score shown in the ‘Best F1’ column. Third colum under each model shows the F1-score with the recommended threshold of quality scores.

|  | NanoCaller1 |  |  | NanoCaller2 |  |  |
| --- | --- | --- | --- | --- | --- | --- |
|  | Quality Score Range | Best F1 | F1 with Quality Score=162 | Quality Score Range | Best F1 | F1 with Quality Score=78 |
| HG001 | 120-120 | 95.54 | 95.40 | 65-70 | 94.93 | 94.89 |
| HG002 | 179-201 | 98.03 | 97.99 | 99-117 | 98.03 | 97.97 |
| HG003 | 157-185 | 98.88 | 98.88 | 62-85 | 98.88 | 98.88 |
| HG004 | 166-191 | 98.78 | 98.77 | 71-87 | 98.82 | 98.82 |
| HX1 | 191-194 | 90.52 | 90.28 | 93-97 | 91.07 | 90.94 |

**Table S14.** F1 scores of NanoCaller1 and NanoCaller3 SNP models on CLR datasets. The thresholds of quality scores are recommended for higher coverage genomes HG001/HG002 and lower coverage genomes HG003/HG004. The column ‘Quality Score Range’ shows the range of quality scores that give the optimal SNP F1-score shown in the ‘Best F1’ column. Third and fourth columns under each model shows the F1-score with the recommended thresholds of quality scores.

|  | NanoCaller1 |  |  |  | NanoCaller3 |  |  |  |
| --- | --- | --- | --- | --- | --- | --- | --- | --- |
|  | Quality Score Range | Best F1 | Reported Quality Cut-off | F1 score using reported Cut-off | Quality Score Range | Best F1 | Reported Quality Cut-off | F1 score using reported Cut-off |
| HG001 | 30 | 93.97 | None | 93.97 | 30-37 | 94.95 | 55 | 94.90 |
| HG002 | 32-34 | 98.26 | None | 98.26 | 79-111 | 98.29 | 55 | 98.24 |
| HG003 | 108-114 | 94.33 | 110 | 94.33 | 202-210 | 93.53 | 210 | 93.53 |
| HG004 | 118-132 | 93.49 | 110 | 93.47 | 209-218 | 92.61 | 210 | 92.59 |

**Table S15.** F1 scores of NanoCaller ONT and PacBio indel models. 44 and 25 are recommended as quality thresholds for ONT and PacBio models, respectively. The column ‘Quality Score Range’ shows the range of quality scores that give the optimal indel F1-score shown in the ‘Best F1’ column. Third colum under each model shows the F1-score with the recommended thresholds of quality scores.

|  | ONT model |  |  | PacBio model |  |  |
| --- | --- | --- | --- | --- | --- | --- |
|  | Quality Score Range | Best F1 | F1 with Quality Score=144 | Quality Score Range | Best F1 | F1 with Quality Score=25 |
| HG001 | 243-247 | 56.79 | 53.30 | 2-8 | 94.16 | 94.09 |
| HG002 | 145-152 | 79.75 | 79.74 | 2-5 | 94.52 | 94.40 |
| HG003 | 101-101 | 81.31 | 81.23 | 52-55 | 93.68 | 93.53 |
| HG004 | 101-104 | 79.76 | 79.71 | 56-64 | 93.61 | 93.37 |

**Table S16.** Quality thresholds used for different variant callers for different sequencing technologies. For Clair and PacBio, the developer recommended thresholds are used, whereas DeepVariant developers do not recommend using any threshold. For Medaka, the average of best quality scores for SNPs over five ONT genomes are calculated, and used as final SNP quality cut-off (the same procedure for the quality thresholds of indels). For WhatsHap, best results are obtained without any quality cut-off as well.

| Variant Caller | Sequencing Type | Threshold |
| --- | --- | --- |
| Medaka | ONT | 2 (SNPs), 23 (indels) |
| Clair | ONT | 748 |
|  | PacBio | 113 |
| Longshot | ONT | 65 |
|  | PacBio | 50 |
| WhatsHap | ONT, PacBio | None |
| DeepVariant | PacBio | None |

## Runtime Comparisons

**Table S17.**
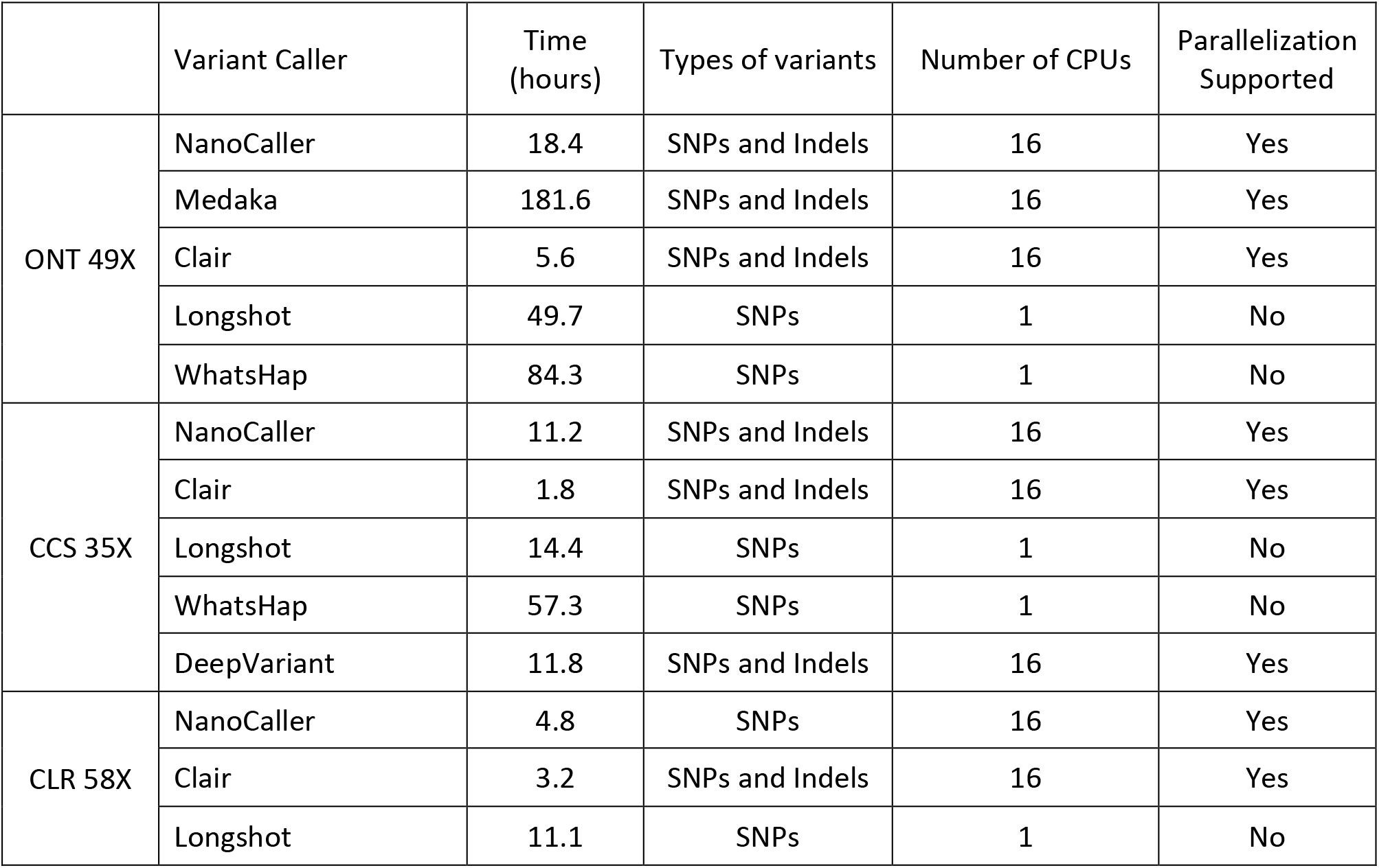
Wall-clock runtimes for various variant callers using Intel Xeon CPU E5-2683 v4 @ 2.10GHz. For variant callers that support parallelization, 16 CPUs are used. The runtime evaluation is tested on 49X HG002 ONT reads basecalled with Guppy 3.6, 35X HG002 PacBio CCS reads (15kb library size) from precisionFDA challenge, and 58X HG002 PacBio CLR reads.

## Selection of nearby potentially heterozygous sites for SNP calling feature generation

Please note that in Tables S18-S23, NanoCaller models are evaluated on precisionFDA challenge HG002-4 ONT reads basecalled with Guppy3.6, HG002-4 CCS (15kb library size) datasets from precisionFDA truth challenge V2, HG002-4 PacBio CLR reads.

**Table S18.**
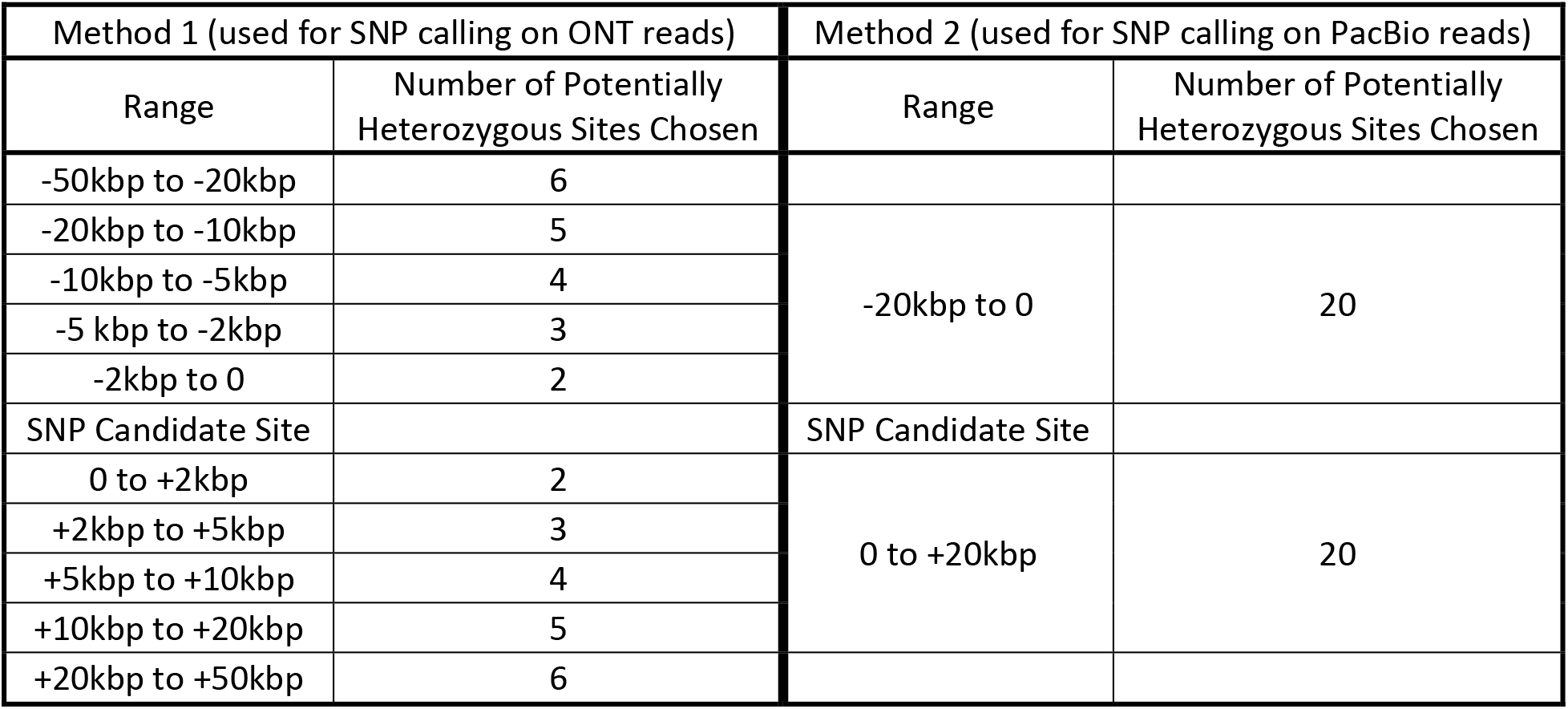
Number of potentially heterozygous Sites chosen for each candidate site under two methods. Method 1 is used for SNP calling on ONT reads and selects the given number of sites from each range given. Method 2 is used for SNP calling on PacBio reads. In each range for either method, the specified number of sites closest to the candidate site are selected, because they share more reads with the candidate site. The design of these two methods is motivated by the difference in read length distribution of ONT and PacBio reads, shown in Figure S2. An illustration of the two methods is shown in Figure S3.

**Table S19.** Performance of NanoCaller SNP calling on HG004 ONT, CCS and CLR reads with various thresholds used to define heterozygous SNPs. The best F1-scores with their own thresholds on v4.2 benchmark variants are shown here.

| Oxford Nanopore Reads |  |  |  |
| --- | --- | --- | --- |
| Threshold | Precision | Recall | F1 |
| 0.30-0.60 | 0.9846 | 0.9843 | 0.9845 |
| 0.30-0.70 | 0.9840 | 0.9833 | 0.9837 |
| 0.30-0.80 | 0.9814 | 0.9797 | 0.9806 |
| 0.35-0.65 | 0.9878 | 0.9857 | 0.9868 |
| 0.40-0.60 | 0.9888 | 0.9867 | **0.9878** |
| 0.40-0.70 | 0.9882 | 0.9847 | 0.9865 |
| 0.40-0.80 | 0.9845 | 0.9790 | 0.9818 |
| 0.45-0.55 | 0.9880 | 0.9851 | 0.9866 |
| PacBio CLR Reads |  |  |  |
| Threshold | Precision | Recall | F1 |
| 0.20-0.70 | 0.9527 | 0.8829 | 0.9165 |
| 0.30-0.60 | 0.9595 | 0.9114 | **0.9349** |
| 0.30-0.70 | 0.9559 | 0.8971 | 0.9256 |
| 0.30-0.80 | 0.9410 | 0.8730 | 0.9058 |
| 0.35-0.65 | 0.9534 | 0.8989 | 0.9254 |
| 0.40-0.60 | 0.9457 | 0.8906 | 0.9174 |
| 0.40-0.70 | 0.9409 | 0.8689 | 0.9035 |
| 0.40-0.80 | 0.9215 | 0.8400 | 0.8789 |
| 0.45-0.55 | 0.9228 | 0.8607 | 0.8907 |
| PacBio CCS Reads |  |  |  |
| Threshold | Precision | Recall | F1 |
| 0.30-0.60 | 0.9920 | 0.9944 | 0.9932 |
| 0.30-0.70 | 0.9936 | 0.9963 | **0.9950** |
| 0.30-0.80 | 0.9935 | 0.9967 | 0.9951 |
| 0.35-0.65 | 0.9932 | 0.9950 | 0.9941 |
| 0.40-0.60 | 0.9900 | 0.9899 | 0.9900 |
| 0.40-0.70 | 0.9936 | 0.9950 | 0.9943 |
| 0.40-0.80 | 0.9935 | 0.9957 | 0.9946 |
| 0.45-0.55 | 0.9769 | 0.9702 | 0.9735 |

**Table S20.** Performance of NanoCaller SNP calling on HG004 ONT and CCS reads with various thresholds for minimum number (first column) of heterozygous SNPs required for a a candidate site. The best F1-scores with their own thresholds on v4.2 benchmark variants are shown here.

| **Oxford Nanopore Reads** |  |  |  |
| --- | --- | --- | --- |
| Minimum Number | Precision | Recall | F1 |
| 1 | 98.88 | 98.67 | 98.78 |
| 2 | 98.84 | 98.69 | 98.77 |
| 3 | 98.84 | 98.65 | 98.75 |
| 5 | 98.90 | 98.35 | 98.63 |
| 7 | 98.94 | 97.66 | 98.30 |
| 9 | 98.96 | 96.43 | 97.68 |
| 13 | 98.93 | 92.21 | 95.46 |
| **PacBio CCS** |  |  |  |
| Minimum Number | Precision | Recall | F1 |
| **1** | **99.37** | **99.63** | **99.50** |
| 3 | 99.41 | 95.54 | 97.44 |
| 5 | 99.44 | 90.53 | 94.77 |
| 7 | 99.44 | 87.04 | 92.83 |
| 9 | 99.44 | 84.79 | 91.53 |

**Table S21.** Performance of NanoCaller SNP calling, in whole genome and various difficult genomic regions, for HG002, HG003 and HG004 ONT reads using Method1 and Method2 described in Table S18. Shown are the best F1-scores achieved by NanoCaller1 or NanoCaller2 SNP models with evaluation on v4.2 benchmark variants.

| NanoCaller1 |  |  |  |  |  |  |  |
| --- | --- | --- | --- | --- | --- | --- | --- |
| Genomic Regions | Genomes | Method1 (for ONT reads) |  |  | Method2 (for PacBio reads) |  |  |
|  |  | Precision | Recall | F1 | Precision | Recall | F1 |
| Whole genome | HG002 | 98.23 | 97.82 | 98.03 | 98.19 | 97.44 | 97.82 |
|  | HG003 | 99.00 | 98.76 | 98.88 | 98.98 | 98.51 | 98.75 |
|  | HG004 | 98.88 | 98.67 | 98.78 | 98.80 | 98.45 | 98.63 |
| All difficult regions | HG002 | 96.59 | 94.68 | 95.63 | 96.19 | 94.65 | 95.42 |
|  | HG003 | 97.13 | 96.44 | 96.79 | 97.02 | 96.34 | 96.68 |
|  | HG004 | 97.19 | 96.19 | 96.69 | 97.03 | 96.07 | 96.55 |
| Low Mappability Regions | HG002 | 96.63 | 97.40 | 97.02 | 96.61 | 97.14 | 96.88 |
|  | HG003 | 96.77 | 98.37 | 97.57 | 96.80 | 98.18 | 97.49 |
|  | HG004 | 97.00 | 98.15 | 97.58 | 96.80 | 98.14 | 97.47 |
| Tandem and Homopolymer Repeats | HG002 | 93.87 | 88.04 | 90.87 | 92.98 | 88.27 | 90.57 |
|  | HG003 | 95.32 | 90.47 | 92.84 | 95.04 | 90.33 | 92.63 |
|  | HG004 | 94.99 | 89.99 | 92.43 | 94.83 | 89.66 | 92.18 |
| Segmental Duplications | HG002 | 94.07 | 95.81 | 94.94 | 93.83 | 95.72 | 94.77 |
|  | HG003 | 94.65 | 96.99 | 95.81 | 94.51 | 96.97 | 95.73 |
|  | HG004 | 94.51 | 97.26 | 95.87 | 94.41 | 97.20 | 95.79 |
| MHC | HG002 | 98.89 | 98.18 | 98.54 | 98.83 | 98.12 | 98.48 |
|  | HG003 | 99.36 | 99.11 | 99.24 | 99.39 | 98.88 | 99.14 |
|  | HG004 | 99.21 | 98.96 | 99.09 | 99.21 | 98.82 | 99.02 |
| NanoCaller2 |  |  |  |  |  |  |  |
| Genomic Regions | Genomes | Method1 (for ONT reads) |  |  | Method2 (for PacBio reads) |  |  |
|  |  | Precision | Recall | F1 | Precision | Recall | F1 |
| Whole genome | HG002 | 98.26 | 97.79 | 98.03 | 98.09 | 97.37 | 97.73 |
|  | HG003 | 99.03 | 98.72 | 98.88 | 98.95 | 98.40 | 98.68 |
|  | HG004 | 98.97 | 98.66 | 98.82 | 98.90 | 98.29 | 98.60 |
| All difficult regions | HG002 | 96.07 | 94.65 | 95.36 | 95.99 | 94.26 | 95.12 |
|  | HG003 | 96.93 | 96.21 | 96.57 | 96.91 | 95.91 | 96.41 |
|  | HG004 | 96.91 | 96.09 | 96.50 | 96.87 | 95.77 | 96.32 |
| Low Mappability Regions | HG002 | 96.11 | 97.17 | 96.64 | 96.01 | 96.86 | 96.44 |
|  | HG003 | 96.18 | 98.21 | 97.19 | 96.26 | 97.90 | 97.08 |
|  | HG004 | 96.24 | 98.25 | 97.24 | 96.27 | 97.88 | 97.07 |
| Tandem and Homopolymer Repeats | HG002 | 93.88 | 87.39 | 90.52 | 93.92 | 86.76 | 90.20 |
|  | HG003 | 95.56 | 89.51 | 92.44 | 95.35 | 89.25 | 92.20 |
|  | HG004 | 95.25 | 89.01 | 92.03 | 95.17 | 88.64 | 91.79 |
| Segmental Duplications | HG002 | 92.73 | 95.63 | 94.16 | 92.24 | 95.65 | 93.92 |
|  | HG003 | 93.21 | 97.14 | 95.14 | 93.25 | 96.85 | 95.02 |
|  | HG004 | 93.59 | 97.12 | 95.33 | 93.50 | 96.91 | 95.18 |
| MHC | HG002 | 98.90 | 98.25 | 98.58 | 98.88 | 97.95 | 98.42 |
|  | HG003 | 99.43 | 98.98 | 99.21 | 99.47 | 98.69 | 99.08 |
|  | HG004 | 99.43 | 98.94 | 99.19 | 99.38 | 98.83 | 99.11 |

**Table S22.** Performance of NanoCaller whole genome SNP calling for HG002, HG003 and HG004 PacBio CCS and CLR reads using Method1 and Method2 described in Table S18. Shown are the best F1-score achieved by NanoCaller1 SNP model with evaluation on v4.2 benchmark variants.

| PacBio CCS Reads |  |  |  |  |  |  |  |
| --- | --- | --- | --- | --- | --- | --- | --- |
|  | Genomes | Method1 (for ONT reads) |  |  | Method2 (for PacBio reads) |  |  |
|  |  | Precision | Recall | F1 | Precision | Recall | F1 |
| Whole genome | HG002 | 99.27 | 99.53 | 99.40 | 99.31 | 99.60 | 99.45 |
|  | HG003 | 99.33 | 99.56 | 99.45 | 99.36 | 99.63 | 99.49 |
|  | HG004 | 99.33 | 99.57 | 99.45 | 99.37 | 99.63 | 99.50 |
| PacBio CLR Reads |  |  |  |  |  |  |  |
|  | Genomes | Method1 (for ONT reads) |  |  | Method2 (for PacBio reads) |  |  |
|  |  | Precision | Recall | F1 | Precision | Recall | F1 |
| Whole genome | HG002 | 98.68 | 97.73 | 98.20 | 98.55 | 97.98 | 98.27 |
|  | HG003 | 96.15 | 92.47 | 94.28 | 96.46 | 92.28 | 94.33 |
|  | HG004 | 95.80 | 91.15 | 93.42 | 95.95 | 91.14 | 93.49 |

## Effects of allowing WhatsHap to change genotype on SNP calling performance

**Table S23.** Performance of SNP calling by NanoCaller with and without the use of ‘distrust genotypes’ for phasing by WhatsHap. All the other results shown elsewhere besides this table are generated without ‘distrust genotype’ option whiich allows WhatsHap to change genotypes. NanoCaller users can enable ‘distrust genotype’ option by setting ‘enable_whatshap’ flag in NanoCaller run. By default this option is turned off in NanoCaller.

| Oxford Nanopore Reads |  |  |  |  |  |  |  |  |  |  |  |  |
| --- | --- | --- | --- | --- | --- | --- | --- | --- | --- | --- | --- | --- |
|  | NanoCaller1 |  |  |  |  |  | NanoCaller2 |  |  |  |  |  |
|  | Without ‘enable_whatshap’ |  |  | With ‘enable_whatshap’ |  |  | Without ‘enable_whatshap’ |  |  | With ‘enable_whatshap’ |  |  |
|  | Precision | Recall | F1 | Precision | Recall | F1 | Precision | Recall | F1 | Precision | Recall | F1 |
| HG001 | 96.50 | 94.59 | 95.54 | 96.53 | 94.90 | 95.71 | 96.13 | 93.75 | 94.93 | 96.20 | 94.07 | 95.13 |
| HG002 | 98.23 | 97.82 | 98.03 | 98.24 | 97.87 | 98.06 | 98.26 | 97.79 | 98.03 | 98.19 | 97.92 | 98.06 |
| HG003 | 99.00 | 98.76 | 98.88 | 98.93 | 98.80 | 98.87 | 99.03 | 98.72 | 98.88 | 99.05 | 98.68 | 98.87 |
| HG004 | 98.88 | 98.67 | 98.78 | 98.83 | 98.70 | 98.77 | 98.97 | 98.66 | 98.82 | 98.94 | 98.66 | 98.80 |
| HX1 | 92.61 | 88.51 | 90.52 | 92.05 | 88.41 | 90.20 | 93.39 | 88.85 | 91.07 | 92.89 | 88.89 | 90.85 |
| PacBio CCS Reads |  |  |  |  |  |  |  |  |  |  |  |  |
|  | NanoCaller1 |  |  |  |  |  | NanoCaller3 |  |  |  |  |  |
|  | Without ‘enable_whatshap’ |  |  | With ‘enable_whatshap’ |  |  | Without ‘enable_whatshap’ |  |  | With ‘enable_whatshap’ |  |  |
|  | Precision | Recall | F1 | Precision | Recall | F1 | Precision | Recall | F1 | Precision | Recall | F1 |
| HG001 | 96.97 | 99.37 | 98.15 | 97.44 | 99.72 | 98.57 | 96.69 | 99.36 | 98.01 | 97.28 | 99.81 | 98.53 |
| HG002 | 99.31 | 99.60 | 99.45 | 99.56 | 99.79 | 99.68 | 99.16 | 99.56 | 99.36 | 99.52 | 99.83 | 99.68 |
| HG003 | 99.36 | 99.63 | 99.49 | 99.58 | 99.78 | 99.68 | 99.22 | 99.58 | 99.40 | 99.54 | 99.82 | 99.68 |
| HG004 | 99.37 | 99.63 | 99.50 | 99.58 | 99.77 | 99.67 | 99.23 | 99.59 | 99.41 | 99.53 | 99.81 | 99.67 |
| PacBio CLR Reads |  |  |  |  |  |  |  |  |  |  |  |  |
|  | NanoCaller1 |  |  |  |  |  | NanoCaller3 |  |  |  |  |  |
|  | Without ‘enable_whatshap’ |  |  | With ‘enable_whatshap’ |  |  | Without ‘enable_whatshap’ |  |  | With ‘enable_whatshap’ |  |  |
|  | Precision | Recall | F1 | Precision | Recall | F1 | Precision | Recall | F1 | Precision | Recall | F1 |
| HG001 | 97.72 | 90.51 | 93.97 | 98.24 | 90.98 | 94.47 | 96.78 | 93.20 | 94.95 | 97.68 | 94.01 | 95.81 |
| HG002 | 98.55 | 97.98 | 98.27 | 98.78 | 98.25 | 98.51 | 98.11 | 98.46 | 98.29 | 98.17 | 99.09 | 98.63 |
| HG003 | 96.46 | 92.28 | 94.33 | 96.56 | 93.42 | 94.97 | 95.58 | 91.56 | 93.53 | 95.93 | 92.52 | 94.20 |
| HG004 | 95.95 | 91.14 | 93.49 | 96.24 | 92.22 | 94.19 | 94.98 | 90.34 | 92.61 | 95.50 | 91.23 | 93.32 |

**Figure S1.**
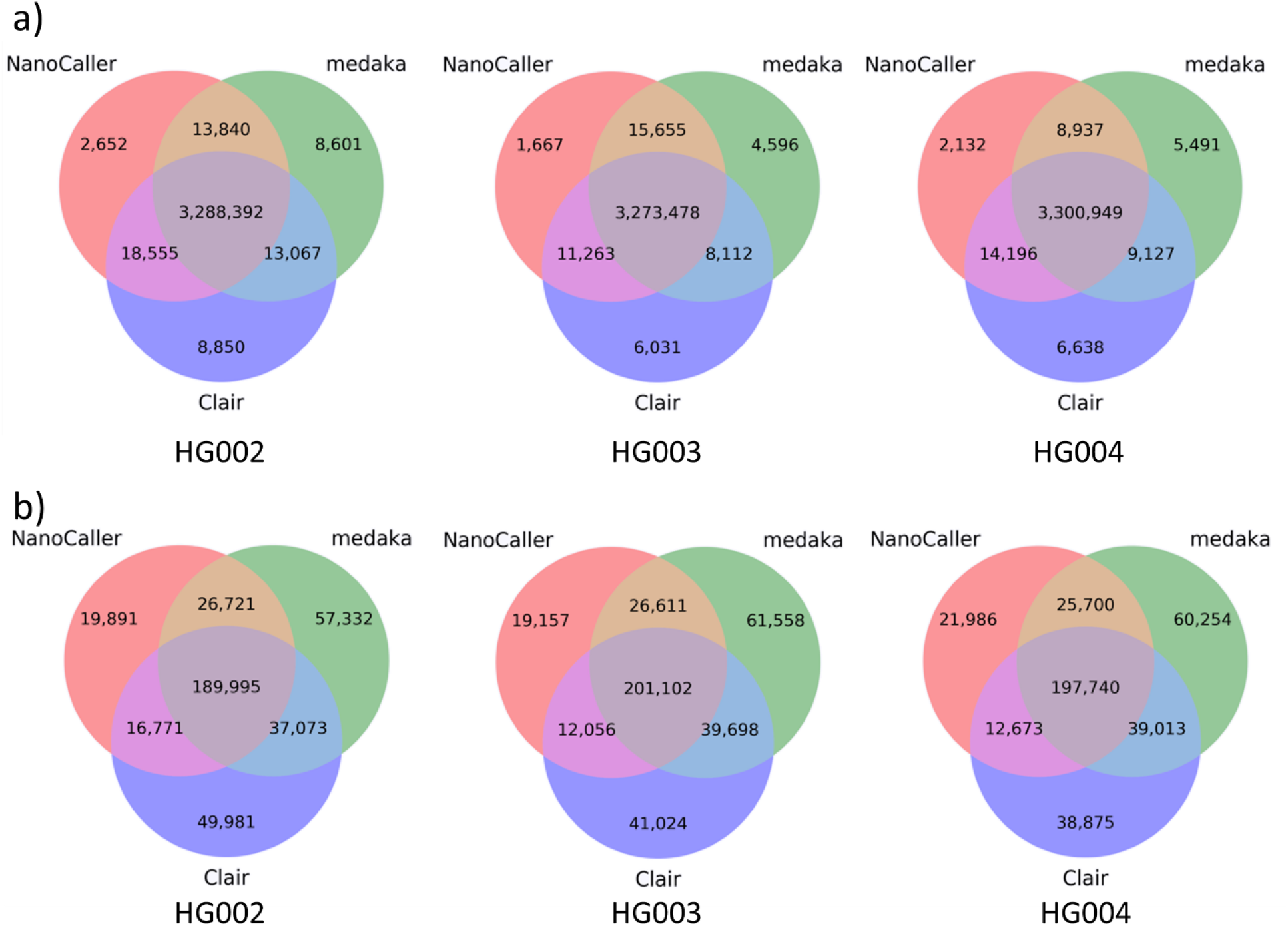
Concordance of ground truth variants correctly predicted by various variant callers on Nanopore reads of the Ashkenazim trio basecalled with Guppy 3.6. Venn diagrams show the overlap of v4.2 ground truth variant calls predicted correctly by NanoCaller, Medaka and Clair. a) SNPs, b) indels. All variants are inside high-confidence regions.

**Figure S2.**
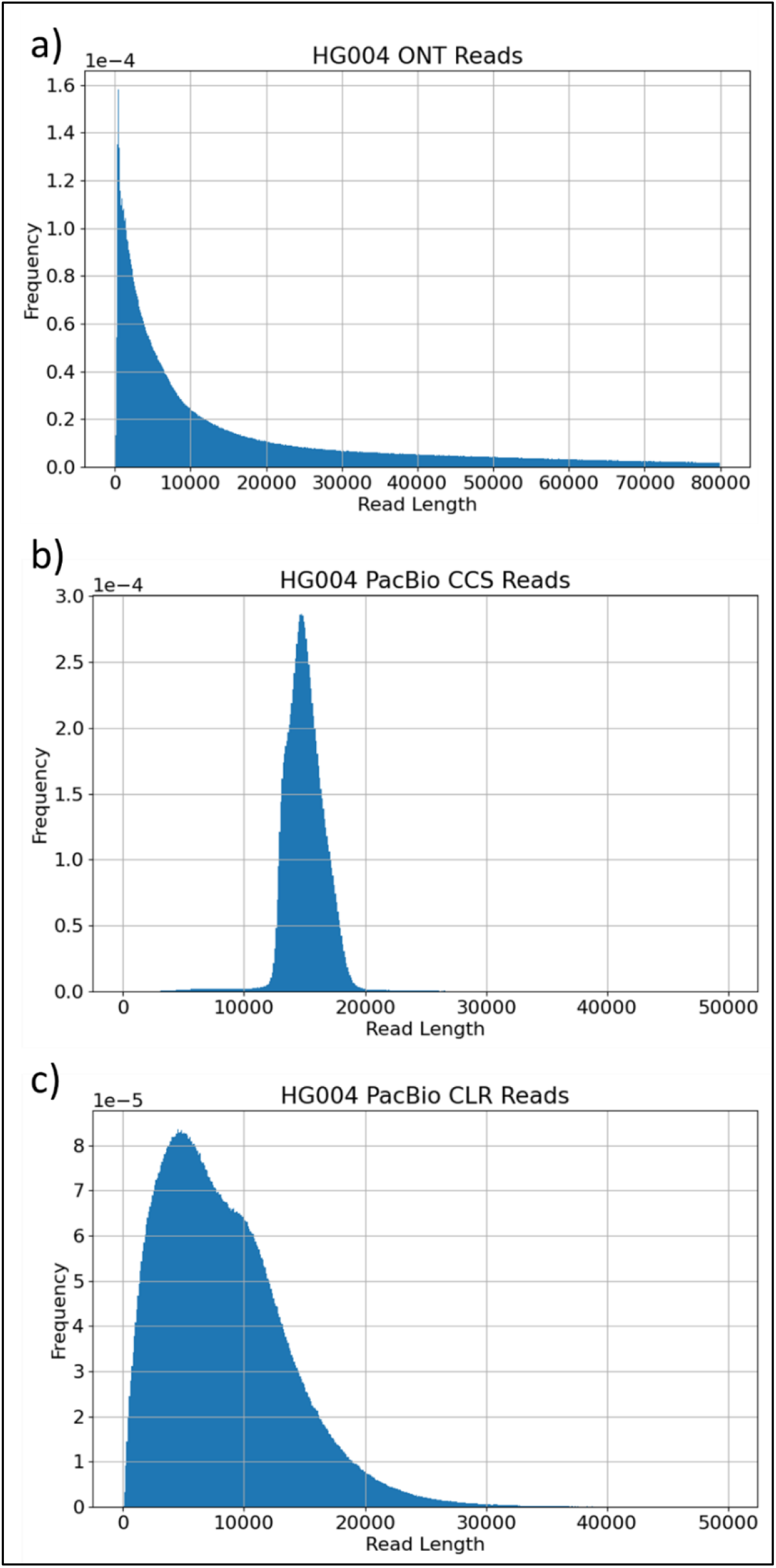
Read length distributions of the following HG004 datasets: 88X ONT reads basecalled by Guppy 3.6, 35X PacBio CCS reads (library size 15kb) and 27X PacBio CLR reads.

**Figure S3.**
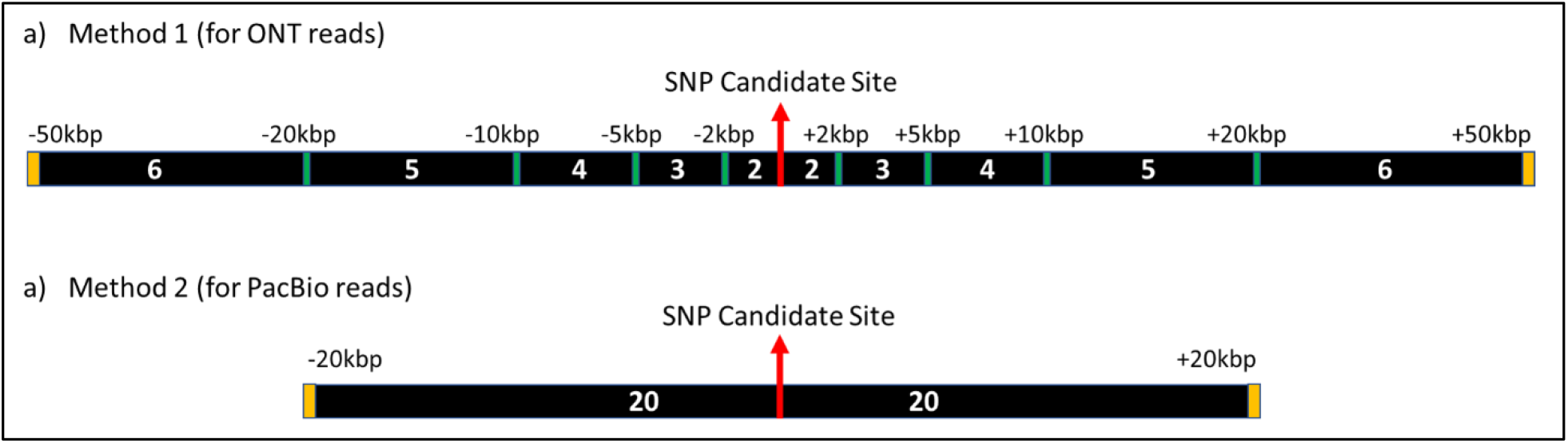
Illustration of number of potentially heterozygous SNP sites chosen in each range for the two methods.

## How to calculate the percentage of high-confidence regions

We download high-confidence regions for HG001-7 from GIAB. For each of them, we calculate the number of bases in high-confidence regions of chromosomes 1-22, and divide it by the total length of chromosomes 1-22 to obtain the percentage of high-confidence regions. According to our calculation, the percentage of high-confidence regions is 81.05% for HG001, 88.44% for HG002, 87.97% for HG003, 87.83% for HG004, 79.67% for HG005, 81.67% for HG006, and 81.59% for HG007. Roughly, high-confidence regions cover 81-88% of human genome. The total length of high-confidence intervals for chr1-22 are calculated using the command below.

Total length of high-confidence regions:

‘cat high_confidence.bed | awk ‘{sum+=$3-$2+1}END{print sum}’’

“high_confidence.bed” is the BED file for GIAB high-confidence intervals.

## The command templates and examples to run NanoCaller and RTG-tools

The commands below can be used to reproduce the results generated by NanoCaller.

For ONT datasets:

‘python NanoCaller_WGS.py -bam HG002.nanopore.bam -ref GRCh38.fa -prefix HG002 -mode both -seq ont -snp_model ONT-HG002_guppy4.2.2_giab-4.2.1 -indel_model HG002_ont_indel -o output -sample HG002 -cpu 16 -min_allele_freq 0.15 -min_nbr_sites 1 -exclude_bed hg38 -mincov 8 -maxcov 160 -nbr_t ’0.4,0.6’ -ins_t 0.4 -del_ t 0.6 -win_size 10 -small_win_size 4’

For PacBio CCS datasets:

‘ python NanoCaller_WGS.py -bam HG002.CCS.bam -ref GRCh38.fa -prefix HG002 -mode both -seq pacbio -snp_model CCS-HG002 -indel_model CCS-HG002 -o output -sample HG002 -cpu 16 -min_allele_freq 0.15 -min_nbr_sites 1 -exclude_bed hg38 -mincov 4 -maxcov 160 -nbr_t ’0.3,0.7’ -ins_t 0.4 -del_t 0.4 -win_size 10 -small_win_size 4 ’

For PacBio CLR datasets:

‘ python NanoCaller_WGS.py -bam HG002.CLR.bam -ref GRCh38.fa -prefix HG002 -mode snps_unphased -seq pacbio -snp_model CLR-HG002 -o output -sample HG002 -cpu 16 -min_allele_freq 0.15 -min_nbr_sites 1 -exclude_bed hg38 -mincov 4 -maxcov 160 -nbr_t ’0.3,0.6’’

Users can replace the reference file (specified by “-ref”), and input BAM file (specify by “-bam”) for different datasets with snp or indel calling or both (specified by “-mode”). The well-trained models for SNP and indel calling can be specified by “-snp_model” and “-indel_model” parameters.

The command template below is used to evaluate predicted VCF against benchmark variant sets.

‘rtg vcfeval -b benchmark.vcf.gz -c variant_calls.vcf.gz -t GRCh38.sdf -e evaluation_region.bed -Z -f

‘QUAL’ -o performance_statistics’

Users can replace the reference .sdf folder specified by ‘-t’, benchmark VCF file specified by ‘-b’, output VCF file by NanoCaller or other tools specified by ‘-c’, and evaluation regions can be specified by *‘-e’*.

.sdf folder for a reference genome using the following command:

‘ rtg format -f fasta GRCh38.fa -o GRCh38.sdf’

## Creating homopolymer and non-homopolymer evaluation regions for indels

First, we download GRCh38_AllHomopolymers_gt6bp_imperfectgt10bp_slop5.bed and GRCh38_SimpleRepeat_homopolymer_4to6_slop5.bed BED files from GIAB genome stratification V2.0 as described on Supplementary Materials Page 5.

Commands to create homopolymer regions:

‘cat GRCh38_AllHomopolymers_gt6bp_imperfectgt10bp_slop5.bed GRCh38_SimpleRepeat_homopolymer_4to6_slop5.bed |bedtools sort > all_homopolymers.bed’

Commands to intersect homopolymer regions with high-confidence regions:

‘bedtools intersect -a high_confidence.bed -b all_homopolymers.bed > high_confidence_homopolymers.bed’

Commands to remove homopolymer regions from high-confidence regions:

‘bedtools subtract -a high_confidence.bed -b GRCh38_SimpleRepeat_homopolymer_4to6_slop5.bed| bedtools subtract -a - -b GRCh38_AllHomopolymers_gt6bp_imperfectgt10bp_slop5.bed > high_confidence_minus_homopolymer_repeats.bed’

## Size distribution of indels in GIAB benchmark variant sets

**Table S24.** Size distribution of indels in high-confidence intervals of GIAB v4.2.1 benchmarks for HG002-4. Each column is for indels in a specific length range.

|  | 2-10bp | 11-20bp | 21-30bp | 31-40bp | 41-50bp | >50bp |
| --- | --- | --- | --- | --- | --- | --- |
| HG002 | 494630 | 23400 | 5382 | 1479 | 559 | 19 |
| HG003 | 477163 | 21169 | 4573 | 1153 | 439 | 3 |
| HG004 | 483180 | 21289 | 4485 | 1143 | 422 | 0 |

## Statistics and performance with old NanoCaller model and datasets

In **Table S26** and Figure S4-Figure S5 below, three NanoCaller SNP models are evaluated: NanoCaller1 (trained on HG001 ONT reads basecalled with Guppy2.3.8), NanoCaller2 (trained on HG002 ONT reads basecalled with Guppy2.3.4), and NanoCaller3 (trained on HG003 PacBio CLR reads). Two NanoCaller indel models are also tested: NanoCaller1 (trained on HG001 ONT reads basecalled with Guppy2.3.8) and NanoCaller3 (trained on HG001 PacBio CCS reads (11kb library size)). Testing datasets include: ONT HG001 reads basecalled by Guppy 2.3.8, ONT HG002-4 reads from precisionFDA challenge basecalled by Guppy 3.6, HX1 reads basecalled with Albacore, HG001 CCS (11kb library size) reads obtained GIAB’s database, and HG002-4 CCS (15kb library size) reads downloaded from precisionFDA truth challenge V2.

**Table S25.**
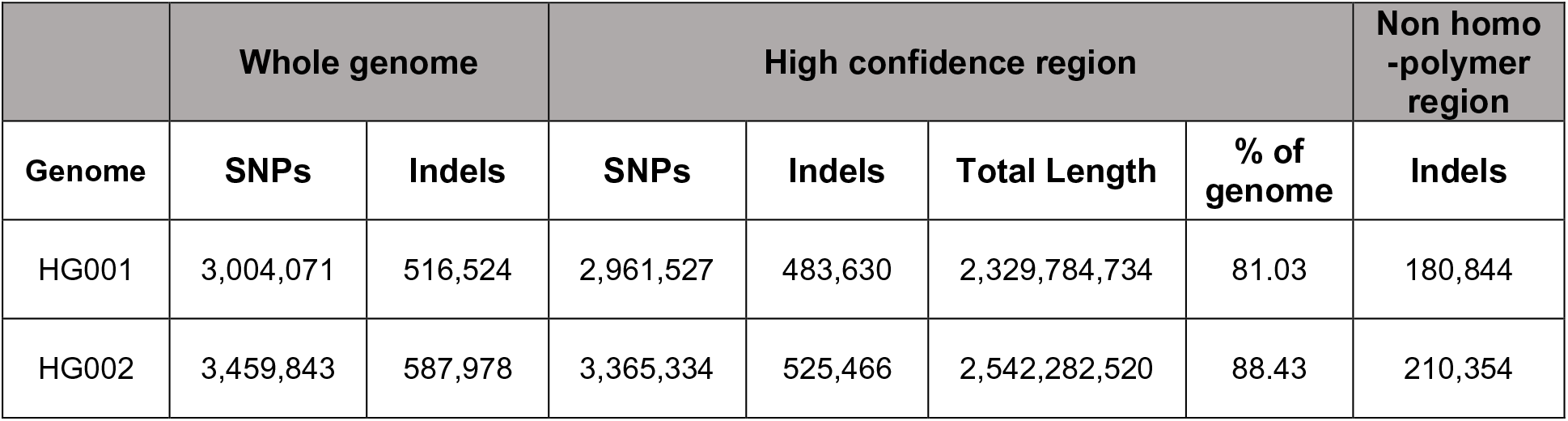
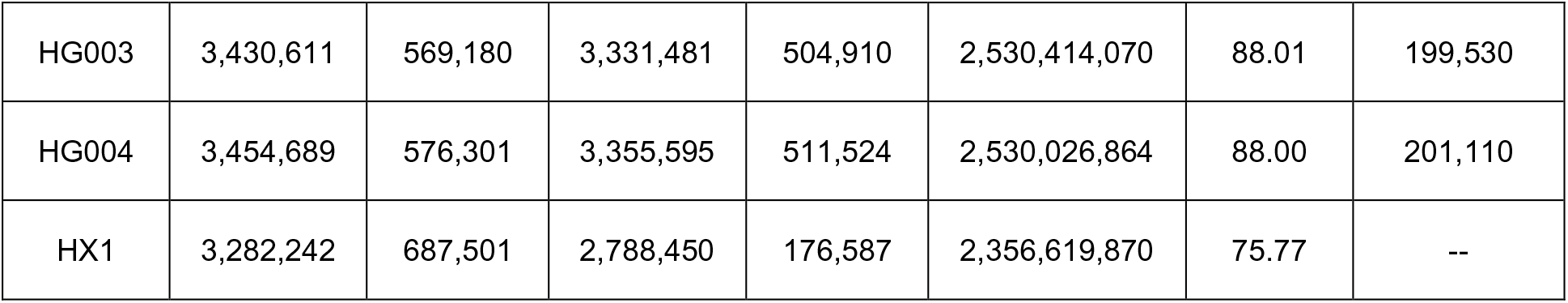
Statistics of benchmark variants in chromosomes 1-22 of each genome aligned to the GRCh38 reference genome. Four genomes with GIAB benchmark variant calls (with v3.3.2 for HG001 and v4.2 for HG002- 4), and statistics within the high confidence regions are also given. For HX1, high confidence regions are created by removing GIAB low complexity regions from the GRCh38 reference genome.

|  | Whole genome |  | High confidence region |  |  |  | Non homo-polymer region |
| --- | --- | --- | --- | --- | --- | --- | --- |
| Genome | SNPs | Indels | SNPs | Indels | Total Length | % of genome | Indels |
| HG001 | 3,004,071 | 516,524 | 2,961,527 | 483,630 | 2,329,784,734 | 81.03 | 180,844 |
| HG002 | 3,459,843 | 587,978 | 3,365,334 | 525,466 | 2,542,282,520 | 88.43 | 210,354 |
| HG003 | 3,430,611 | 569,180 | 3,331,481 | 504,910 | 2,530,414,070 | 88.01 | 199,530 |
| HG004 | 3,454,689 | 576,301 | 3,355,595 | 511,524 | 2,530,026,864 | 88.00 | 201,110 |
| HX1 | 3,282,242 | 687,501 | 2,788,450 | 176,587 | 2,356,619,870 | 75.77 | -- |

**Table S26.**
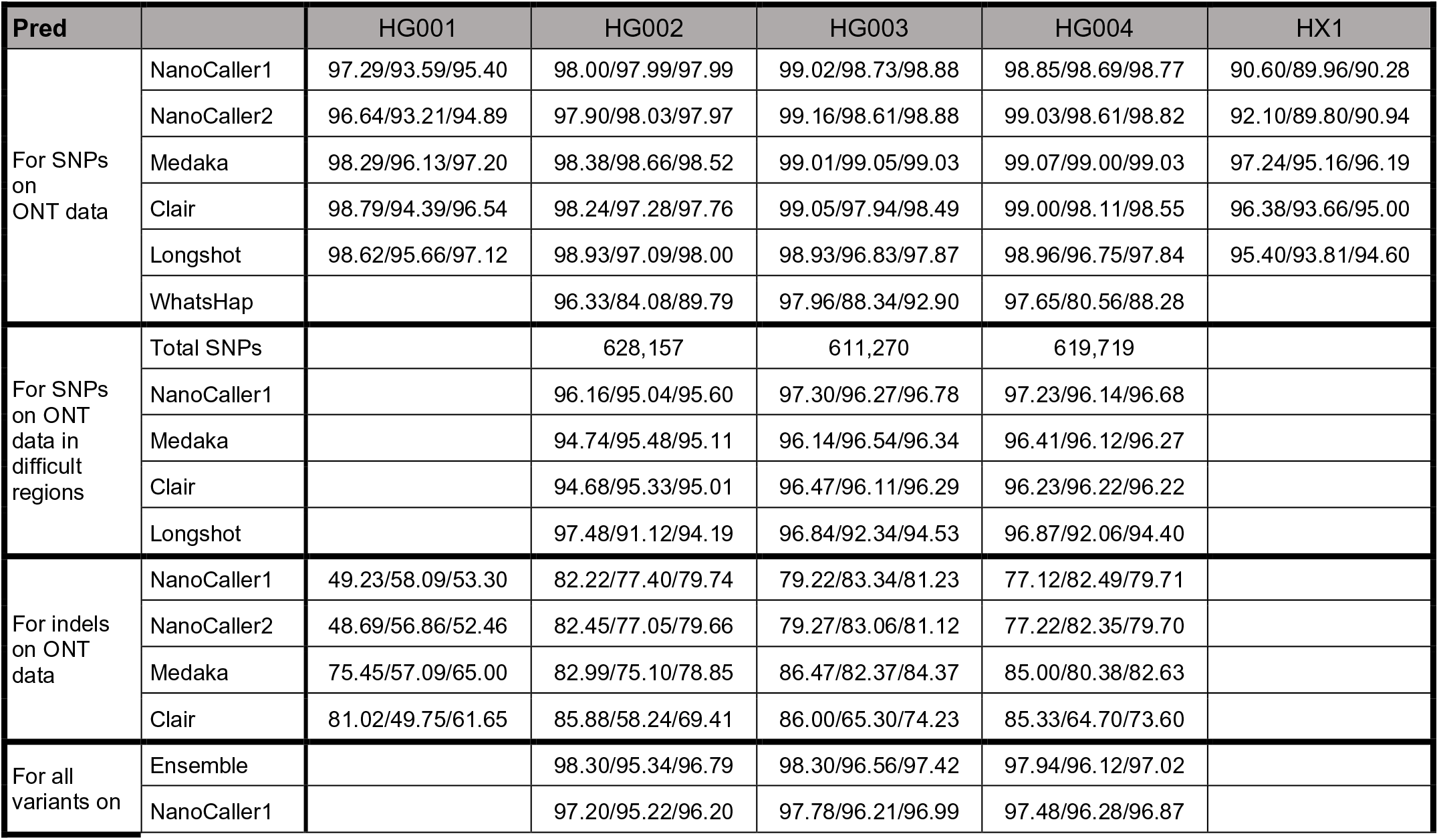
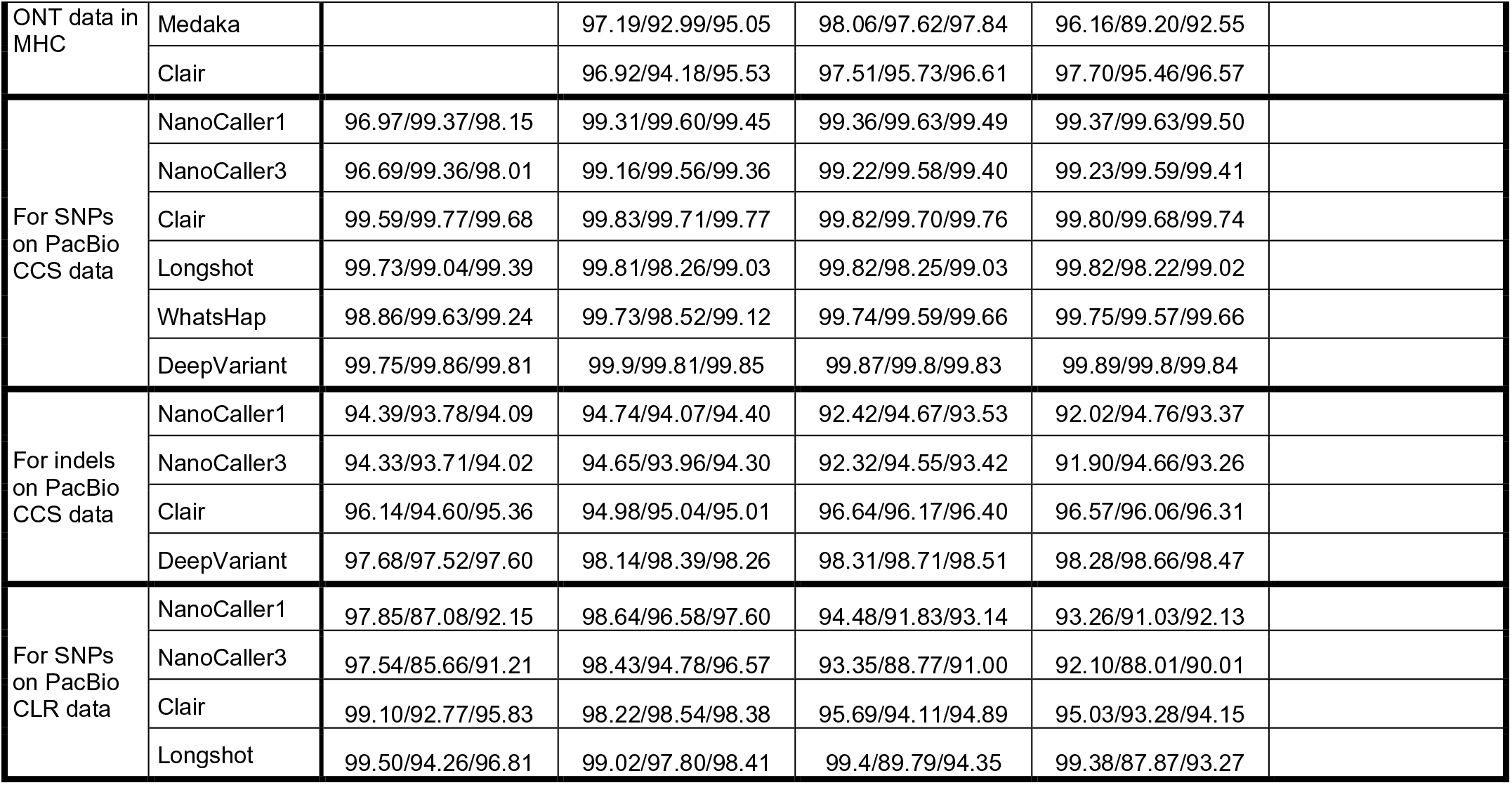
Performances (Precision/Recall/F1 percentages) of SNP and indel predictions by NanoCaller1, NanoCaller2 and NanoCaller3 on ONT and PacBio (CCS and CLR) data and on difficult-to-map genomic regions along with the performance of existing variant callers. These evaluation is based on v3.3.2 benchmark variants for HG001 and v4.2 benchmark variants for the Ashkenazim trio (HG002, HG003, HG004).

**Figure S4.**
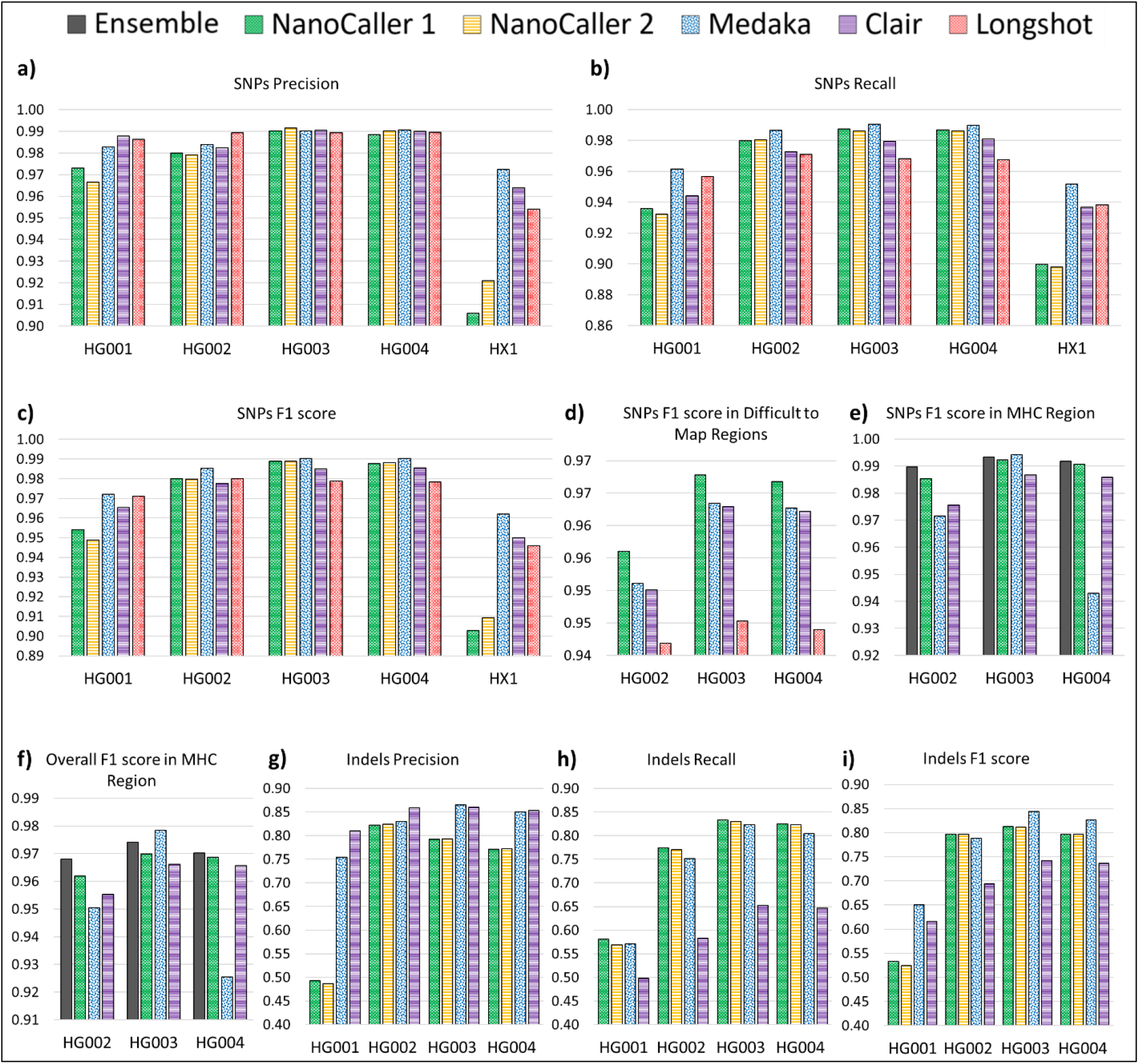
Performance of NanoCaller and state-of-the-art variant callers on five whole-genome Oxford Nanopore sequencing data sets. The performance of SNP predictions on ONT reads: a) precision, b) recall, c) F1 score. d) The performance of SNP predictions on HG002 (ONT), HG003 (ONT) and HG004 (ONT) in difficult-to-map genomic regions. The performance of variant predictions on HG002 (ONT), HG003 (ONT) and HG004 (ONT) in Major Histocompatibility Complex regions: e) SNPs, f) overall variants. The performance of indel predictions on ONT reads in non-homopolymer regions: g) precision, h) recall, i) F1 score. For HX1, the variants, which were called on high-coverage short-read data are used as benchmark with complement of difficult-to-map genomic regions used in d) as high-confidence regions. Benchmark variants v3.3.2 for HG001 and v4.2 for the Ashkenazim trio (HG002, HG003, HG004) are used for evaluation.

**Figure S5.**
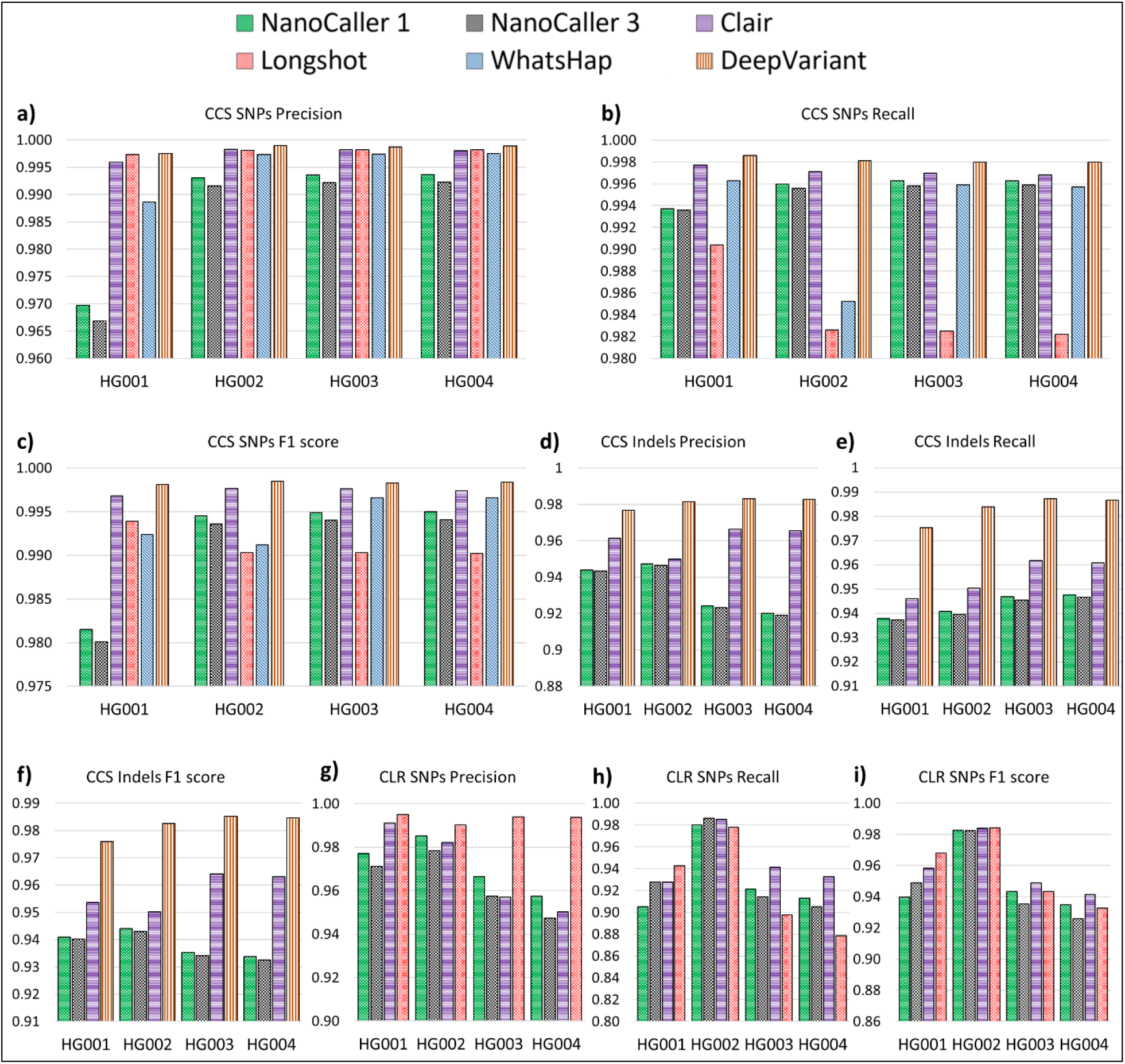
Performance of NanoCaller and state-of-the-art variant callers on PacBio sequencing data sets. The performance of SNP predictions on PacBio CCS reads: a) precision, b) recall, c) F1 score. The performance of indel predictions on PacBio CCS reads: d) precision, e) recall, f) F1 score. The performance of SNP predictions on PacBio CLR reads: g) precision, h) recall, i) F1 score. Benchmark variants v3.3.2 for HG001 and v4.2 for the Ashkenazim trio (HG002, HG003, HG004) are used for evaluation.

**Figure S6.**
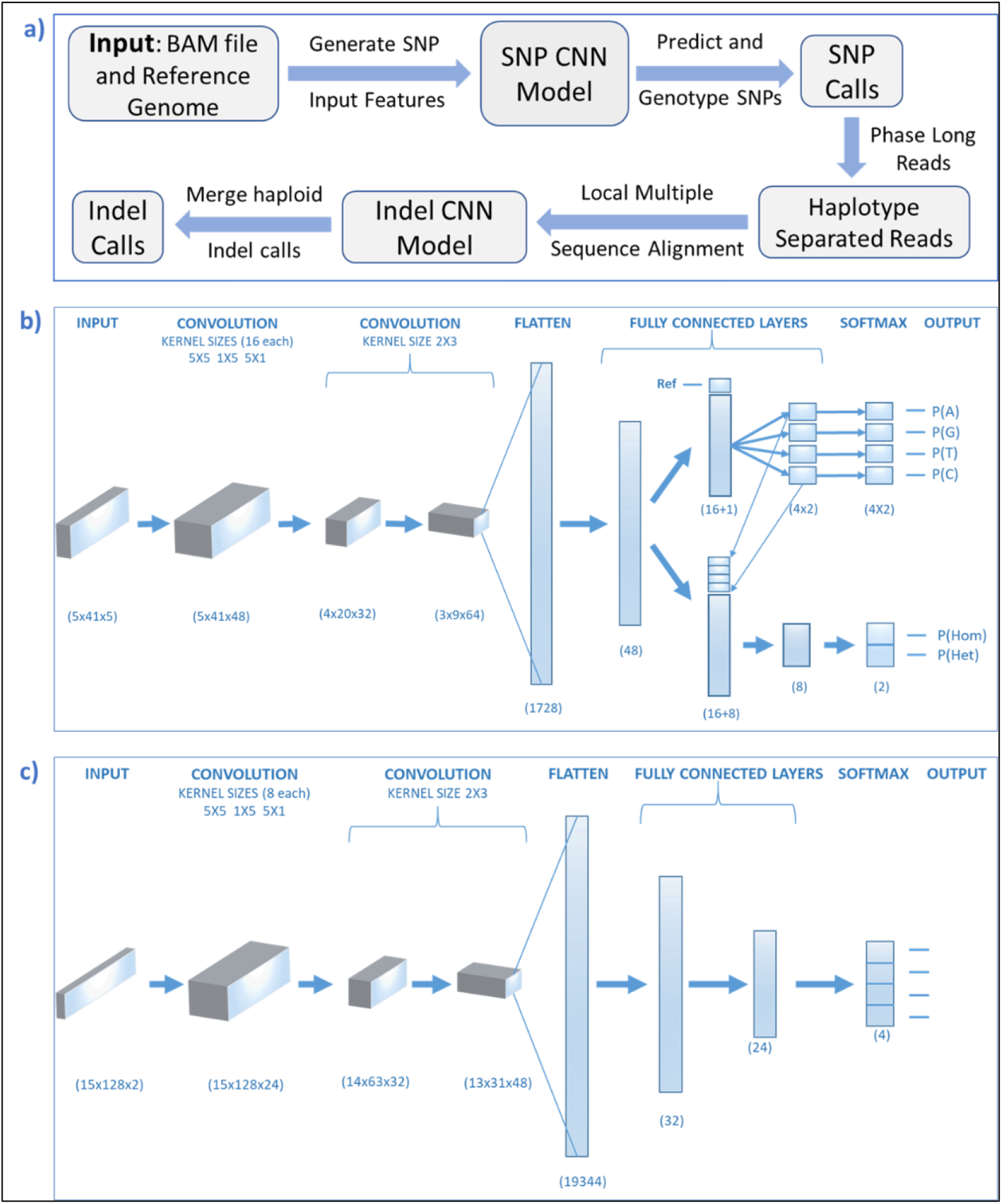
The deep-learning framework for SNP and indel calling. a) The overall workflow of NanoCaller. b) An illustration of the convolutional neural network model for SNP calling; c) An illustration of the convolutional neural network model for indel calling. In both models, first convolutional layer uses 3 kernels of sizes 1x5, 5x1 and 5x5, whereas the second and third convolutional layers use kernels of size 2x3. Output of third convolutional layer is flattened and fed to a fully connected layer followed by a hidden layer with 48 nodes with a 50% dropped rate. In b), output of the hidden layer is split into two independent pathways: one for calculating probabilities of each base and the other for calculating zygosity probabilities. Zygosity probability is only used in the training process. In c), output of first fully connected layer is fed into two fully connected hidden layers to produce probabilities of four possible cases of zygosities.

**Figure S7.**
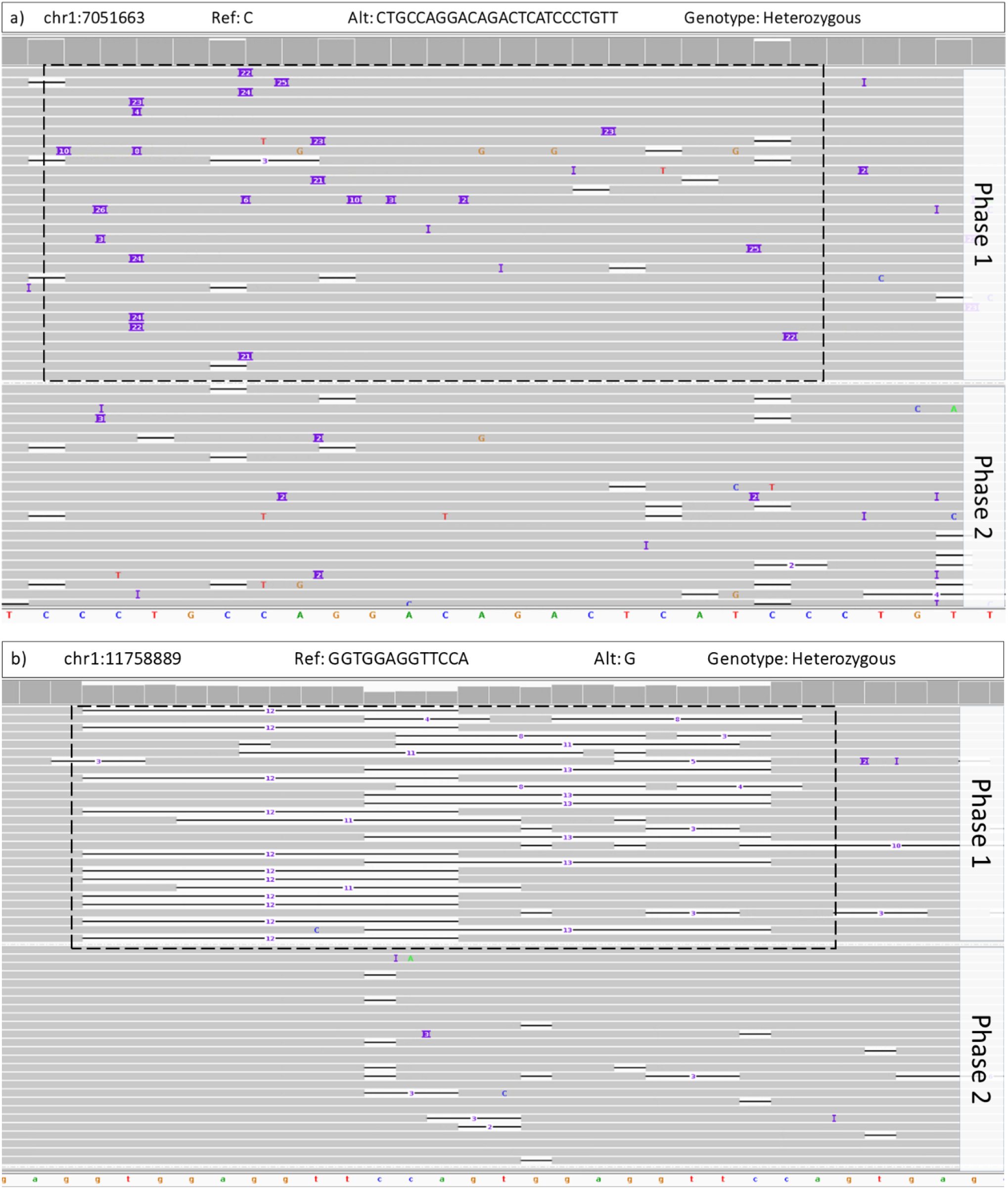
Two indels from HG002 GIAB benchmark v4.2.1 identified correctly by NanoCaller but missed by Clair and Medaka. Both indels lie in tandem repeats, therefore with high discordance among ONT reads for the locus of the indel (shown in black rectangle). NanoCaller uses a sliding window to detect such indels hat do not have high allele frequency at any single reference genome locus. a) An insertion; b) An deletion.

